# Minimal essential requirements for neural tube self-organisation

**DOI:** 10.64898/2026.06.06.730555

**Authors:** Hannah T. Stuart, Elena Costantini, Jingkui Wang, Teresa Krammer, M. Joaquina Delas, Manuela Melchionda, Jake Cornwall-Scoones, Gerald Schmauss, Thomas Lendl, Keisuke Ishihara, David Rand, Meritxell Sáez, Elly M. Tanaka, James Briscoe

## Abstract

The reliable generation of diverse cell types in precise proportions is essential for the formation of functional tissues during embryonic development. Three-dimensional organoid models derived from pluripotent stem cells (PSCs) provide powerful systems for identifying principles governing tissue self-organisation. Neural tube organoids (NTOs), initiated from single PSCs with a pulse of retinoic acid (RA), self-organise into structures containing floorplate cells that secrete SHH morphogen to pattern adjacent neural tissue. Yet, how the initial cellular diversity arises and how appropriate cell-type proportions are allocated has remained unclear. Here, using time-resolved single-cell transcriptomics, quantitative immunofluorescence, and dynamical systems modelling we show that RA triggers a transient co-expression state of the transcription factors PAX6 and FOXA2 from which cells asynchronously resolve into two opposing fates: PAX6⁺ neural precursors and FOXA2⁺ floorplate precursors. PAX6 and FOXA2 are both necessary and sufficient to reconstitute self-organisation, establishing these transcription factors as key determinants of emergent tissue pattern. Rather than operating as a simple feed-forward system in which cells are guided solely by RA, feedback between the alternative cell fates, mediated by BMP signalling from floorplate precursors, determines cell type proportions and ensures reproducible cell-type diversity in each NTO. This dual expression state was also identified in mouse embryos, demonstrating how *in vitro* models inform *in vivo* biology. These findings establish a general design strategy – symmetry breaking through opposing fate determinants coupled to proportioning via signal feedback control – that may operate broadly across developmental contexts to generate tissues with predictable cellular compositions.

## Introduction

The formation of complex multicellular tissues depends on the precisely choreographed differentiation of cells to ensure appropriate cell types emerge at the right time, place and proportions. Tissue patterning can be instructed by directional cues, such as asymmetric maternal factors or morphogens secreted by embryonic organisers (Gregor et al., 2007; Kicheva and Briscoe, 2023; Yan et al., 2018). Alternatively, tissue pattern can arise autonomously by self-organisation, emerging from local interactions between cells (Abitua et al., 2024; Kirillova et al., 2018; Wennekamp et al., 2013). Many cases lie between the two ends of this spectrum – instructed versus self-organised (Brückner and Tkačik, 2025) – motivating framing development as guided self-organisation (Green and Sharpe, 2015; Morales et al., 2021). Patterning involves bi-directional causality: while tissue-level patterns emerge from cellular-scale events, tissue-level properties can feedback to influence cellular decisions. This underpins the regulative nature of vertebrate development, i.e. the ability to sense and respond to perturbations, correcting course to compensate for altered conditions (De Robertis, 2009; Snow and Tam, 1979; Tarkowski, 1961).

In mammals, the contribution of self-organisation to development and the mechanisms governing regulative decisions are challenging to study after the blastocyst stage, due to the embryo’s complexity and *in utero* location. *In vitro*, mammalian stem cells reveal emergent collective properties, including the spontaneous generation of differentiation patterns in organoids and embryoids (Eiraku et al., 2011; Lancaster et al., 2013; Sato et al., 2009; Van Den Brink et al., 2014). *In vitro* models offer the opportunity for mechanistic study under defined conditions, informing questions to ask of the complex embryo and revealing design principles to harness self-organisation for tissue engineering and regeneration.

Here, we exploit the manipulability and modularity of embryonic stem cell (ESC) systems to ask how cells diversify from a clonal population and how the correct proportions of cell types are generated for emergent self-organisation. We chose mouse neural tube organoid (NTO) differentiation to ask these questions (Meinhardt et al., 2014) (Fig. 1A). NTOs combine the phenomena of symmetry breaking, organiser self-organisation and morphogen patterning. By starting from a single ESC, NTOs are unconfounded by pre-existing heterogeneity. NTOs are efficient and reproducible, accessible to genetic and chemical perturbations, and tractable for quantitative whole-tissue analyses.

**Figure 1:**
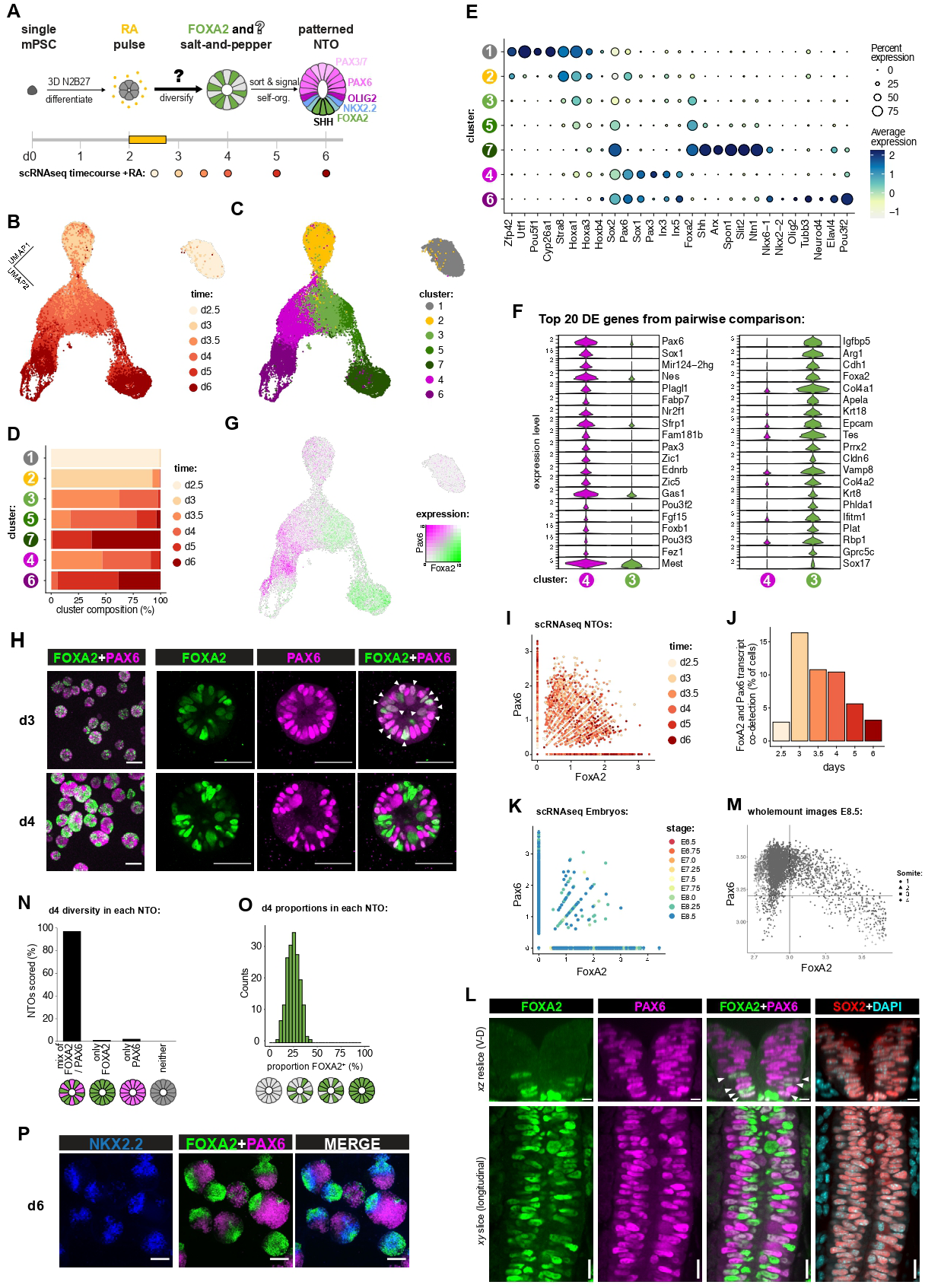
RA triggers diversification of cell states to yield a mix of pFP and eNP in each NTO. **A:** Schematic of NTO formation, from a single cell to a patterned tissue. On day (d)2, a pulse of retinoic acid (RA) is added for 18 hours (h). NTOs were harvested for scRNA-seq at the indicated timepoints. **B-C:** UMAP embedding of scRNA-seq timecourse data for RA-treated NTOs from d2.5-6 (B) with 7 distinct clusters identified in Seurat (C). **D:** Percentage of cells from each timepoint contributing to each cluster in (C). **E:** Dot plot showing selected marker genes for each cluster. The circle size represents the fraction of cells expressing the gene and colour represents the average expression. **F:** Violin plots for the top 20 differentially expressed (DE) genes between clusters 3 and 4. **G:** Co-expression feature plot for *FoxA2* and *Pax6* on the same UMAP embedding as (B-C). **H:** Immunofluorescence for FOXA2 (green) and PAX6 (magenta) at d3 and d4 on RA-treated NTOs. Left: representative overview, maximum intensity *z*-projection acquired with 10x objective, scale bar 100μm. Right: representative NTO, single confocal *z* slice acquired with 40x objective, scale bar 50μm. White arrows indicate co-expressing cells. **I-J:** Scatterplot of *FoxA2* versus *Pax6* expression in scRNA-seq data from d2.5-6 for RA-treated NTOs (I), and the percentage of cells at each timepoint in which both transcripts are detected (J). **K:** Scatterplot of *FoxA2* versus *Pax6* expression, from re-analysis of published scRNA-seq atlas of developing mouse embryos from E6.5-E8.5, subsetted for cells annotated as neurectoderm, brain or spinal cord. **L:** Immunofluorescence for FOXA2 (green), PAX6 (magenta), SOX2 (orange) and DAPI (cyan) on a mouse embryo harvested at E8.5 with developmental stage of 7-somites. Representative *xz-*reslice at the level of somite 3 (hindbrain), from wholemount confocal acquisition with 30x objective and 0.41μm *z*-step, scale bar 50μm. **M:** Quantification of FOXA2 and PAX6 expression level in every nucleus of the neural tube at the levels of somite 1-4 (hindbrain) of 7-somite stage embryos (n=2, see also Fig. S1J), from wholemount confocal acquisition with 30x objective and 0.41μm *z*-step. Nuclei were segmented on SOX2 staining using CellPose-SAM, with manual curation to remove SOX2^+^ nuclei from other tissues than the neural tube. See Table S2 for further scoring of additional embryos (n=20). **N:** Quantification of NTOs that express FOXA2 and/or PAX6 at d4. A total of n=1392 NTOs were manually scored, assessing presence/absence of markers. **O:** Proportion of each NTO that is FOXA2^+^ at d4. A total of n=176 NTOs were quantified, with a mean of 27% and standard deviation of 7%. **P:** Immunofluorescence for FOXA2 (green), PAX6 (magenta) and NKX2.2 (blue) at d6 on RA-treated NTOs. Representative maximum intensity *z*-projection acquired with 10x objective, scale bar 100μm. See also Figure S1.

To form NTOs, a pulse of retinoic acid (RA) is applied early during 3D neural differentiation. This triggers self-organisation of a floorplate (FP) organiser at an emergent ventral pole. SHH morphogen secreted from the FP drives ventral-to-dorsal neural progenitor (NP) patterning across a pseudostratified epithelium with a single apical lumen, yielding an *in vivo*-like tissue pattern and architecture (Meinhardt et al., 2014) (Fig. 1A). Previously, we showed that FP self-organisation occurs through a combination of cell sorting to agglomerate initially scattered FOXA2^+^ FP precursors, and long-range BMP-mediated signalling competition between FOXA2^+^ clusters to establish a ‘winner’ (Krammer et al., 2024). But this left unanswered how the initial heterogeneity and appropriate cell type proportions are established from a clonal population of cells in a uniform environment. Addressing these questions places NTOs as a powerful quantitative platform to dissect fundamental mechanisms governing the generation of the cellular diversity in the context of emergent tissue patterning.

Here, we combine quantitative time-resolved analyses with dynamical systems modelling to uncover the mechanisms governing cellular diversification and proportioning in NTOs. We validate these findings with systematic functional perturbations in clonal and chimeric systems. We demonstrate that successful self-organisation requires proportional generation of two opposing cell states: FOXA2^+^ floorplate precursors (∼25%) and PAX6^+^ neural precursors (∼75%). A dynamical landscape model that predicts experimental outcomes reveals that cell fate diversification and proportions arise through regulative feedback rather than cell-autonomous decisions. This led us to uncover that early differentiating floorplate precursors signal via BMP to constrain their own proportion, ensuring robust tissue-level outcomes. Remarkably, the entire RA-dependent symmetry-breaking process can be bypassed synthetically by inducing transient FOXA2 and PAX6 co-expression, identifying this dual transcription factor state as the minimal molecular requirement for initiating NTO self-organisation. The observation of analogous co-expression signatures during early floorplate formation in mouse embryos confirms the developmental relevance of these organoid-derived principles. Together, this work deconstructs and reconstructs key principles and minimal requirements to create a self-organising tissue from a single cell, highlighting regulative feedback as a fundamental mechanism for achieving reproducible cellular diversity in mammalian neural development.

## Results

### A pulse of RA triggers diversification to yield a mix of floorplate and neural precursors

To profile the cellular diversity induced by RA treatment in NTOs, we performed a time-resolved scRNAseq timecourse from day (d)2-6, sampling every 12-24 hours (h) (Fig. 1A-B, S1A). After RA addition at d2, we observed a substantial change in the transcriptome at d2.5, followed by another transition at d3 after RA withdrawal (Fig. 1B, S1A-B). Subsequently, RA-treated samples diverged into two major branches from d3.5-d6, in contrast to control untreated samples which followed a single neural differentiation trajectory (Fig. S1A, S1C-E). To identify the discrete cell states induced by RA, we performed clustering and differential gene expression analyses (Fig. 1C-D, S1F). We complemented this with feature selection to find genes associated with the continuous branching in RA-treated expression space (Fig. S1G), integrated the lists (Table S1), and focussed on genes with RA-specific dynamics (Fig. S1B-C, S1H).

Together with an RA response signature (*Cyp26a1*, *Stra8*, *Hox*), we found accelerated pluripotency downregulation (*Oct4/Pou5f1*, *Utf1*) and *Pax6* upregulation from d2-3 during (cluster 1) and after (cluster 2) the RA pulse (Fig. 1E, S1B-C, S1F, S1H-I). RA-accelerated *Pax6* kinetics were not shared by all neurectoderm markers: *Sox1* was upregulated by d4 at a similar pace with or without RA, resulting in an inverted order of *Pax6* then *Sox1* induction with RA (Fig. S1C, S1I). *FoxA2* induction occurred specifically in RA-treated samples, commencing after RA withdrawal (cluster 2) contemporaneous with a transient drop in *Sox2* expression at the ‘head’ of the two branches (Fig. 1E, S1B-C, S1H-I). As the branches diverge at d3.5–4, clusters 3 and 4 represent distinct cell states (Fig. 1C), with *FoxA2* versus *Pax6* and *Sox1* among the top differentially expressed genes, respectively (Fig. 1F-G). In one branch, clusters 5 then 7 capture progressive FP differentiation downstream of cluster 3, culminating in mature FP marker expression (*FoxA2*, *Shh*, *Arx*) (Fig. 1E, S1C, S1F) consistent with FOXA2 induction prior to and independently of SHH (Krammer et al., 2024). In the other branch, cluster 6 comprises neurons (*Tubb3*) and *Sox1*^+^ neural progenitors spanning the ventral-dorsal axis (*Nkx2.2*, *Olig2*, *Pax6*, *Pax3*) (Fig. 1E, S1C, S1F), indicative of successful endpoint NTO patterning (Meinhardt et al., 2014). Together, these data indicate that the two post-RA branches correspond to FP versus neural differentiation, and that the divergence of FP precursors (pFP, cluster 3) from early neural precursors (eNP, cluster 4) represents the major symmetry-breaking event following the post-RA state (cluster 2) (Fig. 1C).

We performed immunofluorescence for FOXA2 and PAX6 protein in RA-treated NTOs. At d3, FOXA2 and PAX6 were widely expressed, with cell-to-cell variability in levels and co-expression (Fig. 1H). We also detected transcriptional co-expression (Fig. 1G, 1I-J). FOXA2 and PAX6 co-expression was surprising given their association with distinct domains in the neural tube. We re-examined published mouse embryo scRNAseq data (Pijuan-Sala et al., 2019) and identified cells co-expressing *FoxA2* and *Pax6* (Fig. 1K). We confirmed *in vivo* co-expression of FOXA2 and PAX6 proteins in the ventral neural plate as NT closure starts at E8.5 (Fig. 1L-M, S1J). This corroborated the existence of FOXA2 and PAX6 co-expression and the *in vivo* relevance of this organoid-identified expression state.

By d4 in NTOs, FOXA2 and PAX6 expression was mostly mutually exclusive (Fig. 1H), consistent with their differential expression in scRNAseq clusters 3 and 4 (Fig. 1F). FOXA2 or PAX6 positive cells were scattered in a salt-and-pepper manner (Fig. 1H), with positive cells for each factor observed in 97% of d4 NTOs (Fig. 1N). Quantification of proportions within each NTO showed an average composition of 27% FOXA2^+^ with a standard deviation of 5% between NTOs at d4 (Fig. 1O), indicating symmetry breaking that yields characteristic proportions within each clonal population. By d6, FOXA2 and PAX6 expression occupied spatially distinct domains with SHH-driven patterning of ventral neural progenitors in between (Fig. 1P).

Together, these data demonstrate that a pulse of RA induces rapid and substantial shifts in cell states, culminating by d4 in diverging differentiation trajectories that produce spatially intermingled PAX6⁺ eNPs and FOXA2⁺ pFPs. This diversity generation is robust with both cell states observed in each clonal NTO in characteristic proportions.

### FOXA2 and PAX6 are required for NTO self-organisation

To test whether PAX6 and FOXA2 are functional drivers of NTO self-organisation downstream of RA, we performed CRISPR-mediated knockout of each gene. Without RA, *FoxA2^−/−^* or *Pax6^−/−^* NTOs underwent neural differentiation with normal SOX1 induction kinetics (Fig. S2A). With RA treatment, at d6 *FoxA2^−/−^* NTOs lacked FP (SHH**^−^**) and did not induce ventral neural progenitor patterning (NKX2.2**^−^**, OLIG2**^−^**) (Fig. 2A), indicating that FOXA2 is essential for FP formation in NTOs. Conversely, RA-treated *Pax6^−/−^* NTOs comprised almost entirely FP (FOXA2^+^, SHH^+^) and ventral-most p3 (NKX2.2^+^) neural progenitors at d6, but lacked OLIG2 (Fig. 2A). Thus, FOXA2 and PAX6 are individually essential for proper NTO patterning downstream of RA, and their knockouts fail with opposite phenotypes, forming no FP or too much FP respectively. We quantified the proportion of cells expressing FOXA2 versus PAX6 at the d4 salt-and-pepper stage in wildtype (*WT*) or knockout NTOs (Fig. 2B). In *FoxA2^−/−^* NTOs, 90% of cells were PAX6^+^ compared to only 65% PAX6^+^ in WT. In *Pax6^−/−^*NTOs, 60% of cells were FOXA2^+^ compared to only 25% in WT. This suggests that FOXA2 and PAX6 normally antagonise each other’s expression, and perturbation of either alters cell fate proportions at d4.

**Figure 2:**
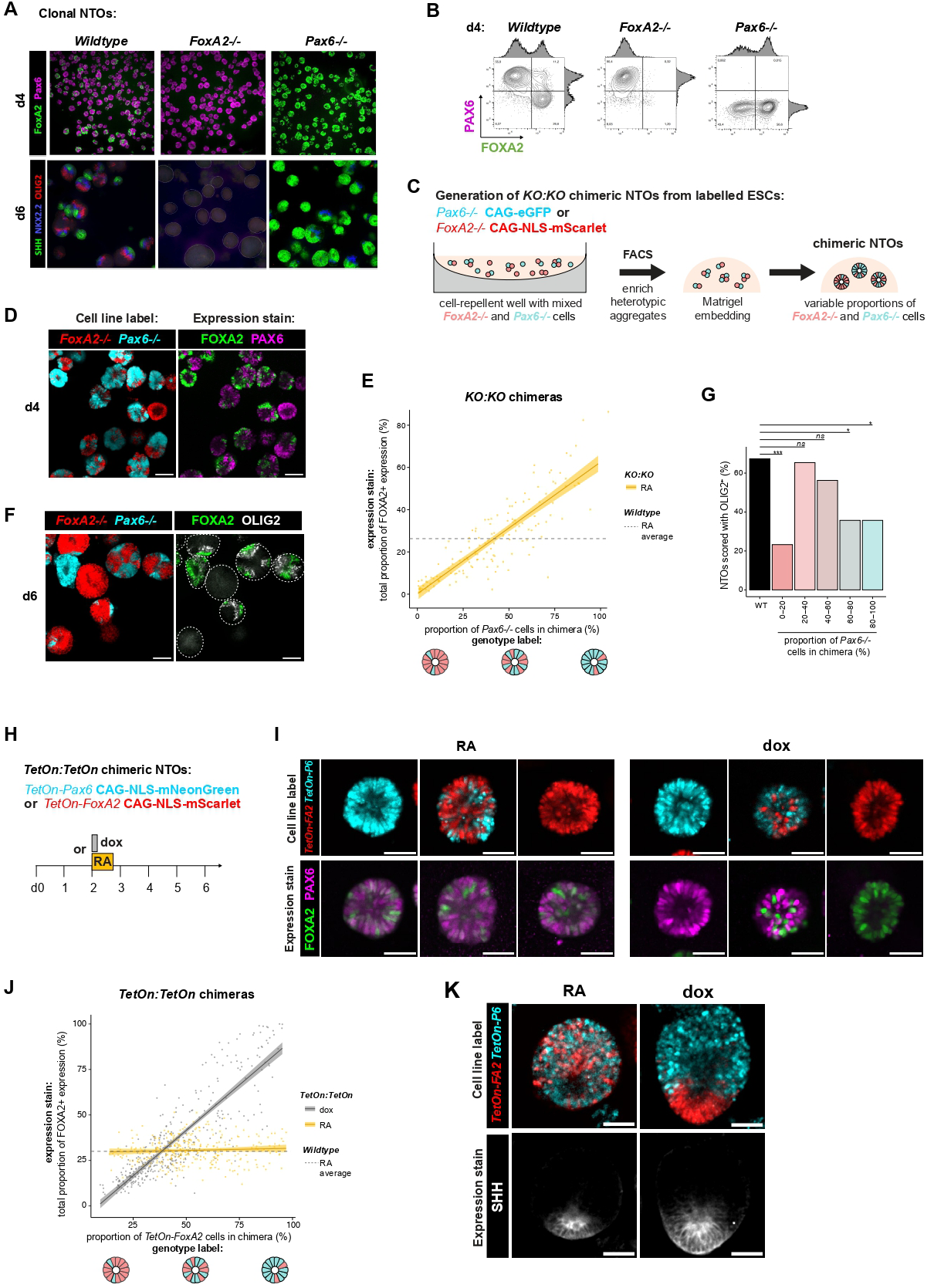
A mix of pFP and eNP is necessary and sufficient for NTO self-organisation. **A:** Wildtype, *FoxA2^−/−^* and *Pax6^−/−^* NTOs were seeded from single ESCs and treated with an RA pulse at d2. Immunofluorescence is shown at d4 for FOXA2 (green) and PAX6 (magenta) as representative maximum intensity *z*-projection, and at d6 for SHH (green), NKX2.2 (blue) and OLIG2 as (red) as representative single confocal *z* slice. **B:** Flow cytometry analysis of FOXA2 and PAX6 immunostaining at d4 on RA-treated NTOs. **C:** Workflow for the generation of *KO:KO* chimeric NTOs from heterotypic ESC aggregates. **D,F:** Immunofluorescence on RA-treated *KO:KO* chimeric NTOs. *FoxA2^−/−^* cells can be traced by their constitutive CAG-NLS-mScarlet label (red), and *Pax6^−/−^* cells by their constitutive CAG-eGFP label (cyan). FOXA2 (green) and PAX6 (magenta) expression are stained at d4 (D). FOXA2 (green) and OLIG2 (grey) expression are stained at d6 (F). Representative single confocal *z* slice, scale bar 100μm. **E:** Percentage of FOXA2^+^ cells in each *KO:KO* chimeric NTO (n = 145) at d4, as a function of the chimeric composition, quantified from 3D images of wholemount immunofluorescence. **G:** Quantification of NTOs that express OLIG2 at d6 (presence/absence of marker) for *KO:KO* chimeras compared to wildtype (n=341). *KO:KO* chimeras were binned according to their genotype composition. *p*-val are indicated on the plot as *** < 10^-9, * < 0.05, or not significant (ns) > 0.2. **H:** Generation of *TetOn:TetOn c*himeric NTOs from heterotypic ESC aggregates. **I,K:** Immunofluorescence of *TetOn:TetOn* chimeric NTOs treated on d2 with a pulse of RA or dox. Whether or not they are dox-induced, cells containing the *TetOn-Pax6* transgene can be traced by their constitutive CAG-NLS-mNeonGreen label (cyan), and cells containing the *TetOn-FoxA2:mCherry* transgene can be traced by their constitutive CAG-NLS-mScarlet label (red). FOXA2 (green) and PAX6 (magenta) expression are stained on d3 after RA or dox pulse (I). SHH (grey) is stained at d6 (K). Representative single confocal *z* slice, scale bar 50μm. **J:** Percentage of FOXA2^+^ cells in each *TetOn:TetOn* chimeric NTO at d4, as a function of chimeric composition, quantified from 3D images of wholemount immunofluorescence for RA or dox treatments (n=1375). See also Figure S2.

### A mix of FOXA2^+^ pFPs and PAX6^+^ eNPs is necessary and sufficient for self-organisation

Since *FoxA2* and *Pax6* are individually required for proper NTO patterning but not for each other’s expression (Fig. 2A–B), we hypothesised that combining *FoxA2⁻^/^⁻* and *Pax6⁻^/^⁻* cells in chimeric *‘KO:KO’* NTOs would restore a mixture of FP and neural cells in more appropriate proportions. To generate chimeric NTOs despite their usually clonal origin, we labelled *FoxA2⁻^/^⁻*or *Pax6⁻^/^⁻* ESCs with CAG-NLS-mScarlet or CAG-GFP respectively and mixed them to form small aggregates. Constitutive fluorescent labels allowed us to FACS-enrich for heterotypic aggregates and trace genotypes in the resultant chimeric NTOs (Fig. 2C). Some aggregates broke upon seeding, producing NTOs with genotype proportions ranging from 0–100%, which we harnessed as an internally controlled system to quantify the relationship between genotype composition and cell-type outcome within the same well.

After RA, salt-and-pepper FOXA2 or PAX6 expression was apparent in d4 *KO:KO* chimeric NTOs, contributed by *Pax6^−/−^* GFP^+^ or *FoxA2^−/−^* mScarlet^+^ lineages, respectively (Fig. 2D). The proportion of each NTO occupied by FOXA2^+^ or PAX6^+^ cells at d4 depended directly on the genotype composition (Fig. 2E), and thus exhibited greater variation than that observed in *WT* NTOs (Fig. 1O). At d6, many *KO:KO* chimeric NTOs exhibited patterning (Fig. 2F, S2B). *Pax6^−/−^* GFP^+^ and *FoxA2^−/−^*mScarlet^+^ lineages spatially segregated into distinct domains (Fig. 2F). FOXA2^+^ cells were organised in discrete clusters, co-expressing SHH (Fig. S2B) indicating mature FP identity that was derived exclusively from the *Pax6^−/−^* GFP^+^ lineage. FP-adjacent bands of OLIG2 expression (Fig. 2F) were observed primarily in the *FoxA2^−/−^*mScarlet^+^ lineage, indicating SHH-induced patterning of motor neuron progenitors that were not observed in either single knockout (Fig. 2A). The efficiency of OLIG2 induction was dependent on the genotype proportions in each chimera, with the highest frequency of OLIG2 rescue when 20-40% of the NTO comprised *Pax6^−/−^*cells (Fig. 2G), i.e. approximately recapitulating the WT proportions (Fig. 1O).

Since RA induces transcriptional changes across numerous genes (Fig. S1), we decided to test whether establishing a mixed population of intermingled FOXA2 or PAX6 expression in the absence of an RA pulse was sufficient to trigger NTO self-organisation. We generated *TetOn-FoxA2:mCherry* + CAG-NLS-mScarlet or *TetOn-Pax6* + CAG-GFP cell lines, mixed them to form ‘*TetOn:TetOn*’ chimeric NTOs, and activated gene expression at d2 with a brief exposure to doxycycline (dox) instead of RA (Fig. 2H). Control dox-treated NTOs consisting only of *TetOn-FoxA2* cells were FOXA2^+^ throughout at d6 and lacked neural progenitors (Fig. S2C). By d6, FOXA2 expression was of endogenous not transgene origin (Fig. S2D), indicating that transient transgene expression initiated endogenous FP differentiation (Epstein et al., 1999). Conversely, purely *TetOn-Pax6* dox-treated NTOs formed neural progenitors but lacked FP at d6 (Fig. S2E).

By contrast, in chimeric *TetOn:TetOn* NTOs, a dox pulse reconstituted FOXA2/PAX6 salt-and-pepper expression, contributed by spatially intermingled *TetOn-FoxA2* mScarlet^+^ vs *TetOn-Pax6* GFP^+^ cells respectively (Fig. 2I). The ratio of FOXA2^+^ vs PAX6^+^ cells in each dox-treated *TetOn:TetOn* chimeric NTO varied as a function of genotype composition (Fig. 2J), whereas RA-treated controls recapitulated *WT* proportions (Fig. 2I-J). By d6, in dox-treated chimeras, *TetOn-FoxA2* mScarlet^+^ and *TetOn-Pax6* GFP^+^ lineages spatially segregated to form FOXA2^+^SHH^+^mScarlet^+^ FP surrounded by GFP^+^ NPs (Fig. 2K). In conclusion, synthetic reconstitution of the FOXA2^+^ vs PAX6^+^ salt-and-pepper expression pattern is sufficient to trigger NTO self-organisation.

Together, these data show that FOXA2 and PAX6 are functional drivers of the diverging differentiation branches initiated by the RA pulse. Thus, NTO patterning can be rescued or reconstituted in *KO*:*KO* or *TetOn*:*TetOn* chimeras by scattered induction of FOXA2^+^PAX6^−^ pFP vs FOXA2^−^PAX6^+^ eNP cell states in appropriate ratios, which together constitute the necessary and sufficient cellular diversity for self-organisation to ensue. The transition from intermingled (Fig. 2D, 2I) to spatially segregated (Fig. 2F, 2K) cell types in chimeric NTOs suggests that cell sorting (Krammer et al., 2024; Xiong et al., 2013) is sufficient to drive spatial self-organisation when fate choices are genetically restricted (*KO*:*KO*) or forced (*TetOn*:*TetOn*).

### Dynamic emergence of pFP and eNP cell states after RA withdrawal

How are cells robustly allocated to both pFP and eNP states in clonal RA-treated *WT* NTOs? This balanced differentiation could arise cell-autonomously, where each cell has an intrinsic probability of adopting a given fate independent of its neighbours, or non-autonomously, where cellular decisions are influenced by the choices of other cells via secreted signals, lateral inhibition or mechanical feedback. To distinguish these possibilities, we first examined the dynamics of cell fate allocation.

To attain a higher temporal resolution of the differentiation trajectories we devised a 5-dimensional flow cytometry antibody panel (OCT4/PAX6/FOXA2/SOX1/SOX2) to quantify cell state proportions during the decision-making process. We assayed every 6h from d2 when RA is added until d4 when pFP versus eNP states emerge (Fig. 3A). The flow cytometry results recapitulated and added temporal resolution to key features identified in the transcriptomic analyses (Fig. 3B-D, S3A). RA-accelerated OCT4 downregulation and PAX6 upregulation were apparent, as was induction of FOXA2 and transient decrease in SOX2 level upon RA withdrawal. FOXA2 and PAX6 co-expression peaked at d3 – d3+6h, and SOX1 upregulation was observed at d3+6h – d4 in the emerging PAX6^+^FOXA2^−^ population (Fig. 3B-D, S3A).

**Figure 3:**
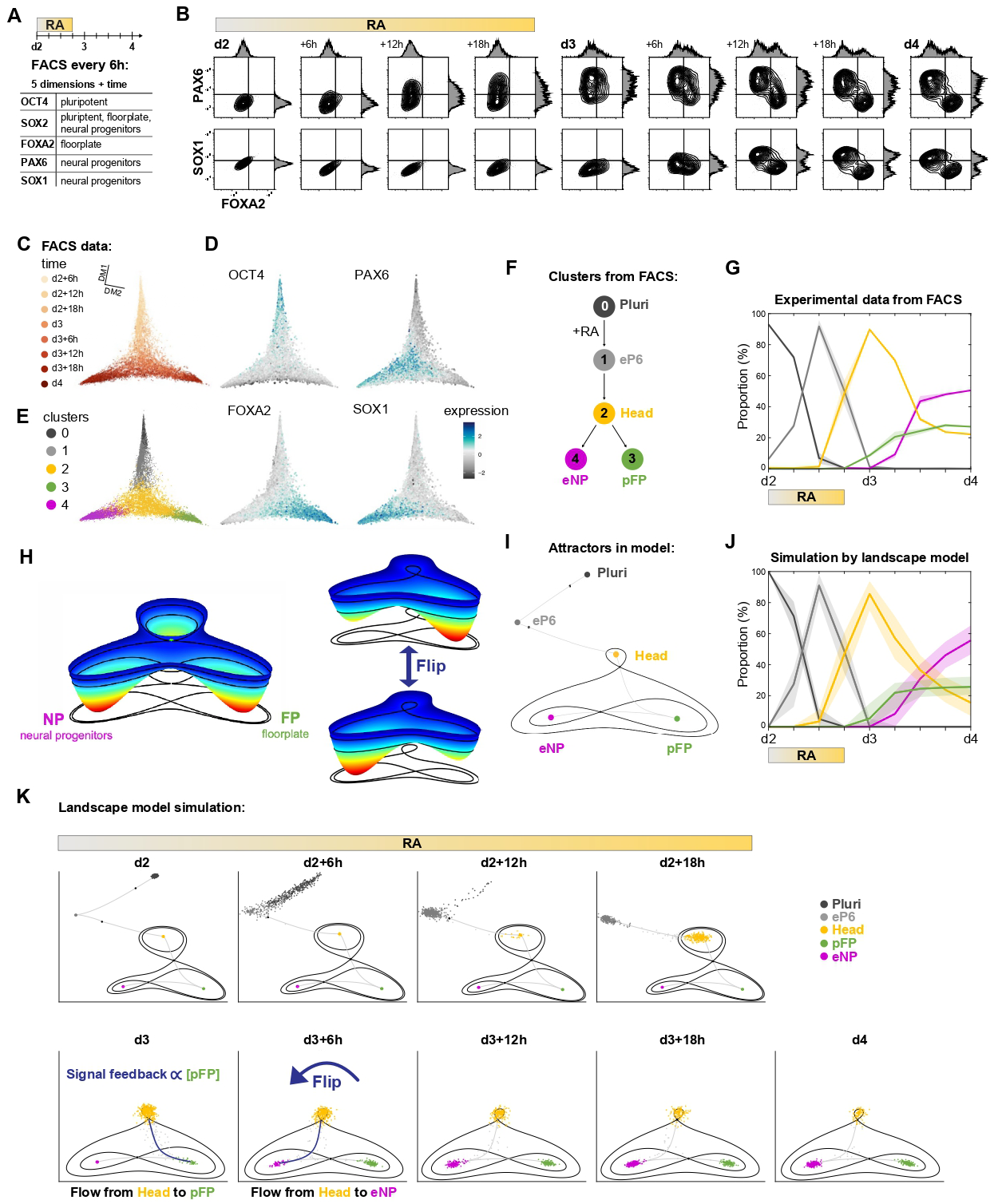
Dynamics and modelling of pFP and eNP cell state emergence. **A-B:** Flow cytometry timecourse of RA-treated NTOs sampled every 6h from d2 to d4. OCT4, SOX2, FOXA2, PAX6 and SOX1 were stained at each timepoint (A). Pairwise contour plots are shown for PAX6 versus FOXA2, and for SOX1 versus FOXA2 (B). **C-D:** Diffusion-map embedding of flow cytometry timecourse data (C) and temporal changes of markers OCT4, PAX6, FOXA2 and SOX1 (D). **E-F:** Five clusters were defined for flow cytometry data based on all assayed markers using a Gaussian mixture model. Colours correspond to the assigned cluster labels. **G:** Time evolution of cell proportions in the different clusters from flow cytometry data. Line corresponds to mean and shaded area to range of replicas. **H:** Dynamical landscape model for the allocation of cells from Head to eNP and pFP. The 3D surface represents the potential of cells transiting the dynamical system. Level curves in the 2D representation show points at the same level as the saddle nodes. A particular illustration of the flip is provided. The landscape tilts changing the cells’ trajectories, showing the way a flip happens in the presented model. For a more general discussion on flips see (Sáez et al., 2022a). **I:** Complete dynamical system with 5 attractors and 4 saddle nodes. The unstable manifolds of the saddle nodes connect the attractors and determine the possible transitions. **J:** Fitting results for simulation of model including feedback from pFP. Line corresponds to mean of simulations, shaded area corresponds to range of all 10000 simulations corresponding to accepted parameter sets during ABC-SMC fitting. **K:** Time evolution of landscape model. Signals affect the topology as shown by unstable manifolds and level curves. Cells move along it by stochasticity and flow determined by the landscape. See also Figure S3.

Visualising cell state transitions as a function of all dimensions of the flow cytometry data (5 markers + time) produced a branching structure in gene expression (Fig. 3C) similar to that observed in the scRNAseq data (Fig. 1B). To identify distinct expression states and examine the dynamics of transitions between them, we used Gaussian mixtures to cluster the data (Fig. S3B-C). We identified five clusters (Fig. 3E-F, S3B-C) in the flow cytometry data: (0) Pluripotent ‘Pluri’ (OCT4^high^SOX2^high^); (1) early Pax6 ‘eP6’ (Oct4^low^SOX2^high^PAX6^low^); (2) ‘Head’ (SOX2^mid^PAX6^mid-high^FOXA2^low-mid^SOX1^low-mid^); (3) ‘pFP’ (SOX2^mid^FOXA2^high^); (4) ‘eNP’ (SOX2^high^PAX6^high^SOX1^high^). The flow cytometry clusters 1-4 approximately correspond to clusters 1-4 in the higher dimensional scRNAseq data, and capture similar proportions of cells at each timepoint (Fig. 1D, 3G). Protein level distributions in the Head population were unimodal but with higher variance than other flow cytometry clusters (Fig. S3B-C), consistent with the variability in FOXA2:PAX6 ratio in double-positive cells observed by staining of d3 NTOs (Fig. 1H) and E8.5 embryos (Fig. 1M).

Quantifying the proportion of cells allocated to each flow cytometry cluster over time (Fig. 3G) suggested an ordering from Pluri to eP6 to Head, followed by emergence of distinct pFP or eNP states (Fig. 3F). State transitions were asynchronous, with some cells occupying the Head state while others had already begun populating pFP or eNP (Fig. 3G). The high temporal resolution revealed that allocation to the pFP cluster started earlier than allocation to the eNP cluster. Subsequently, both pFP and eNP clusters increased until pFP plateaued at 25-30%, whereas eNP continued increasing (Fig. 3G). A simple probabilistic model where each cell has an autonomous 25% chance of adopting pFP fate could explain the final proportions. However, this model would not capture the observed dynamics. In the experimental data, pFP allocation occurred first and saturated, after which eNP allocation continued. This sequential filling pattern suggests that fate decisions are not autonomous but instead depend either on time or the current composition of the organoid. This raises the possibility of tissue-level feedback mechanisms that regulate the timing and order of cell fate specification.

### A dynamical systems model of cell state transitions in NTOs

To understand the observed dynamics of pFP vs eNP allocation, we asked whether a mathematical model could fit the data and make meaningful predictions. We turned to dynamical systems theory, modelling each state as an attractor in a Waddington-inspired geometric landscape (see SI for details) (Fig. 3H-I). Briefly, saddle nodes and their unstable manifolds connect attractors, like passes between valleys in the Waddington metaphor (Sáez et al., 2022a). The flow of cells through the landscape is stochastic and guided by the shape of the underlying landscape, which is determined by the signalling environment through the parameters of the model. When an attractor becomes sufficiently shallow or disappears (bifurcation), cells flow out and transition via the saddle node to another available attractor. We built a landscape (Fig. 3I) informed by the scRNAseq/ flow cytometry clusters (Fig. 1C, 3E): attractors ordered sequentially for Pluri to eP6 to Head, then branching from Head to pFP or eNP with a ‘flip’ topology to allow allocation to both attractors. We then screened parameter space for a configuration that could fit the experimental data (see SI for details).

With model parameters dependent only on the RA level in the media, simulated and experimental data showed good agreement for Pluri, eP6 and Head states, but the model was unable to recapitulate the experimental observation that cells initially accumulated in the pFP state faster than in the eNP state, with this pattern reversing later (Fig. S3D). This discrepancy indicated that the landscape must change over time, initially favouring flow from Head to pFP before reshaping to favour Head to eNP allocation. We therefore tested whether including feedback between pFP allocation and fate dynamics could account for these differences. We modelled this as a signal produced proportional to pFP cells that tilts the landscape in favour of the eNP state (see SI for details). Through this, early filling of cells from the Head to pFP state then alters (‘flips’) the landscape to favour flow of cells from Head to eNP (Supplementary Video 1) (see SI for details). Addition of this feedback parameter was sufficient for the model to accurately recapitulate the experimental data (Fig. 3J-K, S3E-F). Thus, model construction suggests that allocation of cells to pFP vs eNP is a regulative process, with signalling feedback emanating from pFP cells to limit their own proportion.

### Regulative feedback controls proportional allocation to pFP and eNP states

Regulative feedback implies sensing and modulation of decisions based on other cells in the tissue. To test whether pFP versus eNP allocation is regulative, we made chimeric NTOs mixing *WT* + CAG-eGFP cells, to which both states are available, with *FoxA2^−/−^* + CAG-NLS-mScarlet cells that cannot make pFP (Fig. 2A-B, 4A). We quantified the decisions made by *WT* cells as a function of the ratio of *WT*:*FoxA2^−/−^* cells in each chimera. If fate allocation is cell autonomous, we would expect the same proportion of the *WT* cells to make pFP irrespective of chimera composition. Conversely, if allocation is regulative, we would expect *WT* proportions of pFP to change to compensate for the limitations of their neighbours (Fig. 4A).

**Figure 4:**
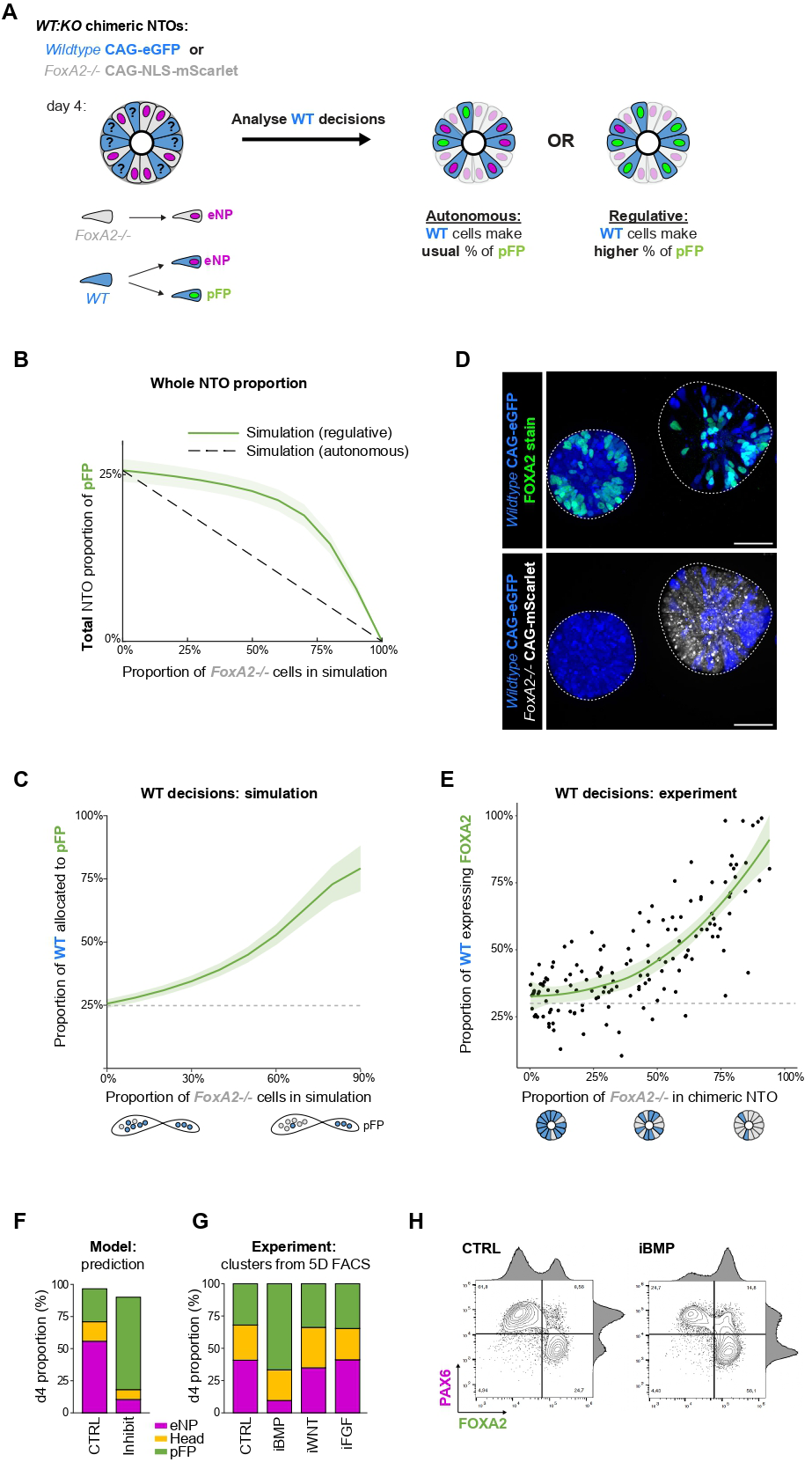
Regulative feedback balances generation of pFP and eNP. **A:** *WT:KO* chimeric NTOs are made by combining wildtype (*WT*) and *FoxA2^−/−^* (*KO*) ESCs in heterotypic aggregates. *KO* cells cannot make pFP, and the proportion of *KO* cells in *WT:KO* chimeras modulates tissue-level properties. Possible outcomes for *WT* cell decisions within *WT:KO* chimeras are indicated, to distinguish whether eNP versus pFP allocation is autonomous or regulative. **B-C:** Model predictions of pFP allocation in *WT:KO* chimeric NTOs at d4, as a function of the chimeric composition. The total proportion of pFP in the whole NTO (*WT*+*KO*) is shown in (B). The proportion of *WT* cells allocated to pFP is shown in (C). Green line: prediction if *WT* decisions are modulated by regulative feedback. Grey dashed line: prediction if *WT* decisions are autonomous. Shaded area corresponds to variability of the different accepted parameters during ABC SMC fitting. **D:** Immunofluorescence on RA-treated *WT:KO* chimeric NTOs at d4. *WT* cells can be traced by their constitutive CAG-eGFP label (blue), and *FoxA2^−/−^ KO* cells can be traced by their constitutive CAG-NLS-mScarlet label (red). FOXA2 (green) expression is stained. Representative single confocal *z* slice, scale bar 100μm. **E:** Proportion of *WT* cells that are FOXA2^+^ in each *WT:KO* chimeric NTO at d4, as a function of the chimeric composition, quantified from 3D images of wholemount immunofluorescence (n=193). **F:** Model prediction of cell proportions allocated to Head, pFP or eNP states at d4 when the strength of the regulative signalling feedback parameter is reduced (equivalent to “inhibiting” the signal). **G-H:** Experimental inhibition (i) of candidate pathways for the regulative feedback signal. RA-treated NTOs were treated with DMSO (CTRL), 100nM LDN (iBMP), 5uM IWP2 (iWNT) or 1μM PD0325901 (iFGF) then analysed at d4 by flow cytometry with immunostaining for OCT4, SOX2, FOXA2, PAX6 and SOX1. Clusters were defined for flow cytometry data on all 5 markers using a Gaussian mixture model, and the proportion of cells allocated to Head, pFP and eNP are shown (G). Pairwise contour plots are shown for PAX6 versus FOXA2 (H). See also Figure S4.

To make quantitative predictions, we simulated *WT:FoxA2^−/−^*chimeric NTOs in the dynamical landscape model. We modelled *FoxA2^−/−^*cells as unable to enter the pFP attractor (see SI for details), whilst *WT* cells were free to enter any part of the landscape. We ran simulations with ratios of *WT:FoxA2^−/−^* cells ranging from 10-90% (Fig. 4B). A wide range of chimeric composition (40-90% *WT* : 60-10% *FoxA2^−/−^*) produced a narrow range of outcomes (21-26% total pFP) (Fig. 4B), with total pFP dropping below this range only when <30% of cells were *WT*. Focusing on the decisions made by *WT* cells, the lower the *WT:FoxA2^−/−^* ratio, the higher the proportion of *WT* cells allocated to pFP at d4 (Fig. 4C). Examining the simulation dynamics leading up to d4, this was because the pFP attractor continued to fill with cells (*WT* but not *FoxA2^−/−^*) until it contained ∼25% of the total (*WT*+*FoxA2^−/−^*) cell population at which point the flow of cells favoured eNP.

Experimental data agreed with model predictions. The higher the proportion of *FoxA2^−/−^* CAG-NLS-mScarlet cells (total mScarlet^+^ / total NTO), the higher the proportion of the *WT* CAG-eGFP cells allocated to pFP (eGFP^+^FOXA2^+^PAX6^−^ / total eGFP^+^) in chimeric NTOs (Fig. 4D-E). Therefore, proportional allocation to pFP is subject to regulative feedback control, with decisions of *WT* cells influenced by the genotypes of (i.e. by the fates available to and communicated by) their neighbours, in a dose-dependent manner. Thus, regulative feedback canalises tissue-level outcome despite genetic heterogeneity, ensuring balanced emergence of both pFP and eNP cell states that is necessary (Fig. 2A-G) and sufficient (Fig. 2H-K) for correct NTO self-organisation and patterning to ensue.

### BMP signalling provides regulative feedback for pFP and eNP allocation

To identify the regulative signal, we screened the scRNAseq data for signatures of differential signalling pathway expression and activity, complemented with ligand-receptor analyses (Fig. S4A-B). The model predicted the signal to emanate from cells in the pFP state (∼cluster 3), and predicted that reducing the strength of the feedback would result in a higher proportion of cells allocated to pFP and a lower proportion to eNP (Fig. 4F). We experimentally inhibited candidate pathways from d2.5 and analysed the impact on cell state allocation at d4 by multidimensional flow cytometry (Fig. 4G). WNT or FGF inhibition had no significant impact. By contrast, BMP inhibition specifically increased the proportion of pFP, at the expense of eNP (Fig. 4G-H). This is consistent with a role of BMP signalling in limiting the proportion of cells normally allocated to pFP as the Head state emerges then empties after RA withdrawal (Fig. 3G). BMP signalling was previously identified as mediating competitive interactions between FOXA2^+^ clusters later in the NTO self-organisation process from d4–6 (Krammer et al., 2024), together implicating the same pathway in governing both diversity generation and spatial resolution.

### RA gates entry and exit from the Head state

The dynamical landscape model predicted that RA controls entry and exit of cells from the Head state but does not directly determine the final proportions of pFP and eNP cells. This suggests that RA acts as a temporal trigger for fate diversification rather than a dose-dependent instructor of specific fate choices. Experimental data (Fig. 3G) and simulated dynamics (Fig. 3J) agree that filling of the Head state commences during the RA pulse, peaks at d3 after RA withdrawal, then declines as pFP then eNP start to fill. We tested experimentally whether these transitions are causally related to RA withdrawal or merely a coincidence. We added RA at day 2, then withdrew it after 6h or 24h, rather than 18h. The model correctly predicted that pFP allocation commenced after RA withdrawal, irrespective of RA duration (Fig. S5A-B). Short or extended RA exposure both yielded similar d4 proportions of pFP and eNP, though with shifted dynamics (Fig. S5A-B). Invariance in pFP and eNP d4 proportions was further demonstrated with 10-fold higher RA concentration (Fig. S5C). Together these data confirm that the primary action of the RA pulse is to initiate the symmetry breaking event that diversifies cell types but does not instruct the balance between these fates.

### Transient co-expression of FOXA2 and PAX6 is sufficient to initiate NTO self-organisation

Finally, we asked whether the RA pulse can be bypassed by directly establishing cells in the Head state. In model simulations, initiating all cells in the Head attractor is sufficient to recapitulate pFP vs eNP allocation in characteristic proportions (Fig. 5A–B), thanks to asynchronous exit from the shallow attractor and contemporaneous regulative feedback from pFP. The Head state is characterised by broad distributions of PAX6 and FOXA2 expression including cells co-expressing both (Fig. 3D-E, S3B-C, S3F). Therefore, to test this prediction experimentally, we engineered ‘*2xTetOn*’ ESCs containing both *TetOn-Pax6* and *TetOn-FoxA2:mCherry* transgenes in the same cell, and formed clonal NTOs (Fig. 5C). 6h after a brief pulse of dox (instead of RA), *2xTetOn* NTOs exhibited a high proportion (97%) of PAX6⁺FOXA2⁺ double-positive cells (Fig. 5D–E), with levels appearing more uniform than in RA-treated NTOs (Fig. 3B). By d3, however, heterogeneity in FOXA2:PAX6 ratios emerged between cells in clonal *2xTetOn* NTOs, recapitulating the variability characteristic of the Head state and creating a salt-and-pepper spatial distribution (Fig. 5F).

**Figure 5:**
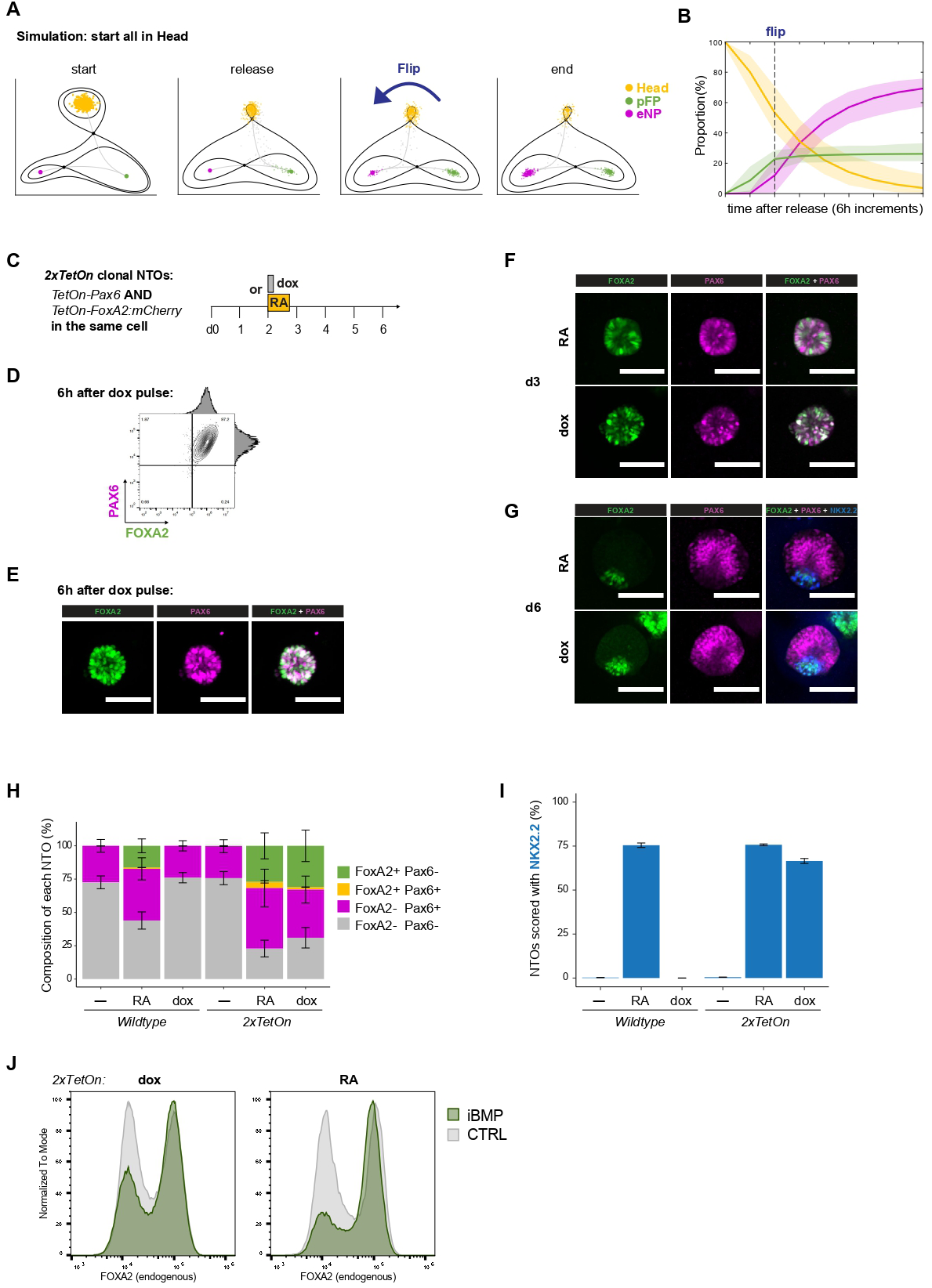
Transient co-expression of FOXA2 and PAX6 is sufficient to initiate self-organisation. **A-B:** Model prediction for starting all cells directly in the Head attractor and allowing its release. The topology of the landscape and how signals affect it are shown in (A). The fitting results for simulations are shown in (B), with shaded area corresponding to variability of the different accepted parameters during ABC SMC fitting. **C:** *2xTetOn* clonal NTOs are formed from single ESCs which contain *TetOn-FoxA2:mCherry* and *TetOn-Pax6* transgenes, for doxycycline (dox)-inducible co-expression of both transgenes at once. **D:** Flow cytometry of FOXA2 and PAX6 immunostaining in clonal *2xTetOn* NTOs, 6h after a brief pulse of doxycycline at d2. **E:** Immunofluorescence of FOXA2 (green) and PAX6 (magenta) in clonal *2xTetOn* NTOs, 6h after a brief pulse of doxycycline. Representative NTO, acquired with 10x objective, scale bar 100μm. **F:** Immunofluorescence of FOXA2 (green) and PAX6 (magenta) at d3 in clonal *2xTetOn* NTOs treated with RA or dox. Representative NTO, acquired with 10x objective, scale bar 100μm. **G:** Immunofluorescence of FOXA2 (green), PAX6 (magenta) and NKX2.2 (blue) at d6 in clonal *2xTetOn* NTOs treated with RA or dox. Representative NTO, acquired with 10x objective, scale bar 100μm. **H:** Quantification of the proportion of cell states in each NTO at d6, for parental wildtype and *2xTetOn* NTOs treated on d2 with a pulse of RA or dox or neither (−), quantified from 3D images of wholemount immunofluorescence. The error bars represent the variance of cell state proportions (n = 3006). **I:** Quantification of NTOs that express NKX2.2 at d6 (presence/absence of marker), for wildtype and *2xTetOn* NTOs treated on d2 with a pulse of with RA or dox or neither (−). The error bars represent the variance of cell proportions between images. n =3006. **J:** Flow cytometry analysis for FOXA2 immunostaining on *2xTetOn* NTOs treated with 100nM LDN (iBMP) after RA or dox. Analysis was performed at d5, when no mCherry signal is detected, to ensure that FOXA2 immunostaining signal derives from endogenous rather than transgenic FOXA2 expression. See also Figure S5.

Remarkably, dox-treated clonal *2xTetOn* NTOs resolved to form a spatially organised FOXA2^+^ versus PAX6^+^ domains by d6 (Fig. 5G-H). SHH-responsive NKX2.2 expression was induced, indicating FP functionality (Fig. 5G). The endpoint patterning efficiency was similar to RA-treated controls (Fig. 5H-I). Thus, transient PAX6^+^FOXA2^+^ co-expression is sufficient to trigger cell state diversification and efficient spatial self-organisation and patterning in clonal NTOs without an RA pulse.

To test whether regulative feedback also mediates the proportional allocation to pFP vs eNP after transient transgene co-expression, we inhibited BMP signalling after the dox pulse. Similar to RA-treated NTOs, BMP inhibition after dox treatment of clonal *2xTetOn* NTOs resulted in a higher proportion of cells adopting an endogenous FOXA2^+^ (pFP) state (Fig. 5J). Therefore, after transient PAX6^+^FOXA2^+^ transgene co-expression kick-starts the process, subsequent proportional allocation of cell states is not cell-autonomous, and is governed by regulative feedback as endogenous expression programs take over.

Together, we conclude that the minimal essential circuitry to initiate NTO self-organisation comprises two transcription factors, PAX6 and FOXA2, and regulative feedback for proportional allocation to the cell states they govern.

## Discussion

In this study, we deconstructed and reconstructed the requirements for NTO self-organisation. We identified early neural progenitors (eNPs) and floorplate precursors (pFPs) as the two major cell states produced after RA triggers diversification. PAX6 and FOXA2 governed the specification of these cell types. Strikingly, placing PAX6 and FOXA2 under exogenous control was sufficient to reconstitute the symmetry breaking event and initiate regulated self-organisation in the absence of an RA pulse. In wildtype NTOs, symmetry breaking and proportion control were robust. A spatially intermingled mix of PAX6^+^ eNPs and FOXA2^+^ pFPs were observed in 97% of wildtype NTOs at d4, with an average composition of 25% pFP in each NTO (Fig. 1N-O). This proportion was ensured by regulative feedback (mediated by BMP) rather than cell-autonomous mechanisms. Together, this offers insight into how reproducible cellular diversity is achieved through symmetry breaking and regulative feedback, providing a model to understand how clonal cell populations generate heterogeneous tissues.

We used dynamical landscape analysis to develop a quantitative understanding of eNP and pFP allocation in NTOs. This transformed complex gene expression data into a predictive model of cell state transitions. Preceding diversification of cell states to eNPs and pFPs, a pulse of RA signalling permitted cells to adopt a transient ‘Head’ state. This was characterised by a distinct gene expression signature involving broad distributions of PAX6 and FOXA2 protein levels including double-positive cells. Asynchronous exit from the Head state allowed proportion regulation during diversification. Signalling feedback, mediated by BMP from the initially generated pFPs, reshaped the landscape to promote eNP specification of subsequent cells exiting the Head state. This ensured reproducible cellular diversity within each NTO. The model accurately predicted experimental outcomes including: how wildtype cells compensate for *FoxA2^−/−^* neighbours in chimeric NTOs; how BMP inhibition shifted proportions toward increased pFP; and that similar ratios of pFP:eNP emerged despite varying RA durations and concentrations. Thus, RA and BMP each modulate orthogonal “axes” of the landscape: RA controls progression through the Head state to initiate diversification, while BMP regulates proportional allocation between fates.

These findings allowed us to reduce the NTO system to its core components. Transient co-expression of FOXA2 and PAX6 in clonal ‘*2xTetOn*’ NTOs was sufficient to reconstitute self-organisation without an RA pulse. Transient co-expression resolved into distinct pFP and eNP states, under regulative feedback control from BMP signalling, and these spatially organised into a patterned tissue. Hence, RA dependence can be bypassed synthetically, identifying the minimal molecular requirements: two transcription factors plus regulative feedback. Importantly, chimeric experiments revealed that the correct ratio of cell types, rather than their specific molecular origins, drives successful patterning. Both *‘KO:KO*’ chimeras, where genetic restrictions force lineage choices (Fig. 2C-G), and *‘TetOn:TetOn’* chimeras, where transgenes dictate initial fates (Fig. 2H-K), achieved proper tissue organisation when they recapitulated wildtype pFP:eNP proportions, demonstrating convergence on tissue-level outcomes from different molecular starting points.

This circuit design we uncovered – dual transcription factors initiating opposing fates under proportional control by regulative signalling – appears to represent a fundamental strategy for reliably generating the cellular diversity required for emergent tissue organisation. Similar dynamical mechanisms have been implicated in the allocation of inner cell mass (ICM) cells to the epiblast and primitive endoderm in the mouse blastocyst (Raju and Siggia, 2024) and the allocation of neuromesodermal progenitors (NMPs) to neural and mesodermal tissue (Sáez et al., 2022b). The parallels are striking. Each of these systems exhibit a Head state that is marginally stable, which leads cells to a critical decision point that the cell population crosses asynchronously. Regulative signals (FGF in blastocysts; BMP in NTOs; WNT and RA in NMPs) then shape the proportional allocation of fates. The underlying design motif in each case - the flip dynamical landscape - ensures robust diversification and proportional cell fate allocation. The inherent two-dimensionality of this landscape allows feedback regulation while maintaining sensitivity to perturbations at specific temporal windows. These features are essential for both developmental robustness and environmental responsiveness. The convergence of disparate developmental contexts on the same underlying decision topology suggests it represents a fundamental and adaptable design principle for multicellular systems to achieve reproducible outcomes.

The transient co-expression of two opposing transcription factors is a recurring motif throughout development when two lineages differentiate from a progenitor pool (Briggs et al., 2018; Chazaud et al., 2006; Farrell et al., 2018; Henrique et al., 2015; Hu et al., 1997; Soldatov et al., 2019). Much like the FOXA2^+^PAX6^+^ co-expression that we describe in NTOs and embryos, opposing TF expression in the ICM (NANOG^+^GATA6^+^) and NMPs (SOX2^+^T^+^) are also transient and exhibit varied levels in a temporal snapshot (Plusa et al., 2008; Wymeersch et al., 2016). Expression levels can correlate with lineage bias if left unperturbed, although plasticity may be observed upon change of environment (Binagui-Casas et al., 2025; Kim-Yip et al., 2025; Linneberg-Agerholm et al., 2024; Wymeersch et al., 2016), and careful consideration must be given to bipotency on the population versus single cell levels. In general, variability in expression levels may reflect the importance of noise to generate order, and to confer the plasticity necessary for regulative development (Balázsi et al., 2011; Paszek et al., 2010). In the future, live-imaging and lineage-tracing of NTOs should elucidate whether fate correlates with expression levels within the Head population, and how outcomes adjust in response to signalling perturbations or genetically compromised neighbours to ensure robust tissue outcomes.

*In vitro* stem cell models offer a powerful opportunity to uncover how order emerges in cell collectives, revealing mechanisms that may contribute *in vivo* and that can be harnessed for tissue engineering (Hofer and Lutolf, 2021; Morales et al., 2021). Here, the FOXA2^+^PAX6^+^ signature that we discovered in NTOs led to identification of an analogous cell state *in vivo* in the neural plate at E8.5, complementing a recent report of FOXA2^+^SOX1^+^ co-expression (Arekatla et al., 2024). Since transient FOXA2^+^PAX6^+^ co-expression is functionally sufficient to reconstitute NTO self-organisation *in vitro*, determining the role of this transient state *in vivo* represents an important future direction. We also observed scattered FOXA2 expression in the embryonic ventral neural plate, mirroring the salt-and-pepper stage of NTO formation (Fig. 1H, 1L), and previously we showed that BMP pathway perturbation alters FP proportion (Krammer et al., 2024). The *in vivo* identification of multiple key NTO hallmarks provides a strong mandate for future investigation into the contribution and guidance of self-organising processes for robust neural development.

Together, this study demonstrates that symmetry breaking and regulative feedback, mediated by a simple transcription factor circuit, enables clonal populations to reliably generate the cellular diversity required for emergent tissue organisation. NTOs serve as a powerful quantitative platform for studying self-organisation, accessible and perturbable across scales from molecular governance of stem cell decisions to tissue-level properties. Clonal NTOs eliminate confounding pre-existing heterogeneity, whereas chimeric NTOs allow separable manipulation of cell- and tissue-level properties to study bidirectional causality across scales. This work opens new avenues for understanding development and engineering tissues, including investigating whether similar regulative mechanisms operate in other organoid systems, and exploring how external context and constraints interact with internal molecular feedback.

### Limitations of the study

The dynamical systems model assumes that all cells experience the same signalling environment, proportional to the number of cells in the pFP attractor but agnostic to their relative position in the tissue. Given the tendency of FOXA2^+^ cells to cluster as self-organisation proceeds (Krammer et al., 2024), local signalling differences may emerge across the epithelium (Abdel Fattah et al., 2023). Future directions to integrate dynamical landscapes within ‘cells’ using agent-based models could provide a more holistic view of the continuous process from fate diversification to spatial rearrangement in NTOs, ideally also integrating proliferation and lineage information. More broadly, an important challenge will be to experimentally distinguish cells that have settled into attractors from those transitioning between states; developing data-analytic tools capable of making this distinction will be essential for rigorously connecting dynamical systems models to experimental measurements.

In this work, we focussed our efforts on transcription factors and signalling pathways governing cell state transitions. We identified BMP signalling as a contributor to the regulative nature of pFP versus eNP allocation, but do not exclude that other pathways or properties may also play a role. Recent work has highlighted the relevance of metabolism to modulate lineage proportions in gastruloids (Dingare et al., 2024; Stapornwongkul et al., 2025; Villaronga-Luque et al., 2025). Mechanical forces should also be considered: tissue stretching can increase FOXA2 proportion in human NTOs (Abdel Fattah et al., 2021), and, in a different setting, the decoding of density-dependent mechanosensor heterogeneity by FOXA1 ensures diversification of cell states during intestinal regeneration (Schwayer et al., 2025), leading to an emerging picture of FOXA factors as widely involved in biological symmetry breaking processes with a degree of mechanical regulation.

## Supporting information

Dynamical Landscape Movie

## Acknowledgements

We thank James Sharpe, Kristina Stapornwongkul, Katharina Lust, Pietro Tardivo, and members of the Briscoe and Tanaka groups for critically reading and constructive comments. We thank Eric Siggia for scientific input, particularly for encouraging the creation of chimeric NTOs. We are grateful to Fernando Becerril Perez, Jiaye Yang and Birgit Ritschka for assisting with embryo harvests, and to Anja Pelzl for tissue culture support. We thank Marco Antonio Leyva Gonzalez for construct design, and Lokesh Pimpale for image analysis advice. We are grateful to Graziano Martello for early discussion and PB-CAG-eGFP plasmid, and to Lucrezia Galli for clearing and wholemount imaging advice. This work would not have been possible without the outstanding support from VBCF BioOptics and NGS facilities, in particular: Alberto C. Moreno, Pawel Pasierbek, Gabriele Bradamante for microscopy and more; FACS and NGS teams for coordination to generate high quality scRNAseq libraries from challenging timecourse samples. For the purpose of Open Access, the authors have applied a CC BY public copyright license to any Author Accepted Manuscript (AAM) version arising from this submission. No AI tools were used in the writing of this manuscript.

## Author contributions

HTS, EMT, JB conceived and supervised the project, acquired funding and interpreted data. HTS and EC designed and performed experiments, acquired and analysed data. JW performed formal analysis of transcriptomic, imaging and flow cytometry data. MS established theoretical results and performed dynamical modelling simulations, with advice from DR. TK and MJD generated knockout cell lines and provided conceptual input. MJD and GS contributed to flow cytometry methodology for multi-metric stainings and chimeric NTO generation respectively. MM and KI assisted in molecular cloning. TL, KI, JCS contributed to image analysis pipelines. HTS, JB, MS, EMT wrote the manuscript, with input from JCS, JW, EC. All authors reviewed and approved the manuscript.

## Data and materials availability

Reagents such as cell lines and plasmids generated in this study are available; requests for biological materials should be addressed to Elly Tanaka and Hannah Stuart.

RNA-seq data generated in this study have been deposited at GEO under accession code GSEXXXXXX.

Upon publication, original code for dynamical systems modelling and bioinformatic analyses will be available at Github, imaging data deposited at BioImage Archive, flow cytometry data deposited at FigShare. Any additional information required to re-analyse the data is available upon request from Hannah Stuart, Meritxell Sáez, James Briscoe and Elly Tanaka.

---

**<u>Figure S1</u>**

**A:** UMAP embedding of scRNA-seq timecourse data from d2 (before RA-treatment) until d6, with or without an RA pulse applied at d2 for 18h.

**B:** Temporal profiles of differentially expressed TFs and genes in signalling pathways in RA-treated versus -untreated trajectories ordered according to pseudotime based on principle curve analysis (d2-4). Clusters were identified with the hierarchical clustering approach by specifying 7 clusters.

**C:** Expression feature plots on the same UMAP embedding as (A). Representative marker genes are shown for pluripotency, RA response, anterior-posterior (AP) identity, neural differentiation, floorplate, and dorsal-ventral (DV) patterning of neural progenitors.

**D:** Pseudotime of scRNA-seq (d2-5) calculated by Palantir, shown on same UMAP embedding as (A). **E:** Cellular heterogeneity (i.e. pairwise distances calculated with top 20 principle components) were compared between RA-treated versus -untreated samples by discretizing Palantir pseudotime into 10 bins for each condition.

**F:** Heatmap of top 20 marker genes for each cluster identified in Fig 1C.

**G-H:** UMAP projection (H) using 108 sparse features identified by three independent methods (geneBasis, Dubstep and SMD), from which representative features are highlighted by dot plot in (I). **I:** Violin plot showing expression of *Oct4*, *FoxA2*, *Pax6* and *Sox1* in scRNA-seq data from d2.5-6 for RA-treated versus -untreated NTOs.

**J**: Quantification of FOXA2 and PAX6 expression level in every nucleus of the neural tube at the levels of somite 1-4 (hindbrain) of 7-somite stage embryos (n=2, see also Fig. 1M), from wholemount confocal acquisition with 30x objective and 0.41μm *z*-step. Nuclei were segmented on SOX2 staining using CellPose 3D, with manual curation to remove SOX2^+^ nuclei from other tissues than the neural tube. See Table S2 for further scoring of additional embryos (n=20).

Related to Figure 1.

**<u>Figure S2</u>**

**A:** Flow cytometry analysis of SOX1 immunostaining at d3 and d4 on wildtype, *FoxA2^−/−^* and *Pax6^−/−^*

NTOs without an RA pulse.

**B:** Immunofluorescence at d6 of RA-treated *KO:KO* chimeric NTOs. *FoxA2^−/−^* cells can be traced by their constitutive CAG-NLS-mScarlet label, and *Pax6^−/−^* cells by their constitutive CAG-eGFP label. FOXA2 (green), SHH (grey) or NKX2.2 (grey) expression are stained. Representative maximum intensity *z*-projection.

**C:** Immunofluorescence at d6 of FOXA2 (green) and PAX6 (magenta) on *TetOn-FoxA2:mCherry* NTOs treated on d2 with a pulse of RA or dox. Representative maximum intensity *z*-projection, acquired with 10x objective, scale bar 100μm.

**D:** Immunofluorescence of FOXA2 (green) and mCherry (red) of *TetOn-FoxA2:mCherry* NTOs treated on d2 with a pulse of dox. FOXA2 antibody stains both transgenic and endogenous FOXA2. mCherry (fusion protein) denotes presence (d3) or absence (d6) of transgene expression. Representative NTO, single confocal *z* slice acquired with 10x objective, scale bar 50μm.

**E:** Immunofluorescence at d6 of FOXA2 (green) and PAX6 (magenta) on *TetOn-Pax6* NTOs treated on d2 with a pulse of RA or dox. Representative maximum intensity *z*-projection, acquired with 10x objective, scale bar 100μm.

Related to Figure 2.

**<u>Figure S3</u>**

**A:** Flow cytometry timecourse on NTOs sampled every 6h from d2 to d4, with (red) or without (blue) RA treatment. OCT4, SOX2, FOXA2, PAX6 were co-stained at each timepoint. Pairwise contour plots are shown for OCT4 versus FOXA2, and for PAX6 versus SOX1.

**B:** Five clusters were defined for flow cytometry data using the Gaussian mixture model. Colours correspond to the assigned cluster labels. The expression distributions of cells in each cluster are unimodal for all markers.

**C:** Contour plots are shown for all pairwise combinations of flow cytometry markers, for all RA-treated flow cytometry data pooled between timepoints. Colours correspond to the assigned cluster labels.

**D:** Fitting results for simulation of model without feedback from pFP. Line corresponds to mean of simulations, shaded area corresponds to range of all 10000 simulations corresponding to accepted parameter sets during ABC-SMC fitting.

**E:** LDA projection of flow cytometry data showing evolution through the different parts of the corresponding landscape. For intermediate time points, Head cells are represented in both landscapes. Level curves show areas corresponding to each cluster (attractor in the landscape). Colours correspond to the assigned cluster labels. Level curves show the distribution of the data pooled between timepoints.

**F:** LDA projection for Head, pFP and eNP clusters. Grey background shows the complete cell population at all time points. Coloured (black/red) cells correspond to Head state clustered cells at the indicated timepoint. Red cells co-express both FOXA2 and PAX6. Level curves show the distribution of cells at each time point.

Related to Figure 3.

**<u>Figure S4</u>**

**A:** Relative strength of signalling pathways (BMP, FGF, WNT and Hedgehog (HH)) using CellChat across clusters identified in scRNA-seq of RA-treated NTOs (Fig. 1C).

**B:** Circos plot of top 150 ligand-receptor pairs between cluster 3 and 4 in scRNA-seq data with LIANA. Colours correspond to cluster labels. The arrows emanate from source cells and end with receiving cells.

Related to Figure 4.

**<u>Figure S5</u>**

**A:** Model prediction for withdrawing RA at different times.

**B-C:** Pairwise contour plots are shown for PAX6 versus FOXA2 immunostaining analysed by flow cytometry at the indicated timepoints, after treatment with different durations (B) or doses (C) of RA.

Related to Figure 5.

**<u>Supplementary Video 1</u>**

**Left**: Stochastic simulation of 500 cells from d2-4 using the landscape model obtained from the average of all accepted parameters in the fitting algorithm. All attractors are shown. Level curves and unstable manifolds for the flip landscape are included. Cells are coloured according to the state assigned in clustering as indicated in right panel. The signalling regime is indicated by background shading (yellow when RA is applied). The feedback (flip) is highlighted by colouring the changing unstable manifold. **Right**: Proportions of the different cell states along the simulation. Lines are coloured according to cell state as indicated in legend. Background shading indicates signalling regime (yellow when RA is applied).

## Materials and Methods

### Cell culture

#### Cell lines

HM1 mouse embryonic cell line (ESC) was used for all experiments and as background of all transgenic and knockout lines. Cell lines were routinely tested for mycoplasma and confirmed negative.

#### Maintenance

ESCs were cultured in N2B27 medium + 2i and LIF and kept at 37C at 5% CO2.

Standards N2B27 was composed of 1:1 DMEM/F-12 and Neurobasal (Gibco), 0.5% v/v N2 (Gibco), 1% v/v B27 (Gibco), 2 mM L-Glutamine (Gibco), 1% penicillin/streptomycin (Sigma) and 0.1 mM ϐ-mercaptoethanol. N2B27 base media was then supplemented with 3 μM CHIR99021 (Tocris), 1 μM PD0325901 (Tocris) and 20 ng/ml murine LIF (Qkine).

Tissue culture plates were pre-coated with 0.15% gelatin (Sigma) in PBS at room temperature for at least 10 min and coating was removed immediately before seeding.

Cells were seeded at a density of 20,000 or 30,000 cells per 6-well and media was exchanged daily. Passaging was performed every third day, incubating cells with Accutase (Gibco) for 3 min at 37C and inactivating the enzyme with base media. Cells were resuspended until dissociated into single cells, centrifuged at 1500 rpm for 5 min at room temperature and plated in a new well.

#### Generation of transgenic cell lines

PiggyBac-TetON-destination-PGK-hygro, PiggyBac-TetON-destination-PGK-zeo, PiggyBac-CAG-destination-hygro and PiggyBac-CAG-rtTA3-PGK-puro vectors were used from Stuart et al 2019. PiggyBac-TetOn-FoxA2:mCherry-hygro was from Delas et al 2023. PiggyBac-CAG-eGFP-hygro was a gift from the Martello lab.

New transgenes generated in this work:

PiggyBac-TetOn-Pax6-zeo
PiggyBac-CAG-NLS-mNeonGreen-bsd
PiggyBac-CAG-NLS-mScarlet-hygro

Expression vectors were generated by Gateway cloning (Invitrogen), by inserting coding sequences into the respective destination vector.

Human PAX6 entry vector was obtained from ORFeome Collab (Cat. n. OHS5893-202492609). 3xNLS-mScarlet-I was cloned into pDONR221 entry vector from 3xNLS-mScarlet-I sequence from Addgene #98816 plasmid, using following primers:

- Forward:
- GGGGACAAGTTTGTACAAAAAAGCAGGCTTCACCatgggatcagatccaaaaaagaagagaaagg
- Reverse: GGGGACCACTTTGTACAAGAAAGCTGGGTCttacttgtacagctcgtccatgcc

3xNLS-mNeonGreen was cloned into pDONR221 entry vector from 3xNLS-mNeonGreen sequence from Addgene #98875 plasmid, using following primers:

- Forward: GGGGACAAGTTTGTACAAAAAAGCAGGCTTCACCatgggatcagatccaaaaaagaagagaaagg
- Reverse: GGGGACCACTTTGTACAAGAAAGCTGGGTCttacttgtacagctcgtccatgcc

Generation of stable transgenic ESC lines was achieved by transfection as described: 1 μg expression vector, 1 μg rtTA3 vector, 1 μg PBase transposase vector and 2 μl Lipofectamine-2000 (Invitrogen) were incubated at room temperature for 20 min and then applied on top of cells immediately after seeding (300,000 cells/well of a 6 well plate).

Antibiotic selection was started 24 h after transfection and kept for at least 5 passages, with 150 μg/ml hygromycin-B, 1 μg/ml puromycin, 20 μg/ml blasticidin or 100 μg/ml zeocin.

Clonal transgenic lines were obtained by either manual picking of single clones upon seeding cells at a low density (5,000 cells in a 10 cm dish) or by sorting single cells into 96-well plates.

To recapitulate the endogenous expression levels of *Pax6* and *FoxA2* observed in WT RA-treated NTOs, we modulated transfection ratios and screened clones for appropriate transgene induction levels. For the generation of the 2xTetON clonal line, a ratio of 1:5 between FoxA2 and Pax6 transgene concentration was chosen.

Induction of transgene expression was achieved by supplementing the media with 1 ug/ml Doxycycline (Sigma-Aldrich). Timing and duration of induction varied according to the experiment.

#### Generation of knockout cell lines

Knockout lines were generated using CRISPR-Cas9 genome editing approach.

The FoxA2 KO line was generated using a dual-gRNA approach to excise the entire genomic locus. Both gRNAs, one binding at the 3′ region (TAGGCCTGGAGTACACTCCTTGG) and the other at the 5′ region (GCACTCGGCTTCCAGTATGCTGG), were cloned into an mU6–gRNA–hU6–gRNA–RFP vector. This construct was co-electroporated with a Cas9–GFP vector into HM1 embryonic stem cells using the Amaxa 4D-Nucleofector. 40 hours after electroporation, double-positive (EGFP⁺/RFP⁺) cells were sorted as single cells into gelatin-coated 96-well plates containing N2B27 + 2iLIF + ROCK inhibitor (Ri), and after three days, the medium was replaced with N2B27 + 2iLIF only, as Ri was included solely to support single-cell survival during the initial post-sorting period. Cell colonies were clonally expanded.

To identify complete KOs, individual clones were subjected to 2D differentiation on 2% Matrigel-coated plates in N2B27, with a RA pulse on day 2. Cells were subsequently stained for FOXA2, and sequencing was performed to confirm disruption of the locus.

The Pax6 KO line was generated using the same dual-gRNA CRISPR workflow using gRNAs (GGCCAGTACTGAGACATGTCAGG, 3’) and (GTGGTGTCTTTGTCAACGGGCGG, 5’).

### Formation of neural tube organoids (NTOs)

#### Clonal NTOs

Cells were dissociated into single cells with Accutase as described above and were centrifuged in advanced N2B27 medium (obtained by substituting DMEM/F-12 with Advanced DMEM/F-12 (Gibco) and additionally supplementing with 1% v/v non-essential amino acids (Gibco)).

After removing the supernatant, subsequent handling of cells was performed on ice. The pellet was resuspended in Matrigel (Corning) at a density of 100 cells/μl and seeded in a drop at the centre of the well, distributing the suspension evenly, without touching the border. Drop size was depending on the plate used: 4 μl for 96-well, 40 μl for 12-well and 120 μl for 6-well. Gellification was performed at 37C for 4 min (4 μl drop), 8 min (40 μl drop) or 10 min (120 μl drop).

Embedded cells were then covered with Advanced N2B27 and medium was exchanged daily starting from day 2.

For standard patterning of neural tube organoids, 250 nM all-trans retinoic acid (Sigma) was supplemented to the medium at day 2 for 18 h.

For perturbation experiments, molecules were supplemented in the media at the indicated concentrations: retinoic acid (Sigma), BMP4 (Gibco), LDN (Selleck Chemicals), PD0325901 (Tocris), FGF-2 (Fisher Scientific).

#### Chimeric NTOs

ESC cultures of different genotypes were dissociated into single cells, counted and mixed together in 1:1 ratio. Mixtures were incubated in suspension above non-adhesive 2% low-melt agarose set in 6-well plates with meniscuses to provide concave surfaces, at 37C for 2 hours in advanced N2B27, to allow them to form small aggregates. Heterotypic aggregates containing at least one cell of each type were enriched by FACS using a large nozzle, then seeded in Matrigel following normal procedures for NTO generation.

### Immunofluorescence

#### Wholemount staining of NTOs

Organoids were fixed at room temperature in 2% PFA for 30 min, by adding 1:1 volume of 4% PFA to the cultures. Fixative was then removed and samples were washed twice in PBS and blocked and permeabilized overnight at 4C in B/P solution (PBS with 1% BSA and 0.5% Triton X-100).

Antibody stainings were done in B/P at 4C for at least 24 h and samples were washed twice with PBS between primary and secondary antibodies.

Primary antibodies used were: FoxA2-goat (Novus Biologicals, #AF2400; 1:400), Pax6-rabbit (Cell Signaling Technology, #60433; 1:800), Sox1-rabbit (Cell Signaling Technology, #4194; 1:400), Shh-rabbit (Cell Signaling Technology, #2207S; 1:400), Nkx6.1-mouse (Developmental Studies Hybridoma Bank, #F55A10-s; 1:100), Nkx2.2-mouse (Developmental Studies Hybridoma Bank, 1:25), Olig2 (). For Shh staining, antigen retrieval was performed before blocking and permeabilization with 1x citrate buffer (DAKO) at 65C for 45 min.

Secondary antibodies used were: Alexa Fluor (Invitrogen Donkey anti-mouse/goat/rabbit; 1:400) and DAPI (Sigma; 1:400).

#### Optical clearing

To reduce light scattering and match the refractive index during imaging, organoids were cleared with CUBIC-R+(N) (Matsumoto et al., 2019). Samples were first adapted in CUBIC-R+(N) for at least 30 min then and then imaged in CUBIC-R+(N) supplemented with 2 mg/mL propyl gallate (Sigma) to avoid photobleaching.

### Imaging

#### Spinning disk confocal imaging

Organoids were imaged at the Olympus Spinning Disk Confocal microscope, using a 10x air (0.4 NA) or 40x silicon oil (1.25 NA) objectives. Exposure time was set at 200 ms and power was set at 100% for all lasers.

Step size in z was 2 μm for 10x and 0.57 μm for 40x acquisition.

#### Image processing and quantification

Firstly, the individual neural tube organoid was segmented in 3D using CellProfiler (Carpenter et al., 2006) based on either the DAPI channel if there is DAPI staining, or signals pooled from all channels if there is no DAPI staining. Secondly, volume parameter was used to exclude organoids that are too small or too large, likely due to nonoptimal water-shedding for touching organoids. To quantify the percentages of FOXA2^+^, PAX6^+^ or double positive cells, or percentages of cells with different genotypes (e.g. WT, FoxA2- , Pax6-, FoxA2-TetOn, Pax6-TetOn or 2xTetOn) at each organoid, we applied a voxel-based analysis, in which we assumed the voxel ratios are proxy of cell ratios in each cyst. In particular, we used the Otsu threshold method in scikit-image (v0.22.0) and python (v3.9.19) to determine if specific genotype is positive in voxel and Mean threshold for FOXA2^+^ and PAX6^+^ voxels. Finally, the results were visualized with ggplot2 (v3.4.4) in R (v4.2.1).

### Embryonic NT imaging and analysis

Mouse embryos were harvested from wildtype Bl6 crosses. All animal experiments were performed in accordance with Austrian animal welfare legislation and conducted under institutional facility approval and oversight. C57BL/6J mice were bred and housed at the Research Institute of Molecular Pathology (IMP)/Institute of Molecular Biotechnology (IMBA) shared animal facility under standard specific pathogen–free (SPF) conditions at a room temperature of 22°C and 55% humidity on a 14-hour light/10-hour dark cycle and unrestricted access to food and water.

We performed an immunofluorescence timecourse on embryos harvested from E6.5-9.5, which we staged developmentally according to morphological hallmarks and somite number. We stained for FOXA2, PAX6, SOX2 and DAPI, cleared embryos in CubicR+(N), mounted in 2% low-melt agarose in CubicR+(N), and acquired wholemount images on Olympus Spinning Disk with 30x silicon immersion objective. As expected, PAX6 expression commenced in embryos from the 4-somite stage onwards. PAX6 expression was observed in the forebrain, in the forming hindbrain alongside somites 1-4, but not in the midbrain, as expected. FOXA2 expression was observed, as expected, in the primitive streak, extraembryonic and definitive endoderm, and the neural plate midline. In E8.0-8.5 embryos that contained PAX6 expression, we scored whether FOXA2^+^PAX6^+^ double-positive cells were present in different anatomical regions of embryos. FOXA2^+^PAX6^+^ double-positive cells were observed in the forming forebrain and hindbrain, and also in more posterior regions of the neural plate/tube as the spinal cord forms from somite 5 onwards. See Table S2 for full scoring results.

#### Cell segmentation with CellPose

For 3D-Segmentation of SOX2^+^ nuclei in embryos we used CellposeSAM. To enhance processing speed we downsampled the datasets in xy. A gaussian blur of 2 provided noise reduction. To improve the segmentation results of the predefined model we retrained it using 2D slices in all 3 directions from our own datasets. We applied this model and saved the resulting labels which were then cleaned up in Fiji via a semiautomated macro. SOX2^+^ cells that did not belong to the spinal cord were removed (e.g. surface ectoderm). The cleaned-up label image was then resized to the original and used for cellwise intensity analysis on the original data.

#### Feature quantification downstream of segmentation

With those cleaned masks, features, including the cell volumes, Wadell sphericity and intensity of FoxA2, Pax6 and Sox2, were collected and quantified from the raw images using scikit-image (v0.22.0) and python (v3.9.19). The collected cells were further filtered based on the size and sphericity. Finally, the intensity of those three marker genes were visualized with scatterplot in R (v4.2.1).

### Flow cytometry

#### Sample collection and nuclear staining

Protocol was adapted from (Delás et al., 2023). Specifically, organoids were dissociated into single cells by manually spraying and resuspended them with Accutase (1 mL for a 6-well), thereby breaking the Matrigel drop. They were initially incubated at 37C for 5 min, manually dissociated a second time and incubated again at 37C for an additional 10 min. Organoids were then again manually dissociated until a single cell suspension was achieved and the enzyme was inactivated using an equal volume of ice-cold PBS containing 0.5% BSA. Cells were then handled on ice for all further steps. Cells were harvested and centrifuged at 4C for 4 min at 400g and pellets were resuspended in 1 μl/mL LIVE/DEADTM Fixable Dead Cell Stain Near-IR fluorescent reactive dye (ThermoScientific) in PBS. They were incubated on ice, protected from light, for 30 min and centrifuged at 4C for 4 min at 400g. Cells were then fixed in 100 μl 4% paraformaldehyde (PFA) for 10 min at room temperature, washed with 1 mL PBS and centrifuged at 4C for 5 min at 2000g.

Pellets were resuspended in 200 μl PBS with 0.5% BSA and either stored at 4C or processed for antibody staining.

Cells were counted and normalized for the lowest number and centrifuged at 4C for 5 min at 2000g in 96 V-bottom well-plates. Pellets were resuspended in 100 μl of PBS with 0.5% BSA and 0.1% Triton-X100 containing the conjugated antibody mix.

Cells were incubated overnight at 4C and washed twice the next day with 100 μl of PBS with 0.5% BSA and 0.1% Triton-X100, at 4C for 5 min at 2000g.

Pellets were resuspended with 120 μl of PBS with 0.5% BSA and analysed at the NovoCyte Penteon (Agilent).

All antibodies used were diluted 1:100: Sox2-V450 (BD Biosciences), Pax6-PerCPCy5.5 (BD Biosciences), Pax6-PE (BD Biosciences), Nkx6.1-A647 (BD Biosciences), FoxA2-AF488 (BD Biosciences), Nkx2.2-PE (BD Biosciences), Oct4-PerCPCy5.5 (BD Biosciences), Sox1-A647 (BD Biosciences).

Compensation was done using beads (ThermoFisher Scientific, 01-2222-42) incubated with the same antibodies.

#### Data pre-processing

Data was processed and analysed using FlowJo software. FlowClean plugin was used as quality control to identify and exclude potential abnormal events during acquisition. Default settings, including all uncompensated fluorescent channels as well as time parameters were used. Only the “GOOD” events were then exported and further analysed.

The following gating strategy was applied to all samples:

- Selection of population using FSC-H vs SSC-H
- Selection of single cells using FSC-H vs FSC-A
- Identification of live cells using FSC-H vs Live-dead dye -H
- Selection of neural progenitors (Sox2^+^ cells) using FSC-H vs Sox2-H

#### Clustering and visualization

The flow cytometry data of five markers (FoxA2, Oct4, Pax6, Sox1 and Sox2) from all time points were used to identify five clusters using R package mclust (v5.4.7), as previously shown (Saez et al., 2022). To visualize the flow cytometry data in a low-dimensional embedding, we converted our matrix to SingleCellExperiment object (v1.16.0) (Amezquita et al., 2020) and computed diffusion map with CATALYST (v1.18.1) (Crowell et al., 2022) by specifying the parameters of the cosine distance and local sigma.

### RNA sequencing

#### Sample collection

NTOs were harvested at each timepoint, dissociated into single cells, then frozen at 1 million cells/mL in Cryostor CS10 (Sigma-Aldrich). After all timepoints were collected, samples were thawed, stained with Draq7, enriched for viable cells (Draq7-) by FACS, then processed for 10x scRNA-seq library preparation.

#### Processing of scRNA-seq data

The raw data of scRNA-seq samples from WT NTOs were aligned and quantified with the mouse genome and annotation from 10x, mm10_10x, using CellRanger (v7.0.0) with default parameters and excluding the intronic reads. With the barcode counts, empty drops were determined and filtered with DropletUtils (v1.14.2) (Lun et al., 2019). The filtered gene count matrix (cell barcodes* genes) were imported into Seurat (v4.3.0) (Hao et al., 2024) as Seurat Object (v4.1.3). The cells were further filtered if they have <1000 or >10k detected genes or percentage of mitochondrial gene expression larger than 5% of total UMIs. Then for each scRNA-seq sample, doublets were identified with R package DoubletFinder (v2.0.3) (McGinnis et al., 2019) and discarded for the downstream analysis. After mitochondrial and ribosomal genes were filtered, the count matrices were normalized in Seurat with the “LogNormalize” method and the top highly variable features were calculated with the “vst” method. To mitigate the technical variability between scRNA-seq samples, the count numbers were regressed out in the data scaling. Moreover, the cell-cycle scores and phases were computed in Seurat using the known G2/M and S phase markers. The UMAP embeddings were calculated in Seurat with 30 PCs calculated from 5000 highly variable genes.

#### scRNA-seq pseudo-time, cellular heterogeneity and clustering

To facilitate the pseudo-time calculation, scRNA-seq data of before (day2), with and without RA-treatment (from day2.5 to day5) were selected. To alleviate the confounding effect of cell cycle, the genes that have high correlation with cell phases (i.e. >5% of expression variance can be explained by cell cycle phase) were not considered in the highly variable gene searching. 30 PCs were calculated using 3000 variable features in Seurat with parameter weight.by.var = FALSE.

Those PCs were used for Palantir (v1.3.0) (Setty et al., 2019) to compute the diffusion-map dimension reduction. By specifying the start cell from before RA (day2) sample and cells in terminal states (from day5 for RA- and noRA-treated samples), we obtained the pseudo-time for each cell in scRNA-seq data. To compare the cellular heterogeneity between RA- and noRA-treatment, the pseudo-time from Palantir was first regrouped into 10 bins. For each bin, 200 cells were then randomly sampled for the sake of computation. Top 20 PCs were calculated with scater (v1.22.0) (McCarthy et al., 2017) and cell heterogeneity in each bin were quantified as Euclidean distance distribution in the PC space as in (Mohammed et al., 2017).

To cluster the scRNA-seq data of RA-treated samples from day2.5 to day5, the genes highly correlated with cell cycle were first discarded and top 30 PCs were calculated using top 2000 variable genes in Seurat with parameter weight.by.var = FALSE. SLM algorithm was applied with resolution = 0.7 and small clusters were manually merged to have 7 clusters. Cluster-specific marker genes were found with Seurat function FindAllMarkers.

#### Sparse feature selection of RA-treated scRNA-seq data

To identify a minimal list of genes recapitulating low dimensional landscape in the transcriptomic space, we have performed three sparse feature selection methods, which were implemented with different assumptions. Moreover, only annotated transcriptional factors (TFs) and genes involved in signalling pathway were considered as the features of interest. Top 50 features were selected with GeneBasis (Missarova et al., 2021), 74 features from DUBStepR (Ranjan et al., 2021) and 31 features from SMD (Melton and Ramanathan, 2021). Many features were consistently predicted by all three methods (see Table S1), e.g. FoxA2, Pax6, Shh and Cdh1.

#### Predict signalling pathways for regulative cell proportions with scRNA-seq data

To predict the key signalling pathways that regulate cell proportions of FP and NP, we set out with previously identified clusters in scRNA-seq of RA-treated samples from day2.5 to day5 and applied CellChat (v2.1.2) (Jin et al., 2025) with mouse database in CellPhoneDB (Efremova et al., 2020). To calculate the cellular communication network, computeCommuProb function was used with type = “triMean”. Signal pathways that contribute to both outgoing and incoming signalling across clusters were shown for BMP, WNT, FGF and sonic Hedgehog (HH) in Fig S4A.

To further identify specific ligand-receptor pairs involved in the cell-cell communication, we considered the clusters 3 and 5 as sender or receiver subpopulation to apply LIANA (v0.1.13) (Dimitrov et al., 2022), in which ligand-receptor database from CellPhoneDB and scoring methods, “natmi”, “connectome”, “logfc”, “sca”, were selected. The outputs from different methods were aggregated and the top 100 ligand-receptor pairs were visualized in Fig S4B using a customized Circos plot modified from Connectome (v1.0.1) (Raredon et al., 2019).

#### Analysis of published mouse embryo scRNA-seq dataset

To investigate the relevance of FoxA2-Pax6 co-expressing cell states, we downloaded processed scRNA-seq data of 36 mouse embryo samples from MouseGastrulationData (v1.8.0) (Pijuan-Sala et al., 2019). The original low-dimensional embeddings (i.e. corrected PC and UMAP) were retained. Moreover, to simplify our analysis, we discarded some irrelevant annotated cell clusters, e.g. Erythroid, Blood, Allantois, mesoderm, Haemato, Cardiomy, Endothelium, Mesenchyme, ExE.

