## Supplemental Figures for "Minimal essential requirements for neural tube self-organisation"

### SUPPLEMENTARY FIGURE 1

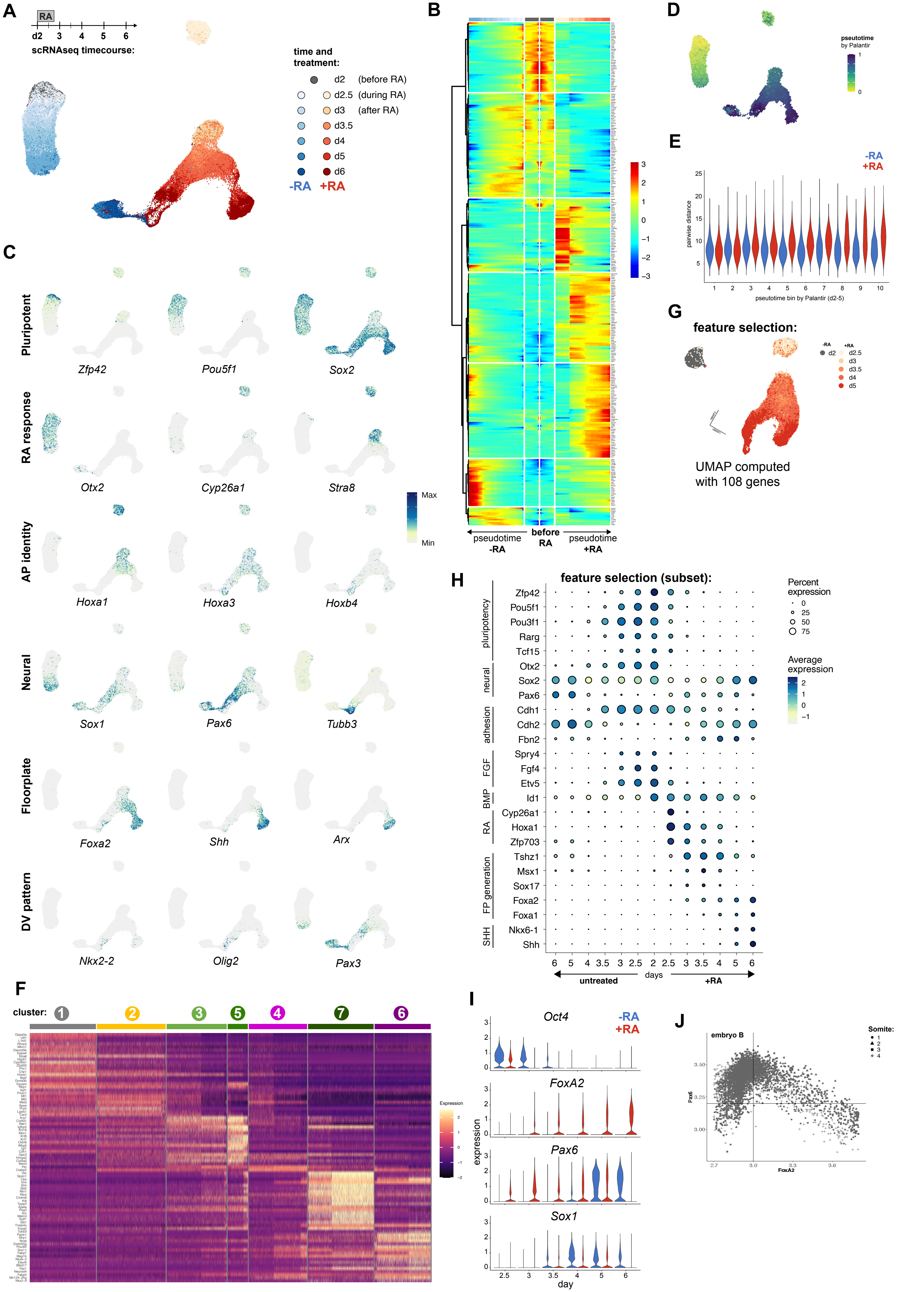

### SUPPLEMENTARY FIGURE 2

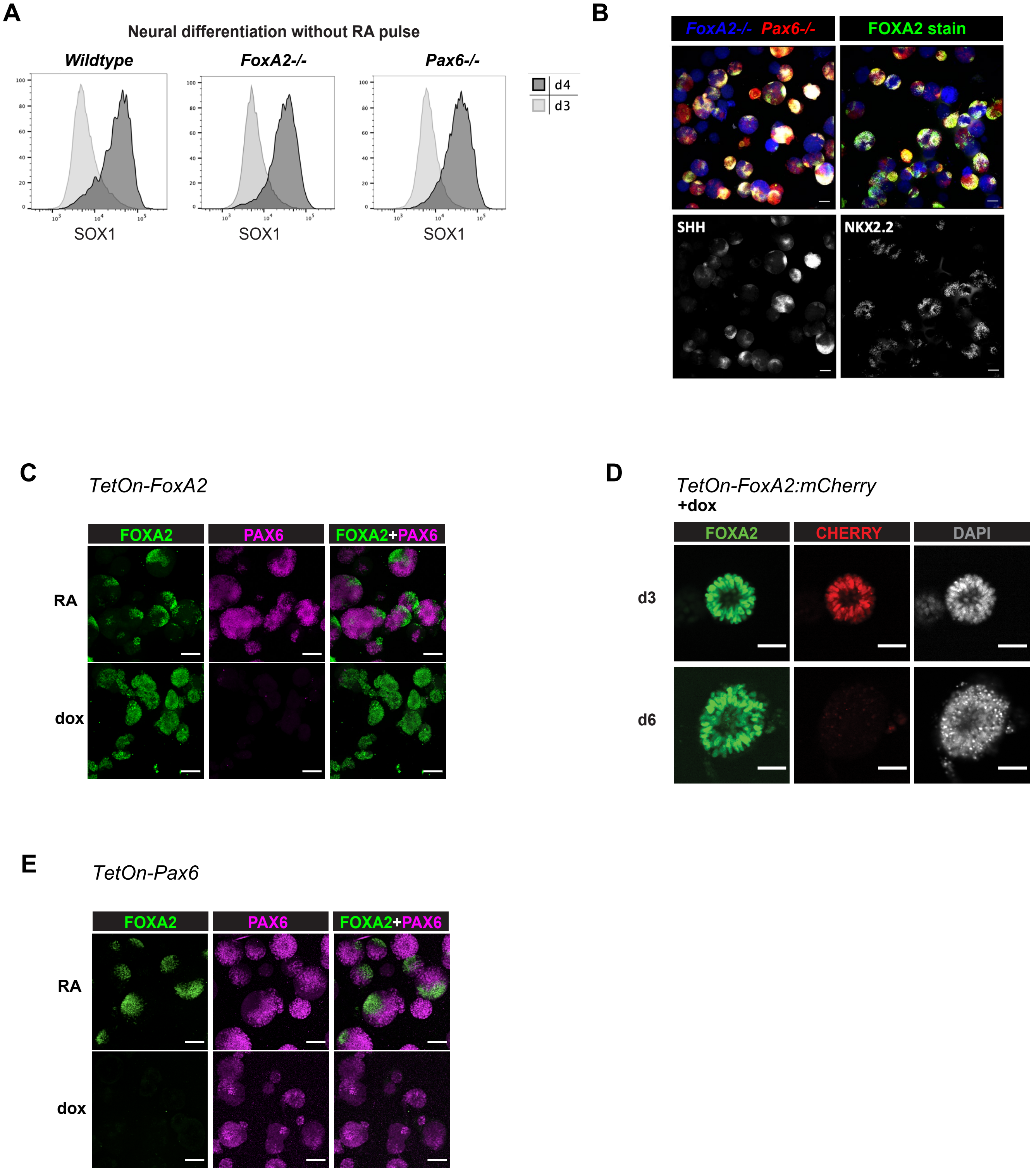

### SUPPLEMENTARY FIGURE 3

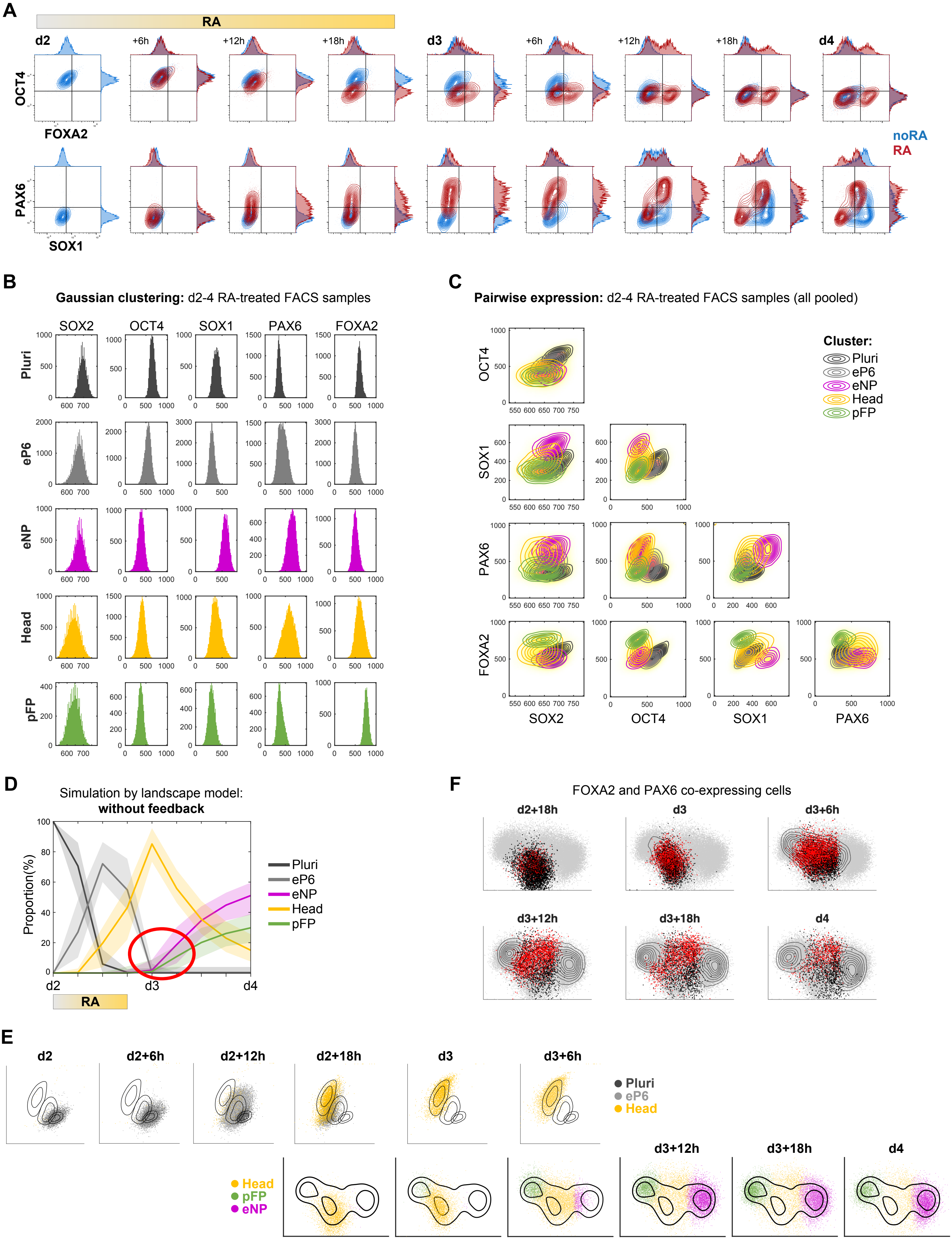

### SUPPLEMENTARY FIGURE 4

A

B

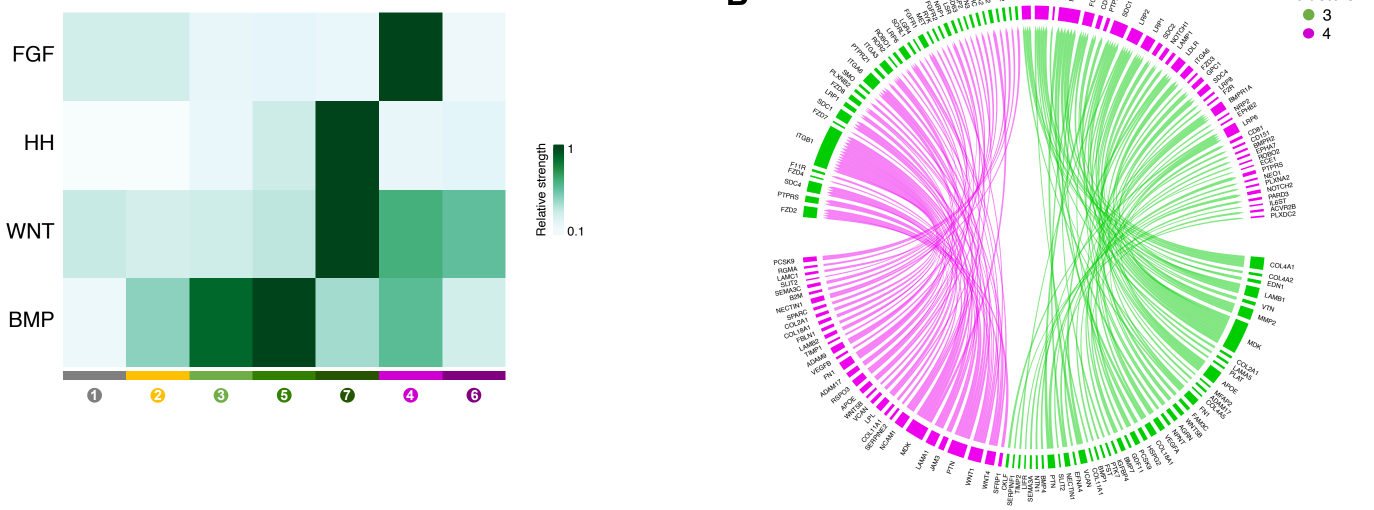

### SUPPLEMENTARY FIGURE 5

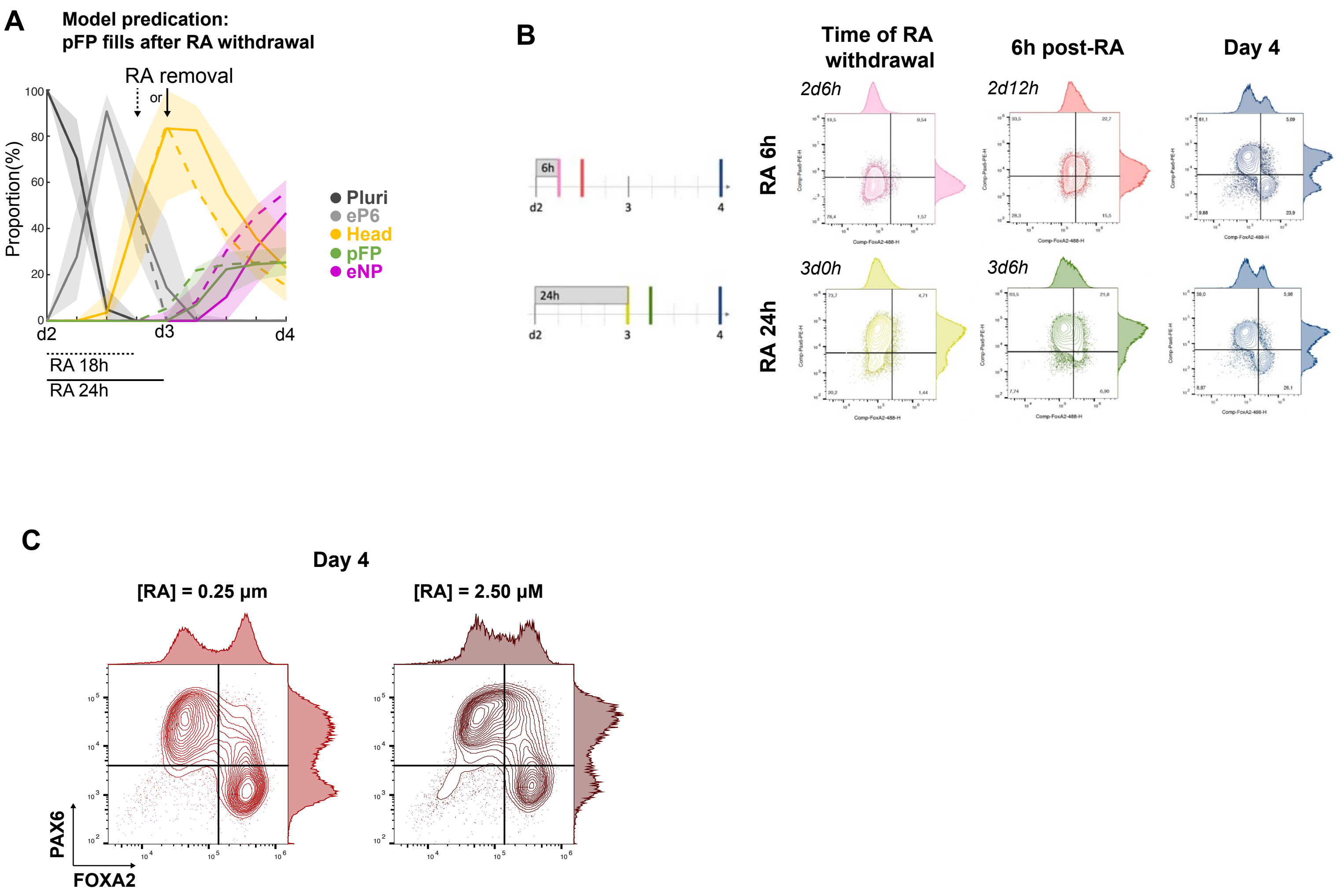
