## Supplementary material for "Minimal essential requirements for neural tube self-organisation": Supplmental Information

### Supplementary Information

May 21, 2026

### 1 Model background

Theoretical models have provided inspiration and insight into the process of cellular differentiation. The Waddington landscape, which conceptualised cell differentiation as a ball rolling down a landscape of branching valleys representing cell fates, is a popular metaphor that incorporates the idea of developmental plasticity and robustness into the process of cell fate decision making ([1]). This metaphor can be mathematically formalised using dynamical systems theory. The resulting dynamical landscape models provide a quantitative and predictive framework for studying cell fate allocation ([2]).

In these models, the dynamics of decision making are represented by a vector field determined by the landscape. Waddington's valleys, the cell fates, correspond to attractors, while saddles separate these attractors and act as decision points in the landscape. The trajectories of cells moving through the landscapes are defined by the unstable manifolds that connect saddles to attractors, creating pathways through the cell state space. Together, the saddles, attractors and unstable manifolds define the topological structure of the landscape and determine how cells move within this cell state space ([2]).

Varying the parameters to represent changes in signalling alters the landscape and therefore the vector field, thereby changing how cells move through the state space. As an attractor approaches a saddle, it becomes shallow, allowing cells to leak out of the attractor basin in response to stochastic fluctuations. Cell fate transitions occur when cells enter new basins of attraction, either through signal-induced bifurcations or stochastic fluctuations. When an attractor and saddle collide, both disappear in what is known as a bifurcation (or in reverse a saddle and attractor are created) fundamentally reshaping the cellular decision-making landscape.

The transitions observed in developmental systems are typically between relatively small numbers of possible outcomes and for many systems we are essentially interested in the transitions between stationary states in a dynamical system. Thus we can expect that although the regulatory network is complex, the resulting dynamics are relatively simple ([3]).

We are interested not just in a single dynamical system but a parameterised family where the parameters $\theta$ can be understood as functions of the signals that the cell is receiving. We call these parameterised landscapes (PLs). The cellular transitions are described by bifurcations of the PL. The bifurcation set divides the parameter space into components consisting of topologically equivalent quasi-gradient Morse-Smale systems. We call these MS-components. These are separated by components of the bifurcation set that in general have a stratified structure where codimension 1 hypersurfaces join together in sets of higher codimension. In the case of 2 and 3 parameters, which we consider here, the hypersurfaces are curves and surfaces respectively [3]. A combination of catastrophe theory and dynamical systems approaches (e.g. [4, 5]) allow the characterisation of the local structure of these bifurcations subject to a mild and precise genericity assumption, if the number of parameters is no greater than 5. For example, in 2 parameter PLs generically the bifurcation set will consist of curves of fold points with a finite number of cusps on them. Not only does the theory provide a qualitative description of the local behaviour but it also provides a normal form ([6]). The normal form will be related to the true model and this opens up the possibility of fitting the biological data to the normal form because summary statistics of the true model and the normal form should be the same. Therefore, if we fit the normal form to the experimental data, then we have determined the qualitative form of the true model and those quantitative aspects of the true model that are preserved.

### 2 Clustering

The clustering method we used is based on the hypothesis that cell identities are attractors of a dynamical system, and therefore gene expression in a population of cells with a particular identity should form an approximately multivariate normal distribution around the attractor because close to the attractor gene expression dynamics will be essentially linear. The method is based on the one used in [5].

Given the data coming from a particular experiment, we pool all the experimental data from all replicas and all time points into a single dataset and fit a Gaussian mixture model (GMM) to it using the MATLAB algorithm `fitgmdist`. A GMM $Y$ consists of a weighted sum of $m$ multivariate normal distributions (MVNs). These MVNs have probability distributions $P_i(x)$, $i = 1, ..., m$ defined by their means $\mu_i$ and covariance matrices $\Sigma_i$. The clusters are in 1-1 correspondence with the MVNs in the GMM and we regard a cell as belonging to one of the $m$ clusters if its 5-dimensional state $x$ has probability greater than some threshold probability $p$ for the corresponding MVN i.e. $P_i(x) > p$. In cases where a cell could belong to more than one cluster it was allocated to the one where it had the highest probability.

The number of components of the GMM is chosen manually by inspection of the distributions in order to guarantee that each cluster has a unimodal distribution. Figure A1 shows the normality of the clusters obtained. We then analysed the protein levels in each cluster and used this to label it with the cell identity it best represented (Pluri, eP6, Head, pFP, eNP). The observation that the clusters obtained by this automatic algorithm are consistent with the already known cell identities provides a validation of the method.

Once the cells had been allocated to clusters, we computed the population proportions at each time point. The proportions for each of the replicas in the RA treated experimental dataset corresponding to each cluster can be found in table A1. These proportions are the data used to construct and parameterise the landscape model.

| | Pl | e6 | eN | H | FP | Pl | e6 | eN | H | FP | Pl | e6 | eN | H | FP |
|---|---|---|---|---|---|---|---|---|---|---|---|---|---|---|---|
| d2 | 92 | 7 | 0 | 1 | 0 | 94 | 6 | 0 | 1 | 0 | – | – | – | – | – |
| d2+6h | 72 | 27 | 0 | 1 | 0 | 73 | 26 | 0 | 1 | 0 | 70 | 29 | 0 | 0 | 0 |
| d2+12h | 3 | 96 | 0 | 1 | 0 | 10 | 88 | 0 | 1 | 0 | 7 | 91 | 0 | 2 | 0 |
| d2+18h | 1 | 41 | 1 | 57 | 0 | 0 | 57 | 0 | 42 | 0 | 0 | 52 | 0 | 47 | 0 |
| d3 | – | – | – | – | – | 0 | 2 | 0 | 90 | 8 | 0 | 1 | 0 | 89 | 10 |
| d3+6h | 0 | 0 | 10 | 71 | 18 | 0 | 0 | 10 | 70 | 20 | 0 | 0 | 7 | 69 | 24 |
| d3+12h | 0 | 0 | 47 | 31 | 22 | 0 | 0 | 42 | 33 | 25 | 0 | 1 | 42 | 32 | 26 |
| d3+18h | 0 | 0 | 48 | 23 | 28 | 0 | 1 | 49 | 24 | 27 | 0 | 0 | 47 | 24 | 29 |
| d4 | 0 | 0 | 50 | 23 | 27 | 0 | 0 | 51 | 22 | 27 | – | – | – | – | – |

Table A1: Proportions obtained by clustering the RA treated samples. Dash means that data was not available for that replica at that particular time point. Fates have been abbreviated for compactness (Pl=Pluri, e6=eP6, eN=eNP, H=Head, FP=pFP).

The observation that protein level distributions in the Head population were unimodal but with higher variance (Supp. Fig. 3BC) than other clusters, together with the heavier tails of the qq-plots for this cluster prompted us to further investigate its structure. The LDA representation of clusters eNP, pFP and Head in Fig. S3E show that the cells labeled as Head correspond to the cells in the attractor state together with some of the cells in transition towards eNP and pFP. Figure A2 shows the movement of cells through the regions corresponding to Head, pFP and eNP. We observe how some of the Head region corresponds to cells transitioning first towards pFP and later towards eNP. We modelled the Head state as an attractor. The structure of the data suggest that the Head attractor is shallow and the cluster may be formed by cells in the Head attractor together with cells already in transition towards pFP and ENP but not yet committed to these later states. This simplification does not affect the purpose of the model, that is, understanding the symmetry breaking mechanism.

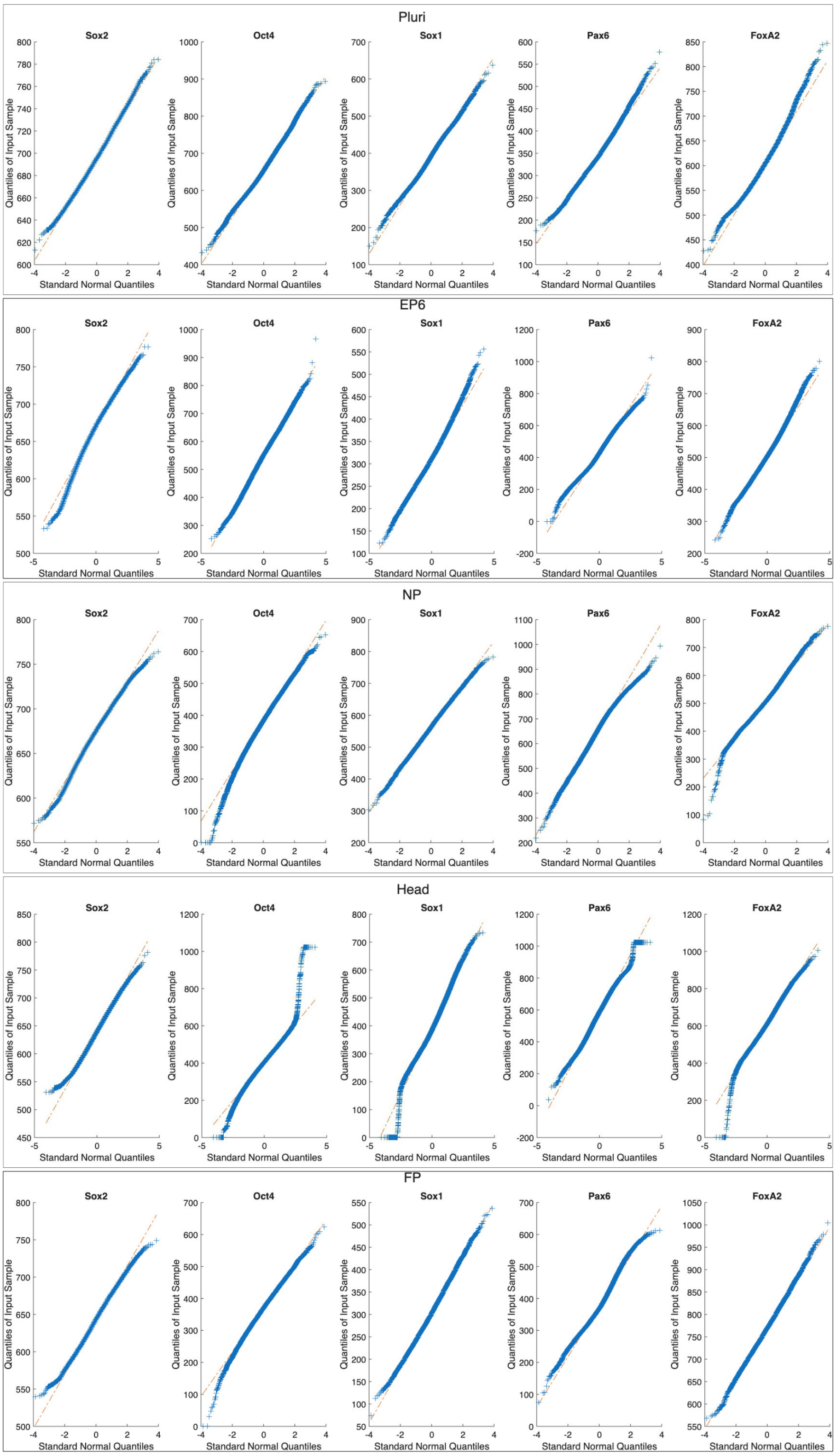

Figure A1: Q-q plots for the clusters obtained by Gaussian mixture model clustering of the original dataset.

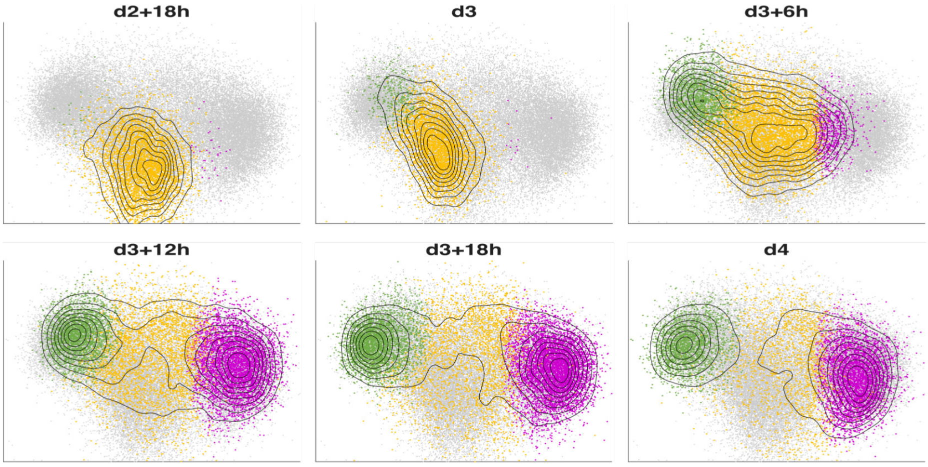

Figure A2: **LDA projections for Head, pFP and eNP clusters**. Coloured areas correspond to the cluster regions with colors corresponding to cell identity. Level curves show the distribution of cells at each time point. Grey background shows the complete cell population at all time points.

### 3 Initial modelling

#### 3.1 Model topology

The structure of gradient dynamical landscapes can be described by a limited number of universal normal forms [7] and Thom's catastrophe theory offers a classification scheme for these [8]. In the case of three attractors - a progenitor cell choosing between two alternative progeny fates - there are two principal arrangements of attractors resulting in two forms of decision: a "binary choice", in which a progenitor attractor is separated from each fate by a distinct saddle; and a "binary flip," in which the progenitor attractor is separated by a single saddle from both fates and the choice a transitioning cell makes depends on the unstable manifold of the saddle [2]. This latter mechanism provides a means for regulating the relative proportions of two fates generated by a progenitor population.

The Pluri, eP6 and Head attractors correspond to a binary choice topology, because the transition between them is ordered and sequential. Albeit there is no choice here, the topology of the binary choice corresponds to an ordered transition between attractors when cells start at the top-most attractor. By contrast, two key observations suggested that a binary flip topology is more appropriate for the arrangement of Head, pFP and eNP. First, both the scRNAseq analyses (Fig. 1) and the LDA analyses of the FACS data (Supp. Fig. 3 and Fig. A3) suggested a single trajectory emanating from the Head attractor that subsequently diverged into separate pFP and eNP trajectories, resulting in the characteristic inverted Y shape evident in the data projection. Second, the pFP and eNP attractors fill simultaneously but accumulate different cell numbers (with an apparent upper bound for pFP). This supports the assignment of a flip topology as stochasticity would allow cells leaving the Head attractor to distribute in different ratios between pFP and eNP attractors in parallel.

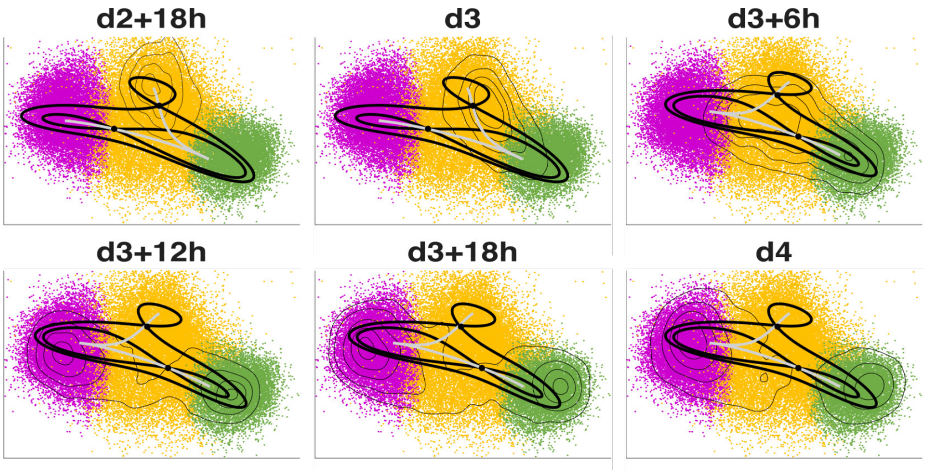

Figure A3: **Landscape structure on LDA projection** Coloured areas correspond to the cluster regions with colors corresponding to cell identity. Thin level curves show the distribution of cells at each time point. The hypothesized landscape structure is overlapped.

We used gradient dynamics defined by polynomial functions to define the landscape dynamics. We used normal forms for the two PLs ((1), (2)). These minimise the number of parameters and state variables needed to capture the changing geometries. We took subfamilies of the double cusp catastrophe ($x^4+y^4$), which contains both PLs mentioned earlier [7], to guarantee that the systems obtained were compact. That is, no trajectories escape to infinity. In general, a Riemannian metric is necessary to incorporate all the dynamical properties of quasi-gradient Morse-Smale systems (see [3]) but was not necessary here, which kept the number of parameters of the model minimal.

The choice (Pluri-eP6-Head) and flip (Head-pFP-eNP) landscapes were connected through the common Head attractor using a function that smoothly transitions a cell's trajectory from one landscape to the other. In particular, we used $\chi(x, y) = \tanh(10(y + 0.5))$ placing the transition around $y = -0.5$.

Since our state variables are abstract, we chose the scale of the model without loss of generality. This allowed us to fix some of the parameters in the PLs and concentrate in two dimensional subfamilies of the double cusp catastrophe. For the first decision, we used the parameterised family of potential functions

$$F_1(x, y; p) = x^4 + y^4 + x^3 - 4xy^2 + x^2 + p_1x + p_2y. \tag{1}$$

The bifurcation locus for the family $F_1$ is as shown in Figure A4A. In this family the region with three attractors is the area inside the middle diamond. Here, the central attractor is always the same and it can bifurcate with any of the two saddles. For our model only the part of the family on the bottom half was relevant, because we considered the ordered progression of cells from the top attractor toward the bottom attractor.

For the second decision we took the family of potential functions

$$F_2(x, y; p) = x^4 + y^4 + 2x^2y - y^3 - y^2 + p_3x + p_4y. \tag{2}$$

The bifurcation locus for the family $F_2$ is shown in Figure A4B. In this family, the region with three attractors corresponds to the large central triangle, excluding the smaller blue-lined triangle(blue curves with cusps) close to (0,0). Inside the small triangle a repeller in the middle of the three attractors appears as part of the landscape. The purple dotted lines denote the regions where the unstable manifold for one of the saddles flips, changing the middle attractor. For our model only the region inside and to the top of the large central triangle is relevant because we assume that neither pFP nor eNP bifurcate.

Each of the two landscapes contains an attractor labelled "Head", by which the two landscapes were "glued" together in order to build the global landscape. Hence, the ordinary differential equation (ode) used for the deterministic part of the simulations is

$$\begin{aligned}\frac{dx}{dt} =& p_5 (1 + \chi(x, y)) \frac{dF_1}{dx}(x, y) - p_6 (1 - \chi(\overline{x}, \overline{y})) \frac{dF_2}{dx}(\overline{x}, \overline{y})\\ \frac{dy}{dt} =& p_5 (1 + \chi(x, y)) \frac{dF_1}{dy}(x, y) - p_6 (1 - \chi(\overline{x}, \overline{y})) \frac{dF_2}{dy}(\overline{x}, \overline{y})\end{aligned}$$

with $\chi(x, y) = \tanh(10(y + 0.5))$ and $(\overline{x}, \overline{y}) = (x - 1, y + 2)$.

In summary, the deterministic model depends on 6 parameters with different effects on the flow of cells through the landscape as detailed in table A2.

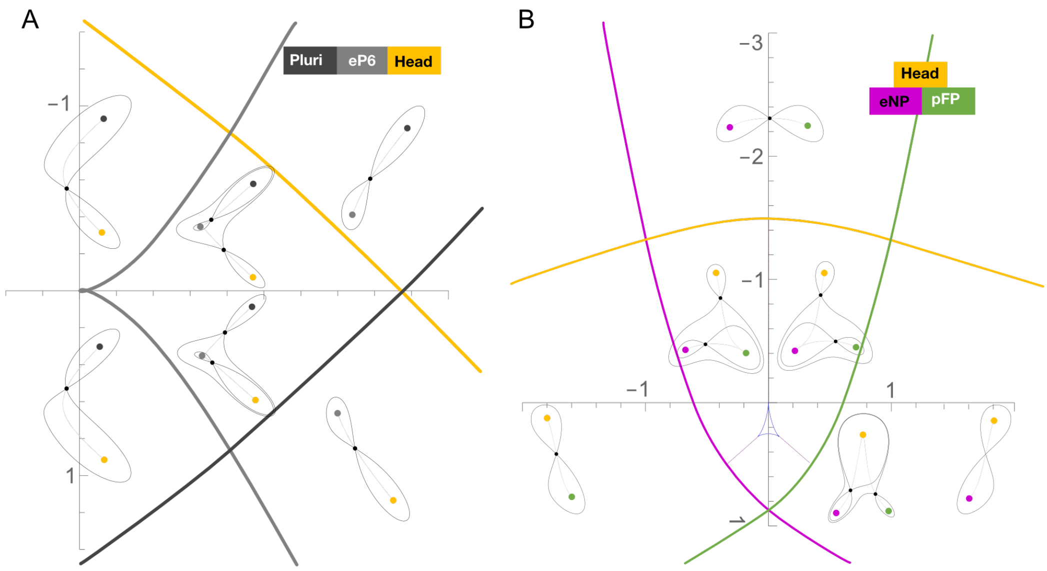

Figure A4: **Bifurcation locus** A. Bifurcation curves for the family $F_1$ corresponding to the first decision. B. Bifurcation curves for the family $F_2$ corresponding to the second decision. Continuous curves correspond to bifurcation curves and they are coloured according to the attractor that bifurcates when crossing it. Purple lines show the flip lines where the middle attractor changes. The landscape corresponding to a point on the purple line includes a saddle connection (the stable/ unstable manifold of one saddle contains the other one). Examples of landscapes in the different regions are shown.

| Parameter | Meaning |
|---|---|
| $p_1$ | topology of first decision |
| $p_2$ | topology of first decision |
| $p_3$ | topology of second decision (Head stability) |
| $p_4$ | topology of second decision (eNP/pFP allocation) |
| $p_5$ | velocity of first decision |
| $p_6$ | velocity of second decision |

Table A2: **Global model parameters** List of parameters for the global model. The first decision is the linear transition from Pluri, to eP6 to Head through a binary choice landscape. The second decision is the binary flip from Head to either eNP or pFP.

The evolution of cells with time, given a signalling regime, is then modelled with a stochastic dynamical system [9]

$$(\dot{x}, \dot{y}) = (\frac{dx}{dt}, \frac{dy}{dt}) + \sigma dW \tag{3}$$

where $dW$ is a two dimensional Wiener random process and $\sigma$ is a parameter controlling the amplitude of the noise perturbation.

#### 3.2 Model fitting

Once a dynamical system with these topological features was built, we proceeded to furnish it with parameters by fitting to the experimental data using an ABC SMC framework [10]. In the initial fit, the parameters depended only on the RA level. The simulation process is similar to the one used in [5]. To fit the model, we used the proportions of each cell type in the time course FACS data for RA-treated NTOs (Fig. 3A-G, S3A-C). The value of the parameters depend on the signalling regime, hence there was a total of 12 parameters to fit: 6 for the no-RA period and 6 for the modification caused by RA addition. Hence,

$$\vec{p} = \vec{p}_{no\_RA} + \alpha \ \vec{p}_{add\_RA}$$

where $\alpha$ is 0 when no RA is included and 1 otherwise. There is an additional parameter $\sigma$ that controls the size of the deterministic time step in the stochastic simulation.

In order to initiate the fitting one must establish priors for the different parameters of the landscape. Since the data inform the landscape with and without RA, we establish the priors for $\vec{p}_{no\_RA}$ and $\vec{p}_{RA}$ and then make the necessary transformations during the simulation process.

Table A3: Tables of priors for the parameters of the initial fitting.

| **Parameter** | **Prior** |
|---|---|
| $p_1^{no\_RA}$ | $\mathcal{U}([-1, 1])$ |
| $p_2^{no\_RA}$ | $\mathcal{U}([0, 1.5])$ |
| $p_3^{no\_RA}$ | $\mathcal{U}([-0.5, 0.5])$ |
| $p_4^{no\_RA}$ | $\mathcal{U}([-2, -1])$ |
| $p_5^{no\_RA}$ | $\mathcal{U}([0.01, 1])$ |
| $p_6^{no\_RA}$ | $\mathcal{U}([0.01, 1])$ |
| $\sigma$ | $\mathcal{U}([0, 0.2])$ |

| **Parameters** | **Prior** |
|---|---|
| $p_1^{RA}$ | $\mathcal{U}([0.5, 2])$ |
| $p_2^{RA}$ | $\mathcal{U}([0, 1.5])$ |
| $p_3^{RA}$ | $\mathcal{U}([-0.5, 0.5])$ |
| $p_4^{RA}$ | $\mathcal{U}([-2, 0])$ |
| $p_5^{RA}$ | $\mathcal{U}([0.01, 1])$ |
| $p_6^{RA}$ | $\mathcal{U}([0.01, 1])$ |

The results of the fitting algorithm after 7 rounds of fitting are shown in Figures A5, A6, A7 and A8.

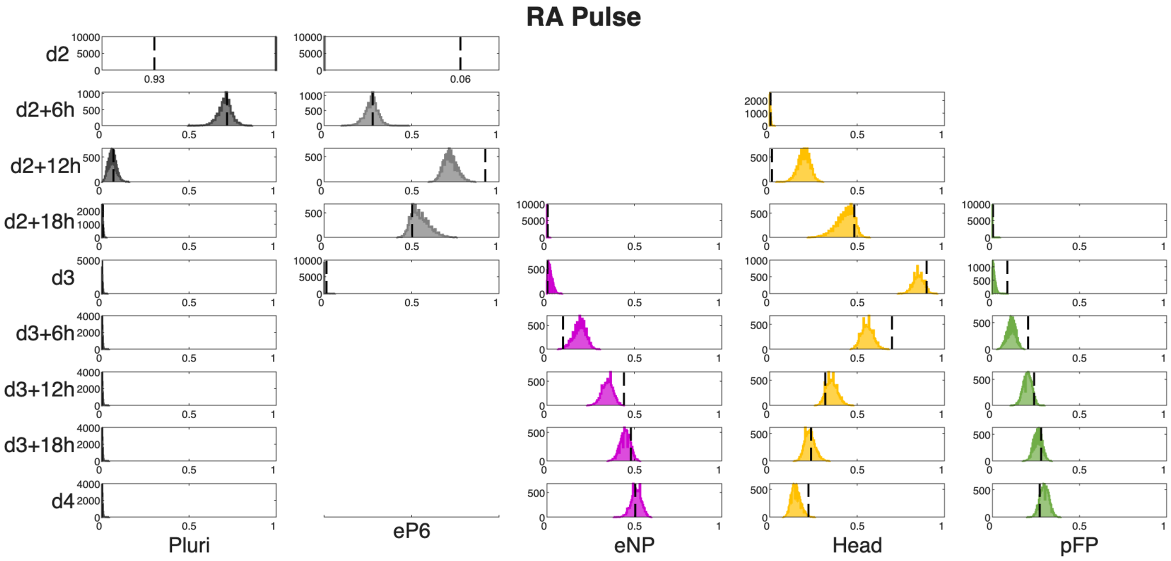

Figure A5: Histograms showing the distributions of cell states using the 10000 parameter vectors accepted in the final iteration of the initial fitting algorithm. The vertical dotted lines indicate the proportions from the experimental series used as training data. The missing panels correspond to populations with no assigned simulated cells and experimental proportion equal to zero. Related to Fig. S3D.

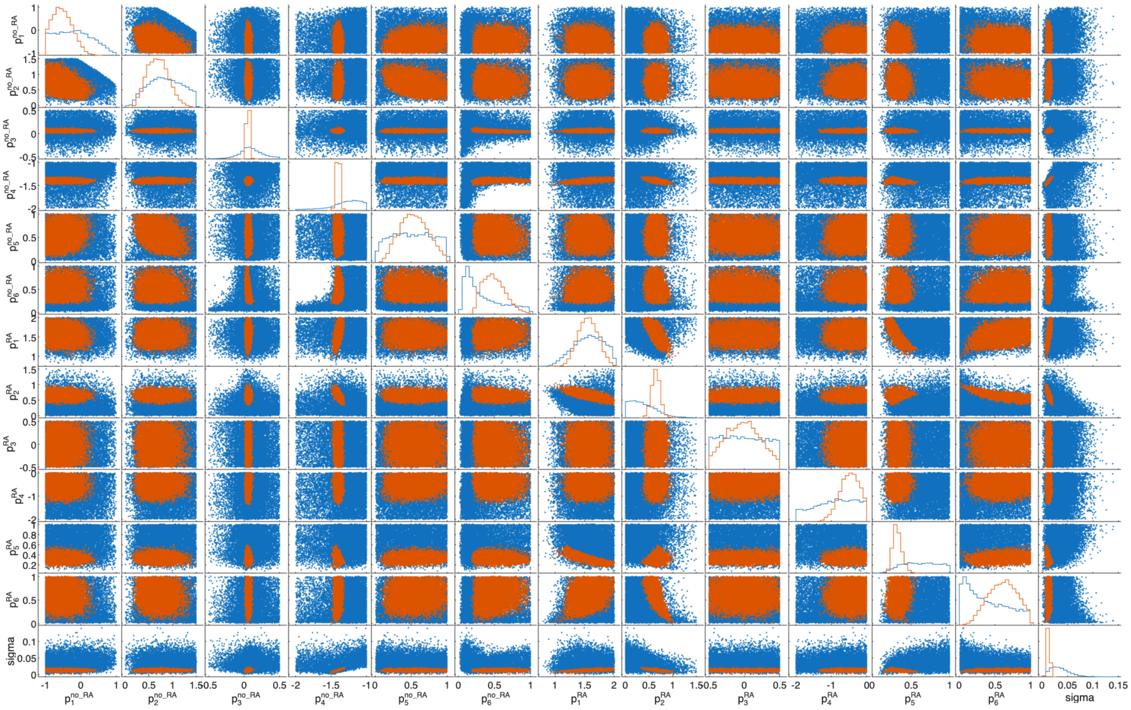

Figure A6: Distributions of accepted parameters in the first iteration (blue) compared to accepted parameters in the last iteration (red) of the initial fitting algorithm.

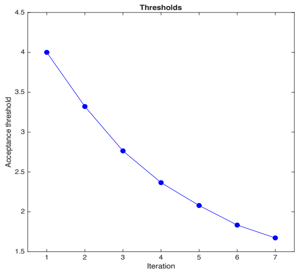

Figure A7: Evolution of acceptance threshold in the 7 iterations of the fitting for the initial model

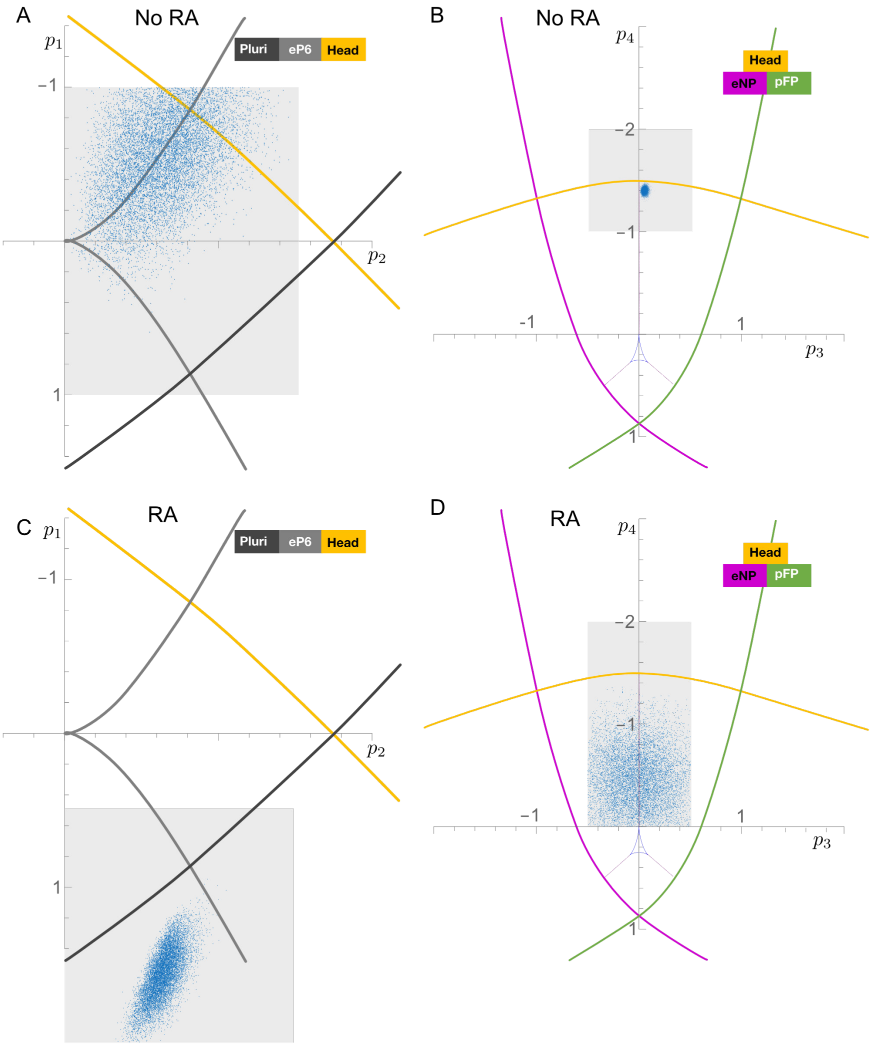

Figure A8: Priors (shaded area) and accepted parameters in the last round of fitting for the initial fitting algorithm together with the corresponding bifurcation locus.

### 4 Refined model including feedback

Following the initial model fit we added two further features: 1. A time evolution of the parameters of the binary choice landscape. 2. Feedback that modifies the parameters of the binary flip, which depends on the number of cells occupying the pFP attractor.

To this end, we added parameters to the model and defined a new parameter vector $p$ as

$$\vec{p} = \vec{p}_{no\_RA} + \alpha\ \vec{p}_{add\_RA} + \tau(p_{t1}, p_{t2}, 0, 0, 0, 0) + \beta\ (0, 0, p_{f1}, p_{f2}, 0, 0)$$

where $\tau$ and $\beta$ are sigmoidal functions that track, respectively, how much time has passed and how many cells occupy the pFP attractor. Specifically,

$$\begin{aligned} \tau =& 0.5(\tanh(T_1 * (t - T_2)) + 1) \\ \beta =& 0.5 * (\tanh(b_1 * (\#pFP - b_2)) + 1) \end{aligned}$$

with $t$ the time point of the simulation, $T_1, T_2, b_1, b_2$ parameters to be fit and $\#pFP$ the number of points in a fixed region around the pFP attractor.

We then fit the model with the 21 parameters to predict the proportions of cell types. We use the results of the initial fitting to inform the priors taken for this second fitting.

Table A4: Tables of priors for the parameters of the second fitting.

| **Parameter** | **Prior** | **Parameters** | **Prior** |
|---|---|---|---|
| $p_1^{no\_RA}$ | $\mathcal{N}(-0.5, 0.3)$ | $p_1^{RA}$ | $\mathcal{N}(1.56, 0.17)$ |
| $p_2^{no\_RA}$ | $\mathcal{N}(0.7, 0.25)$ | $p_2^{RA}$ | $\mathcal{N}(0.64, 0.01)$ |
| $p_3^{no\_RA}$ | $\mathcal{U}([-1, 0])$ | $p_3^{RA}$ | $\mathcal{N}(-0.02, 0.24)$ |
| $p_4^{no\_RA}$ | $\mathcal{N}(-1.4, 0.1)$ | $p_4^{RA}$ | $\mathcal{N}(-0.5, 0.27)$ |
| $p_5^{no\_RA}$ | $\mathcal{N}(0.54, 0.2)$ | $p_5^{RA}$ | $\mathcal{N}(0.3, 0.06)$ |
| $p_6^{no\_RA}$ | $\mathcal{N}(0.5, 0.16)$ | $p_6^{RA}$ | $\mathcal{N}(0.6, 0.2)$ |
| $p_{t1}$ | $\mathcal{U}([-1, 2])$ | $p_{f1}$ | $\mathcal{U}([0, 1])$ |
| $p_{t2}$ | $\mathcal{U}([0.5, 2.5])$ | $p_{f2}$ | $\mathcal{U}([-2, -1])$ |
| $T_1$ | $\mathcal{U}([5, 10])$ | $b_1$ | $\mathcal{U}([5, 10])$ |
| $T_2$ | $\mathcal{U}([10, 20])$ | $b_2$ | $\mathcal{U}([0.1, 0.9])$ |
| $\sigma$ | $\mathcal{U}([0.012, 0.002])$ | | |

The results of the fitting algorithm after 7 rounds of fitting are shown in Figures A9, A10, A11 and A12.

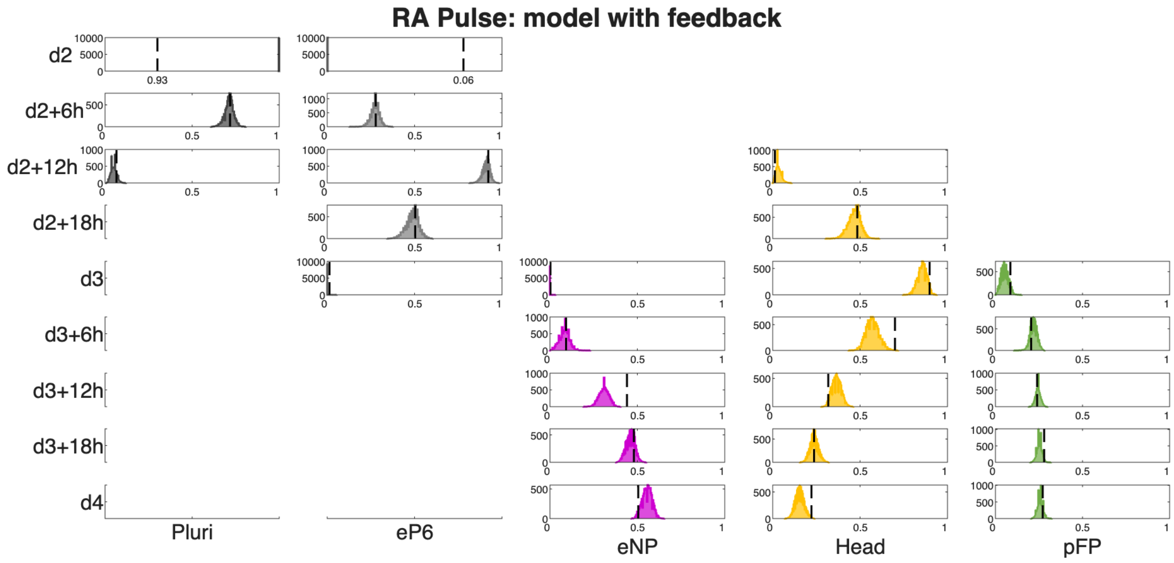

Figure A9: Histograms showing the distributions of cell states using the 10000 parameter vectors accepted in the final iteration of the fitting for the refined model including feedback. The vertical dotted lines indicate the proportions from the experimental series used as training data. The missing panels correspond to populations with no assigned simulated cells and experimental proportion equal to zero. Related to Fig. 3J.

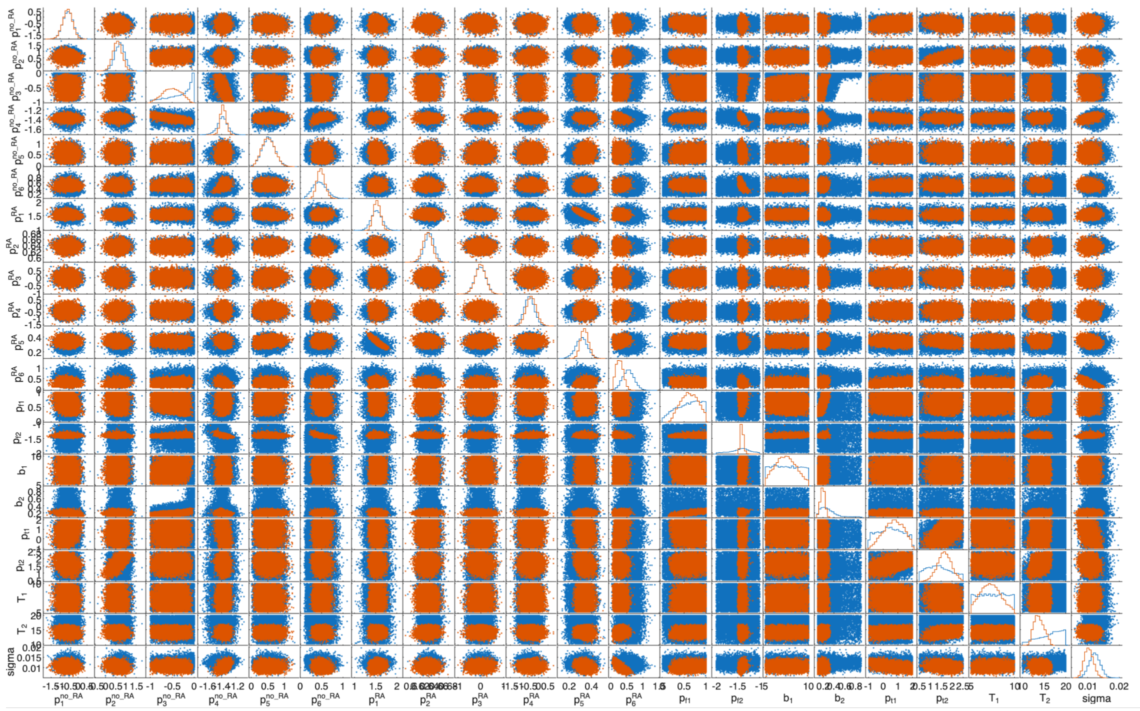

Figure A10: Distributions of accepted parameters in the first iteration (blue) compared to accepted parameters in the last iteration (red) of the fitting for the refined model including feedback.

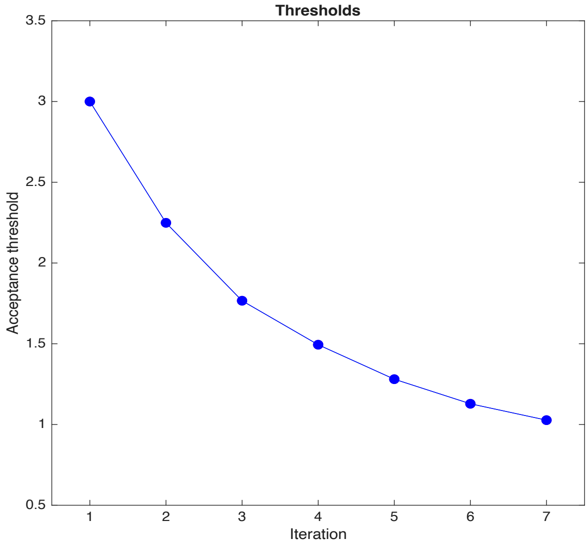

Figure A11: Evolution of acceptance threshold in the 7 iterations of the fitting for the refined model including feedback

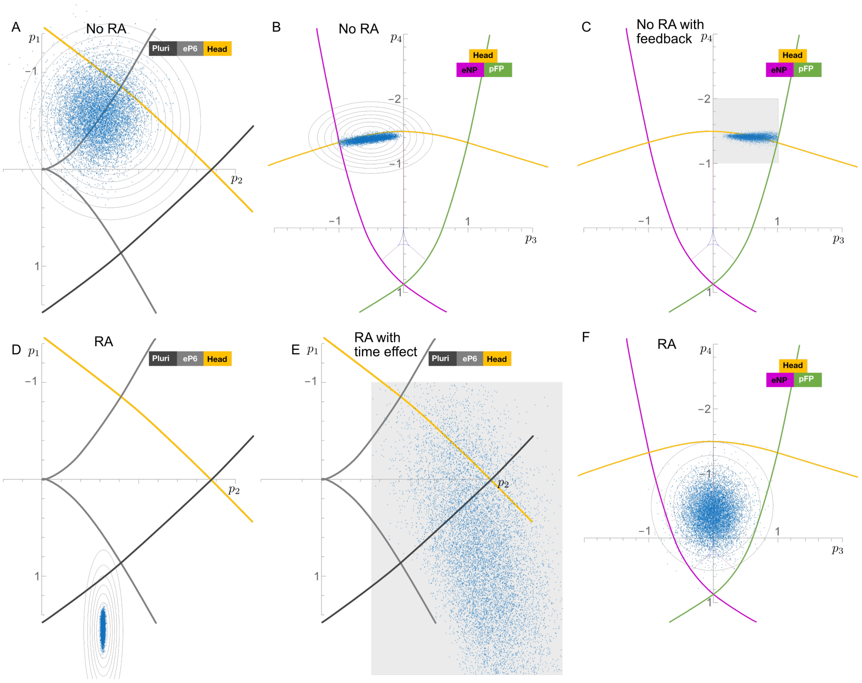

Figure A12: Priors (shaded area or gray level curves) and accepted parameters in the last round of fitting for the the refined model including feedback together with the corresponding bifurcation locus.

### 5 Predictions

After fitting the model to experimental data, we used it to predict cellular behavior in response to different perturbations. Each prediction involved specific modifications to the model characteristics while maintaining the overall landscape structure.

#### 5.1 Knockout of *FoxA2* (*WT:KO* chimeras)

To simulate *FoxA2-/-* cells, we fixed parameter $p_4$ to force the unstable manifold from the Head saddle toward the pNP attractor. For chimeric NTOs containing a defined proportion of knockout cells, we simulated the corresponding fraction of cells with the modified unstable manifold. All other model parameters remained unchanged. Figures A13-A22 show the predicted distributions of cell states across the 10000 parameter vectors accepted in the final iteration of the second fitting (Fig. A10) through the range of chimeras from 10% knockout cells to 100%. These predictions correspond to data presented in Fig. 4B-C.

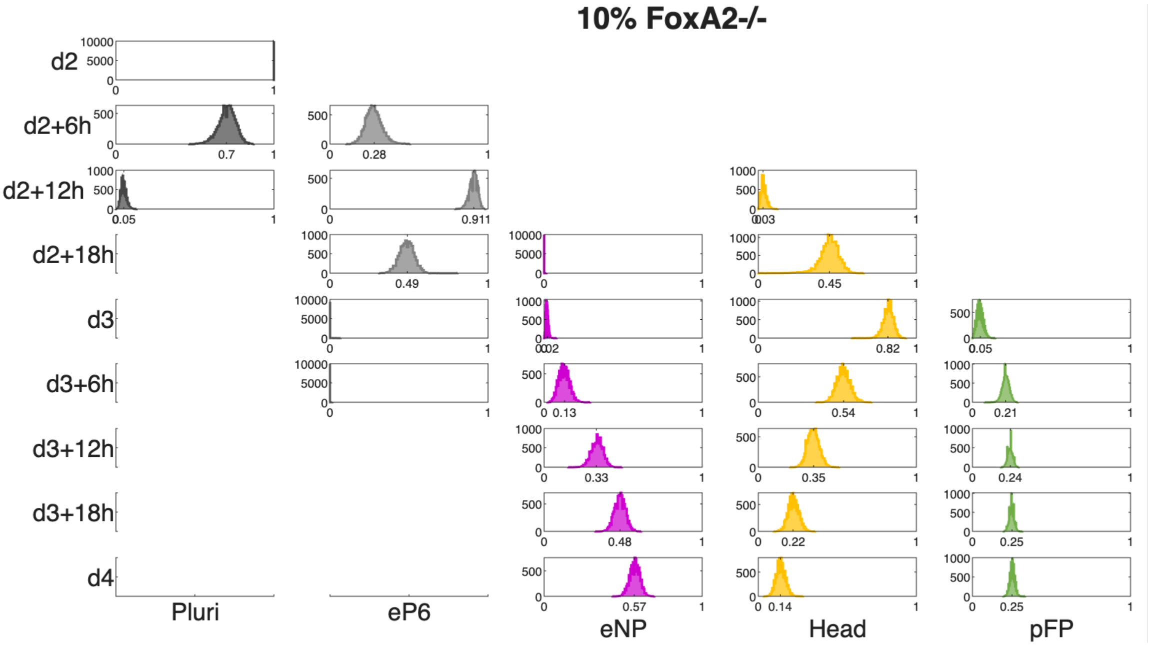

Figure A13: Histograms showing the distributions of cell states for chimera predictions with 10% knockout cells using 10000 accepted parameter vectors from the second fitting. Missing panels correspond to populations with no assigned simulated cells. Related to Fig. 4B-C.

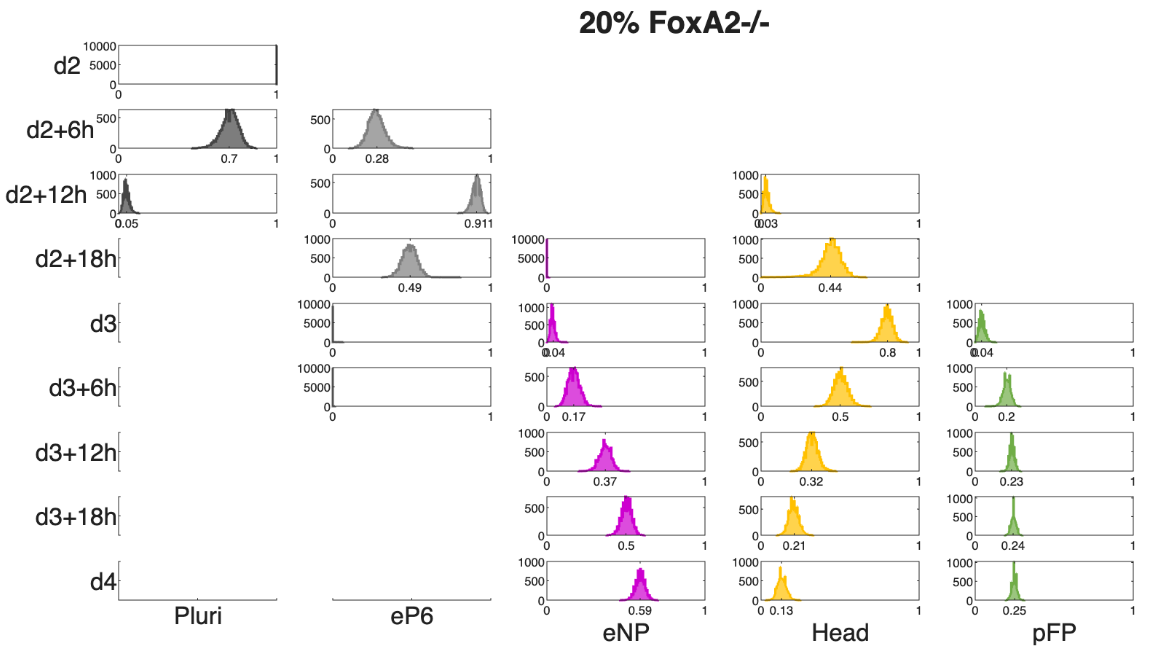

Figure A14: Histograms showing the distributions of cell states for chimera predictions with 20% knockout cells using 10000 accepted parameter vectors from the second fitting. Missing panels correspond to populations with no assigned simulated cells. Related to Fig. 4B-C.

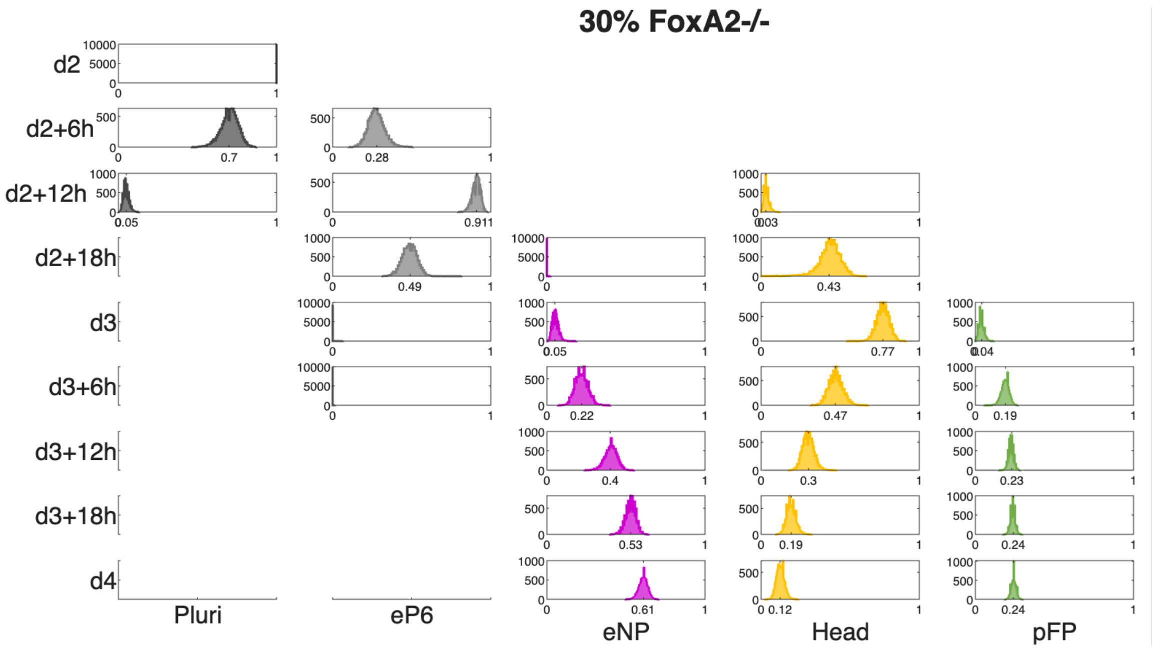

Figure A15: Histograms showing the distributions of cell states for chimera predictions with 30% knockout cells using 10000 accepted parameter vectors from the second fitting. Missing panels correspond to populations with no assigned simulated cells. Related to Fig. 4B-C.

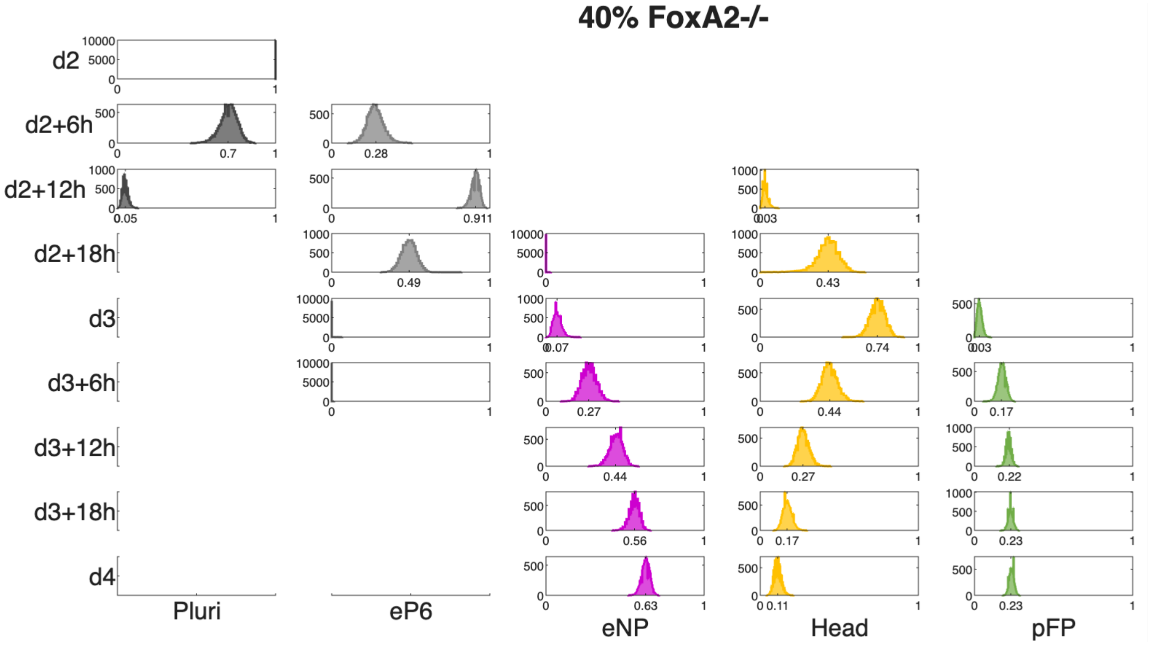

Figure A16: Histograms showing the distributions of cell states for chimera predictions with 40% knockout cells using 10000 accepted parameter vectors from the second fitting. Missing panels correspond to populations with no assigned simulated cells. Related to Fig. 4B-C.

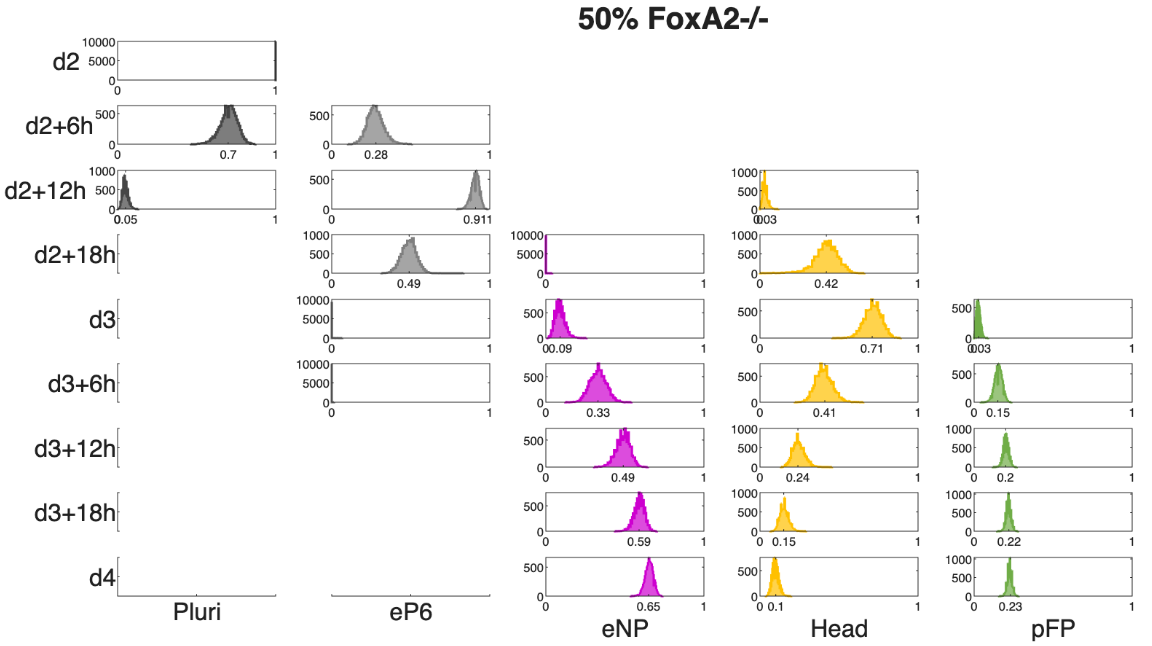

Figure A17: Histograms showing the distributions of cell states for chimera predictions with 50% knockout cells using 10000 accepted parameter vectors from the second fitting. Missing panels correspond to populations with no assigned simulated cells. Related to Fig. 4B-C.

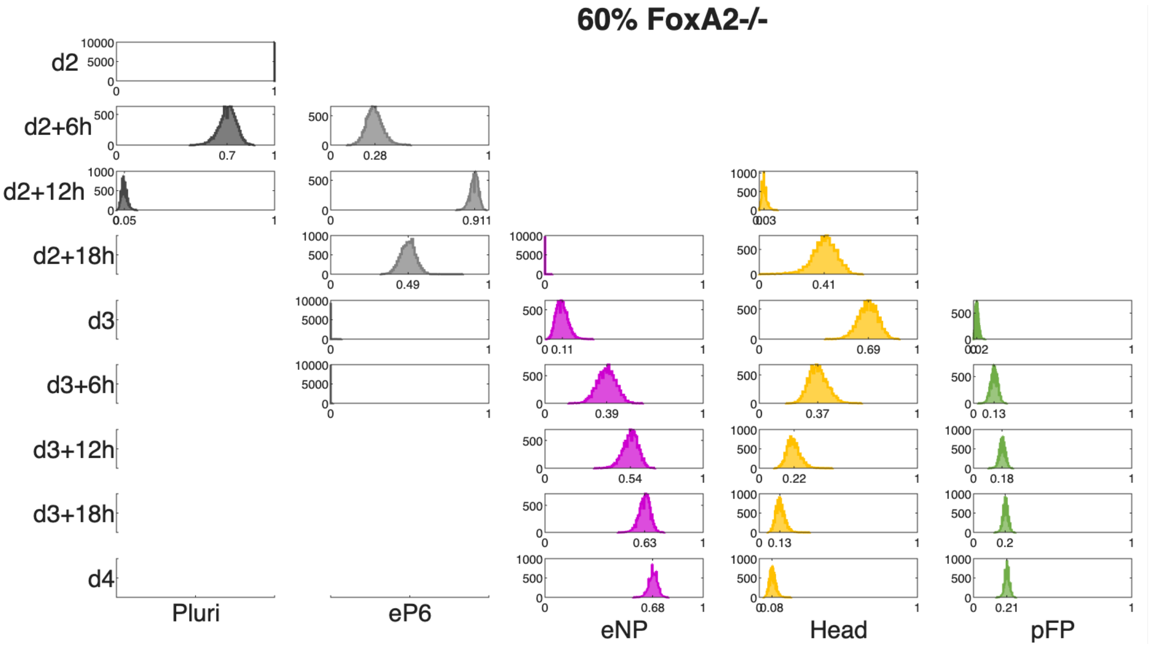

Figure A18: Histograms showing the distributions of cell states for chimera predictions with 60% knockout cells using 10000 accepted parameter vectors from the second fitting. Missing panels correspond to populations with no assigned simulated cells. Related to Fig. 4B-C.

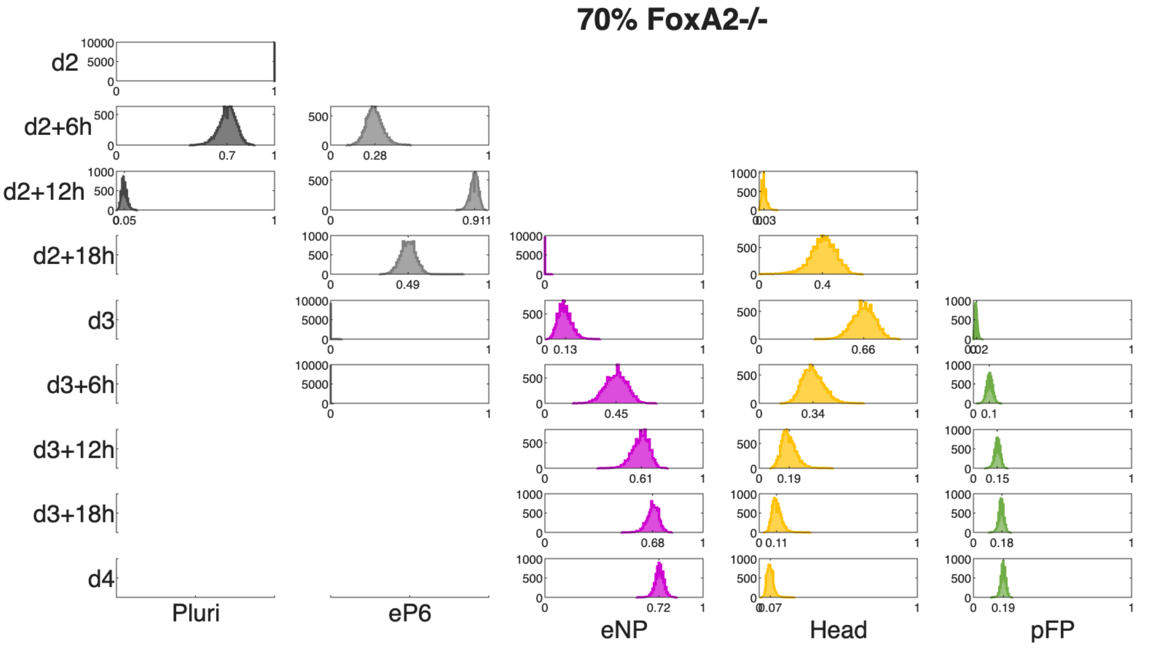

Figure A19: Histograms showing the distributions of cell states for chimera predictions with 70% knockout cells using 10000 accepted parameter vectors from the second fitting. Missing panels correspond to populations with no assigned simulated cells. Related to Fig. 4B-C.

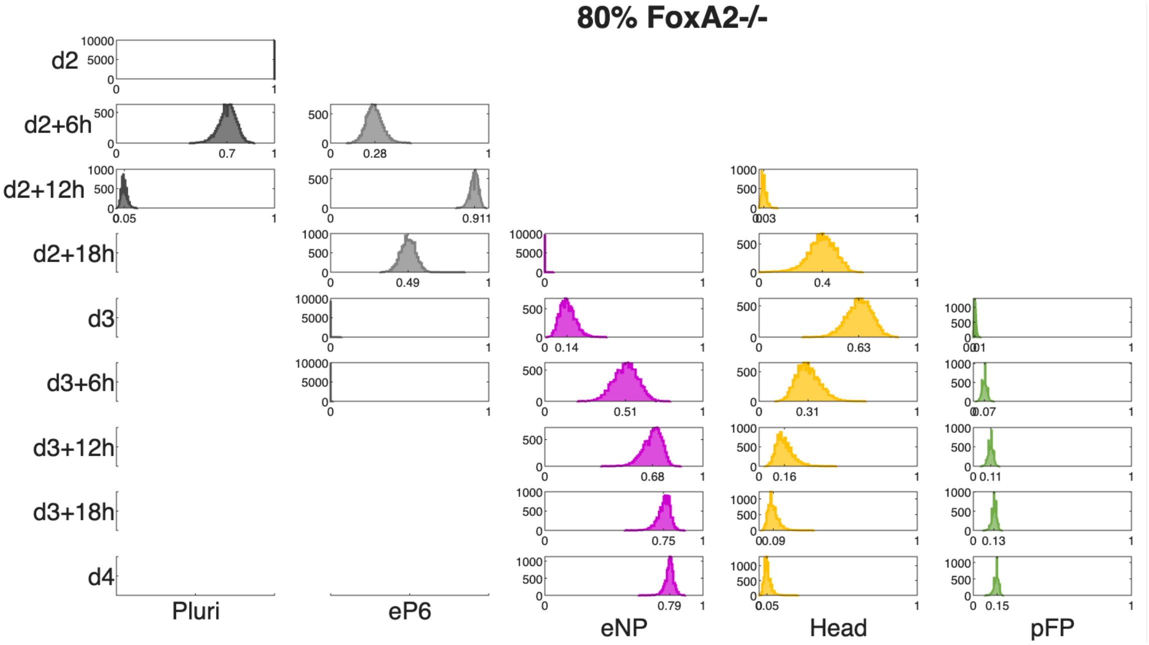

Figure A20: Histograms showing the distributions of cell states for chimera predictions with 80% knockout cells using 10000 accepted parameter vectors from the second fitting. Missing panels correspond to populations with no assigned simulated cells. Related to Fig. 4B-C.

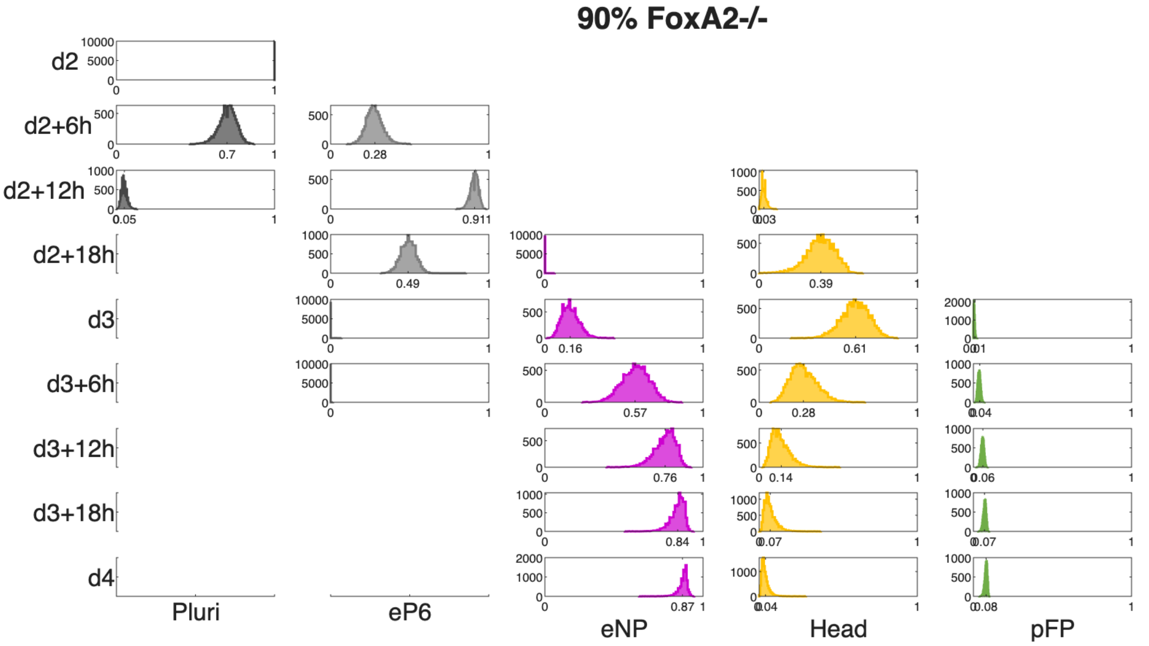

Figure A21: Histograms showing the distributions of cell states for chimera predictions with 90% knockout cells using 10000 accepted parameter vectors from the second fitting. Missing panels correspond to populations with no assigned simulated cells. Related to Fig. 4B-C.

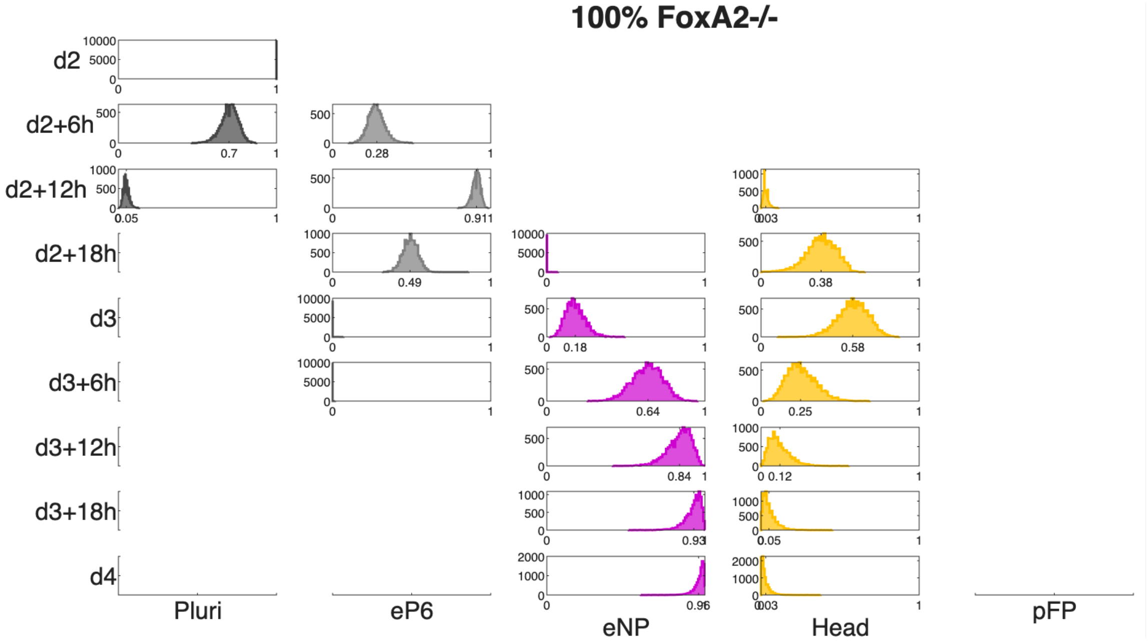

Figure A22: Histograms showing the distributions of cell states for chimera predictions with 100% knockout cells using 10000 accepted parameter vectors from the second fitting. Missing panels correspond to populations with no assigned simulated cells. Related to Fig. 4B-C.

#### 5.2 Feedback inhibition

We simulated regulative feedback inhibition by setting $\beta$ to zero from d2+12h to d3+18h. All other model parameters remained unchanged. Figure A23 shows the predicted distributions of cell states for the 10000 parameter vectors from the second fitting (Fig. A10). These predictions correspond to data in Fig. 4F.

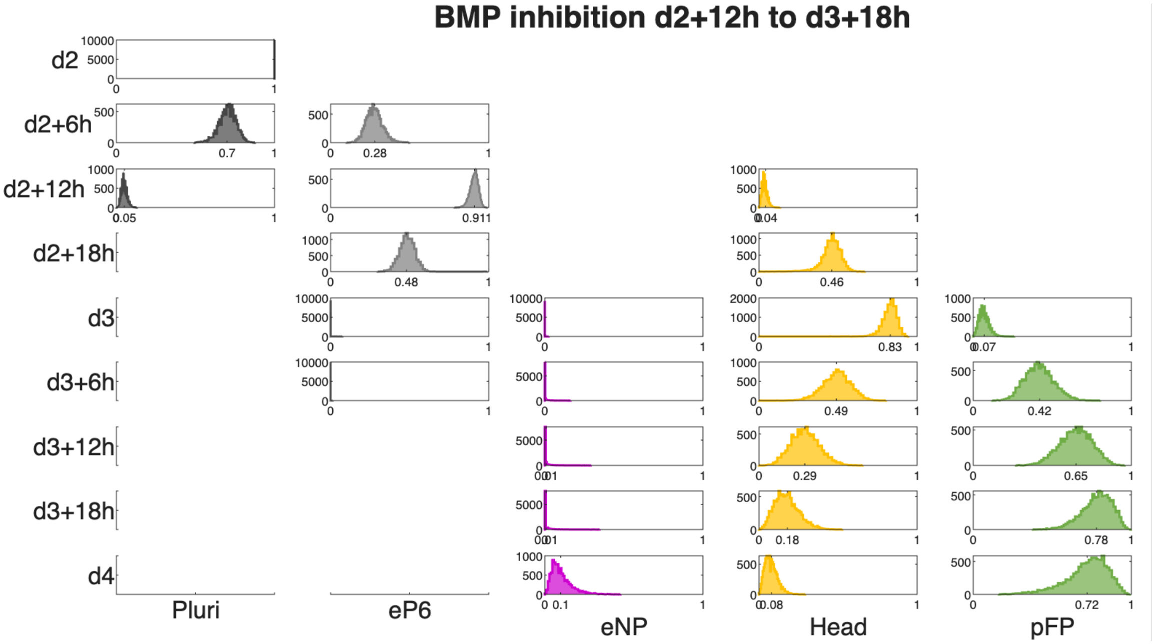

Figure A23: Histograms showing the distributions of cell states for BMP inhibition predictions using 10000 accepted parameter vectors from the fitting of the model including feedback. Missing panels correspond to populations with no assigned simulated cells. Parameters match those in Figure A10, except $\beta$ is set to zero. Related to Fig. 4F.

For feedback inhibition experiments using LDN, PD, or IWP2 treatments, we pooled data from all replicates for both control and treated conditions. We performed separate clustering analyses for each treatment following the methodology described in Section 2. Each treatment yielded three clusters with protein expression profiles consistent with Head, eNP, and pFP identities. Results are shown in Figures A24 and A25 and table A5 for LDN, Figure A26 and A27 and table A6 for PD, and Figure A28 and A29 and table A7 for IWP2 treatments.

Control

| | eNP | Head | pFP |
|---|---|---|---|
| r1 | 40 | 29 | 31 |
| r2 | 39 | 28 | 33 |
| r3 | 43 | 28 | 29 |

LDN

| | eNP | Head | pFP |
|---|---|---|---|
| r1 | 10 | 25 | 65 |
| r2 | 10 | 24 | 66 |
| r3 | 9 | 24 | 67 |

Table A5: Proportions obtained by clustering the RA treated samples as control (left) and the RA+LDN treated samples (right). All data correspond to d4, each row corresponds to a different replicate of the experiment. Related to Fig. 4G.

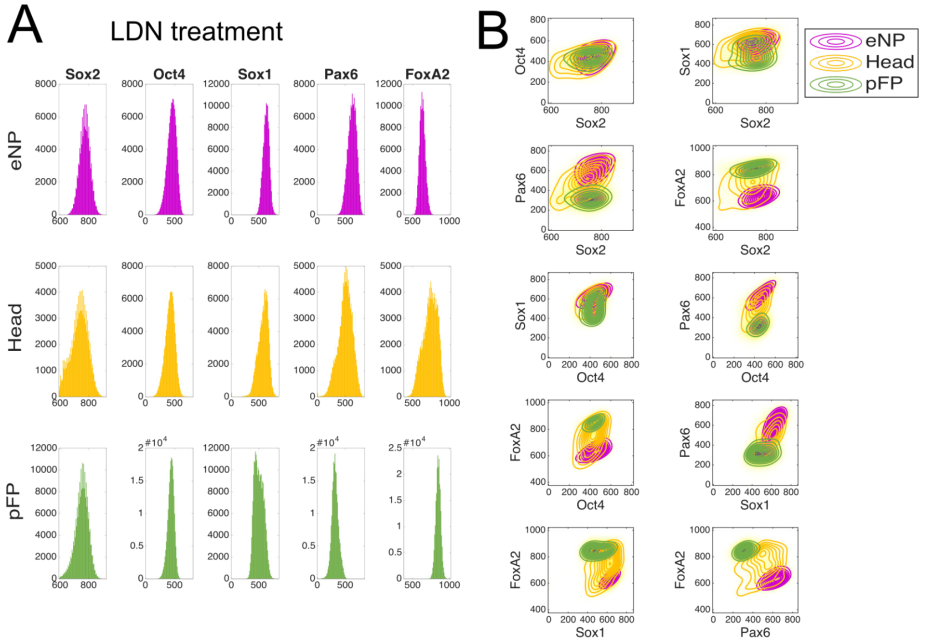

Figure A24: **Clustering results for LDN-treated cells.** A. Histograms showing one-dimensional distributions for each protein in each cluster, with specified cell identities. B. Two-dimensional distributions of protein expression for each cluster, with colors corresponding to panel A. Related to Fig. 4G.

Control

| | eNP | Head | pFP |
|---|---|---|---|
| r1 | 40 | 32 | 28 |
| r2 | 39 | 34 | 27 |
| r3 | 43 | 30 | 27 |

PD

| | eNP | Head | pFP |
|---|---|---|---|
| r1 | 43 | 33 | 24 |
| r2 | 41 | 35 | 24 |
| r3 | 40 | 36 | 25 |

Table A6: Proportions obtained by clustering the RA treated samples as control (left) and the RA+PD treated samples (right). All data correspond to d4, each row corresponds to a different replicate of the experiment. Related to Fig. 4G.

Control

| | eNP | Head | pFP |
|---|---|---|---|
| r1 | 41 | 30 | 28 |
| r2 | 40 | 32 | 28 |
| r3 | 44 | 28 | 27 |

IWP2

| | eNP | Head | pFP |
|---|---|---|---|
| r1 | 36 | 33 | 31 |
| r2 | 36 | 32 | 32 |
| r3 | 33 | 36 | 31 |

Table A7: Proportions obtained by clustering the RA treated samples as control (left) and the RA+IWP2 treated samples (right). All data correspond to d4, each row corresponds to a different replicate of the experiment. Related to Fig. 4G.

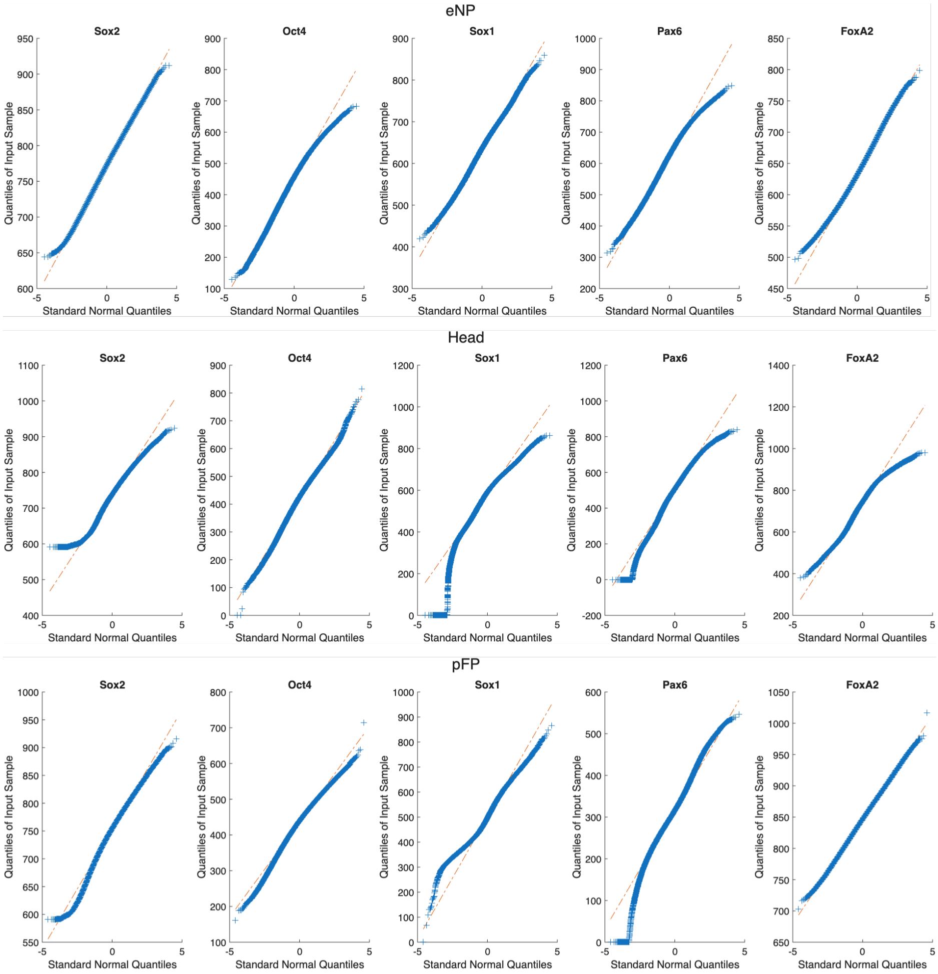

Figure A25: Q-q plots for the clusters obtained by Gaussian mixture model clustering of the dataset including LDN treated samples.

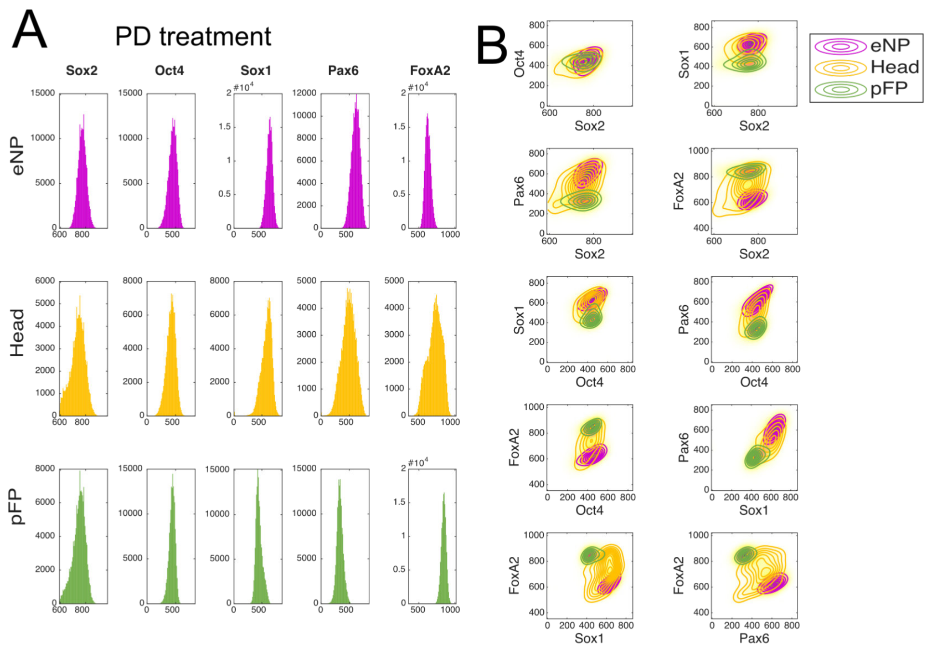

Figure A26: **Clustering results for PD-treated cells.** A. Histograms showing one-dimensional distributions for each protein in each cluster, with specified cell identities. B. Two-dimensional distributions of protein expression for each cluster, with colors corresponding to panel A. Related to Fig. 4G.

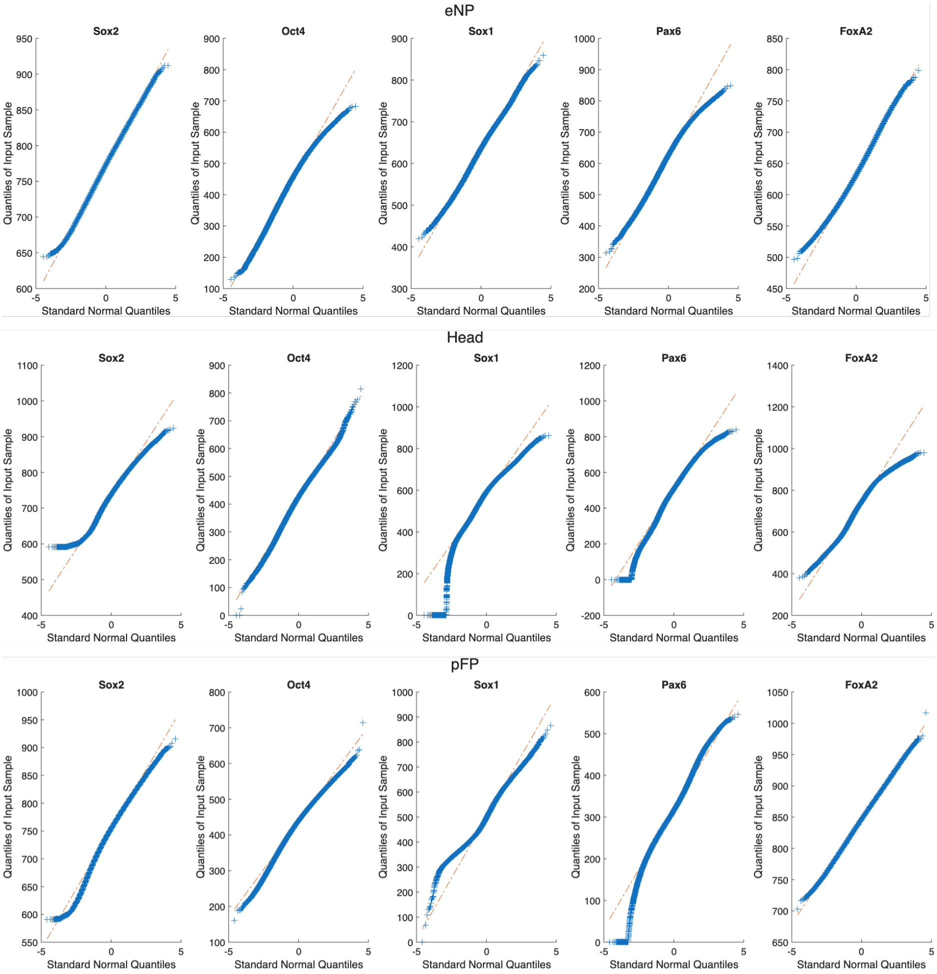

Figure A27: Q-q plots for the clusters obtained by Gaussian mixture model clustering of the dataset including PD treated samples.

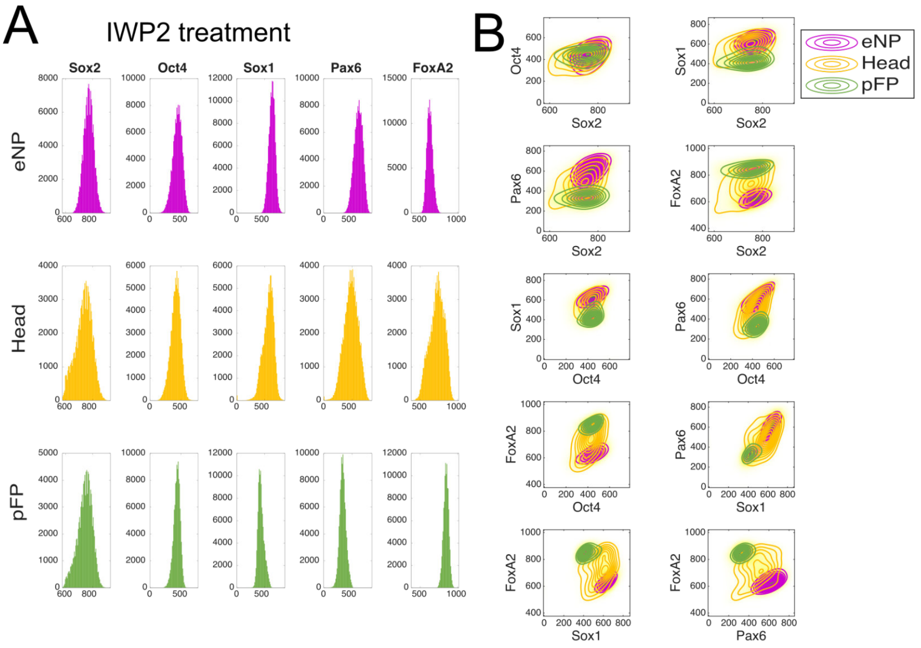

Figure A28: **Clustering results for IWP2-treated cells.** A. Histograms showing one-dimensional distributions for each protein in each cluster, with specified cell identities. B. Two-dimensional distributions of protein expression for each cluster, with colors corresponding to panel A. Related to Fig. 4G.

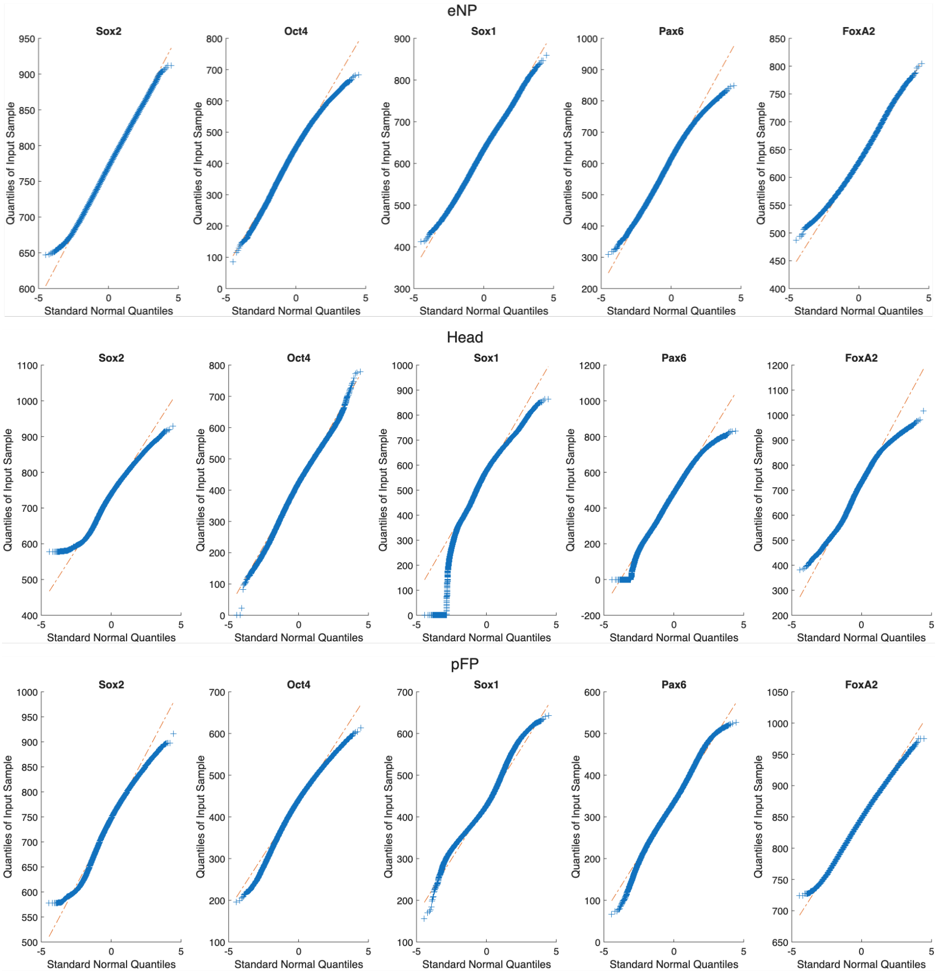

Figure A29: Q-q plots for the clusters obtained by Gaussian mixture model clustering of the dataset including IWP2 treated samples.

#### 5.3 RA withdrawal time

We simulated different RA withdrawal times by modifying the time course of parameter $\alpha$, while keeping all other model parameters unchanged. Figure A30 shows the predicted distributions of cell states across the simulated time course for the 10000 parameter vectors from the second fitting (Fig. A10). These predictions correspond to data in Fig. S5A.

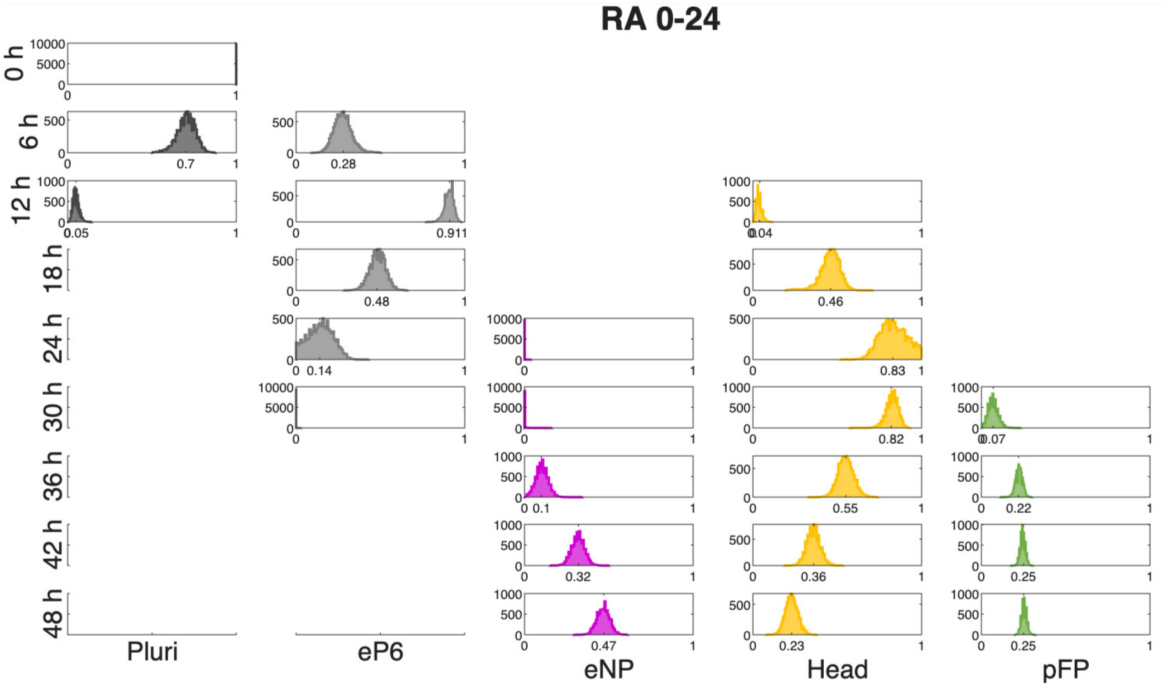

Figure A30: Histograms showing the distributions of cell states for the prediction of the experiment with RA withdrawal after 24 hours using the 10000 parameter vectors accepted in the final iteration of the second fitting. The missing panels correspond to populations with no assigned simulated cells. Related to Fig. S5A.

#### 5.4 Head start

To simulate the $'2xTetOn'$ system, in which FoxA2 and Pax6 are co-induced by doxycycline, we initialized cells directly at the Head attractor. Although double positive cells could correspond to cells already in transition from the Head attractor, since in the simulation cells quickly start to transition from the attractor, it is reasonable and convenient to start all cells in the attractor. This simplification has minimal effect on the results. Since the precise Head attractor position varies with specific parameter values, we used a standard position within the Head region as the initial condition and allowed the dynamics to evolve from there. Figure A31 shows the predicted distributions of cell states across the simulated time course for the 10000 parameter vectors from the second fitting (Fig. A10). These predictions correspond to data in Fig. 5B.

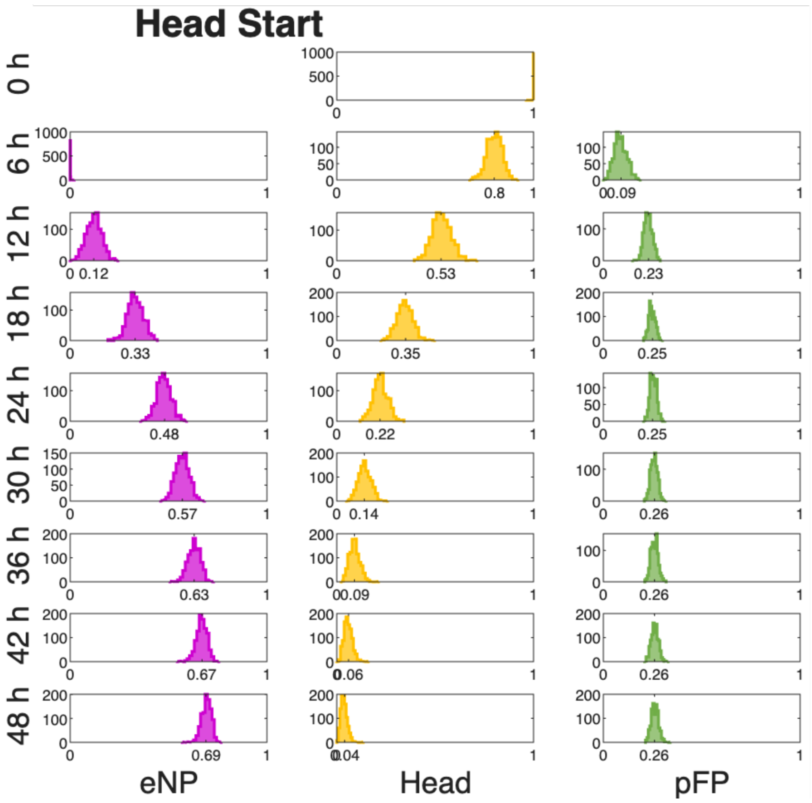

Figure A31: Histograms showing the distributions of cell states for Head start predictions using 10000 accepted parameter vectors from the second fitting algorithm. Missing panels correspond to populations with no assigned simulated cells. Related to Fig. 5B.
